# Ethanolic Extract of Polish Propolis exhibits synergy with selected antifungal agents against yeast pathogens causing candidiasis

**DOI:** 10.64898/2026.04.21.719917

**Authors:** Piotr Bollin, Michał K. Pierański, Piotr Marek Kuś, Patrick Van Dijck, Piotr Szweda

## Abstract

Candidiasis pose a serious health threat, stimulating efforts to develop new antifungal agents and alternative therapies. Given the high mortality of fungal infections and the historical use of natural remedies, there is a growing interest in integrating natural substances into modern treatments. It is particularly important to explore interactions between home remedies and clinically approved antifungals to avoid harmful combinations or enhance beneficial effects. In this study, the chemical composition of the ethanolic extract of propolis (EEP) using UHPLC-DAD-QqTOF-MS was analyzed. The interactions of this extract with several antifungal agents against four yeast pathogens causing candidiasis: *Candida albicans, Nakaseomyces glabratus, Pichia kudriavzevii*, and *Candida auris* were investigated using Checkerboard Titration Assay, Growth Kinetics, and Disc-diffusion assay. Also, a novel simulated infection model was proposed. The results showed synergistic interactions between EEP and amphotericin B, and additive effects with nystatin. Synergy and additivity with fluconazole and voriconazole were observed, but limited to *C. albicans* and *N. glabratus*. In contrast, antagonistic interactions were noted with caspofungin, clotrimazole, and ketoconazole, which may have clinical relevance. Additionally, positive interactions with 2-phenoxyethanol and silver nanoparticles (AgNPs) suggest potential practical applications. Propolis’s synergistic properties could expand antifungal strategies and support the development of multi-target, resistance-preventing therapies.

## 1. Introduction

In recent decades, there has been an increase in the incidence of candidiasis. Candidiasis has long been described as an infection caused by *Candida* spp.^1^ However, recent taxonomic revisions have reclassified several members of this genus into other genera. Accordingly, *Candida glabrata* is referred to in this article as *Nakaseomyces glabratus* (although some authors use the name *N. glabrata*)^2^, and *Candida krusei* as *Pichia kudriavzevii*^3^. There are also proposals to reassign *Candida auris* to the genus *Candidozyma*^4^; however, this reclassification has not yet been consistently confirmed across all clades and multiple strains and remains a matter of ongoing debate^5^. Although the prevalence of *C. albicans* is decreasing in favour of other yeast causing candidiasis, it is still responsible for over 45% of the infections. The second most common cause is *N. glabratus*, which, together with *C. albicans*, accounts for over 60% of infections in Europe. *P. kudriavzevii* is the most common representative among the rare isolates, placing it fifth in terms of the number of patients suffering from candidiasis^6^. Although not as prevalent, it warrants significant attention due to its intrinsic resistance to fluconazole and rapid development of resistance to other antifungals^7^. Even though *C. auris* is the most recently identified member of the *Candida* genus, it is under intense surveillance because nearly 40% of isolates are multidrug-resistant, and it spreads exponentially following outbreaks^8,9,10^. It is estimated that each year, fungal infections contribute to the deaths of approximately 1.5 million people. Beyond drug resistance, a significant issue is the limited arsenal of antifungal drugs, comprising only five classes: polyenes, pyrimidine analogues, azoles, allylamines, and echinocandins, coupled with the very slow introduction of new drugs^11,12^. This is why many researchers are focusing on proposing alternative therapies, including the use of natural products such as propolis. Propolis is a complex resinous material synthesized by bees through the amalgamation of plant-derived resins, beeswax, and salivary secretions^13^. Depending on the botanical source and environmental conditions, its chemical composition is highly variable, yielding a wide range of bioactive components, mainly flavonoids, phenolic acids and their derivatives, terpenoids, aromatic and essential oils, as well as other phenolic compounds including stilbenes, lignans, and coumarins^14,15^. This compositional diversity underpins the extensive pharmacological potential of propolis extracts, which encompasses antimicrobial, anti-inflammatory, antioxidant, and antineoplastic activities, positioning it as a promising agent for diverse biomedical and therapeutic applications^16,17^. The antifungal activity of propolis has already been demonstrated in multiple *in vitro* studies^18,19^, as well as in clinical studies, for example, against oral candidiasis caused by *C. albicans*^20^. Due to high mortality rates and increasing drug resistance, clinicians attempt to overcome these obstacles during patient care by utilizing the phenomenon of synergy. For example, the Infectious Diseases Society of America recommends a combination of amphotericin B and 5-fluorocytosine, and cites case reports of successful treatment with 5-fluorocytosine combined with fluconazole^21^. Clinical success has also been reported with combinations of amphotericin B with fluconazole or caspofungin, caspofungin with 5-fluorocytosine, caspofungin with voriconazole, or fluconazole with micafungin^22^. Positive interactions of propolis extracts have been evidenced with anticancer drugs, metformin, as well as with antimicrobial agents such as antibiotics or antifungal drugs. Unfortunately, interactions between propolis and antifungals have been tested for only a few agents, and most studies employed only a single method of synergy testing^23^. Multiple reports indicate the necessity of testing synergy using more than one method, preferably employing three different groups of tests, such as disc-diffusion assays, checkerboard, and time-kill assays^24,25,26,27,28^. Therefore, in this work, the characterisation of the composition of the Polish ethanolic extract of propolis (EEP) and examination of the interactions of EEP with a variety of clinically approved antifungals and the most common disinfectants were performed. Three different methods for most combinations were utilized, and for azoles and amphotericin B, a new method of synergy testing - the simulated infection model - was proposed. The research was conducted using two of the most common pathogens causing candidiasis: *C. albicans* and *N. glabratus*, and two exhibiting a high level of drug resistance: *P. kudriavzevii* and *C. auris*. The obtained data encompass a wide range of combinations, allowing for comparisons between the drugs and among the genera.

## 2. Results

### 2.1. Determination of Chemical Composition of propolis extracts by UHPLC-DAD-QqTOF-MS

A total of 77 compounds were identified in the investigated propolis extract using UHPLC-DAD-QqTOF-MS, and additionally, their abundance was estimated based on MS ion count and supported by available literature data (Supplementary materials - Table S1). Several of the compounds were quantified by UHPLC-DAD (Table 1). The chromatogram obtained at 280 nm is presented in Figure 1. The propolis contained mainly a variety of flavonoids, including, among others, pinocembrin, pinostrobin, sakuranetin (flavanones), pinobanksin (flavanonol), apigenin, chrysin (flavones), galangin, kampferol (flavonols), 2’,6’-dihydroxy-4’-methoxydihydrochalcone (chalcone), different phenylpropanoids (*p*-coumaric acid, caffeic acid, ferulic acid, isoferulic acid, and their esters). The major part of the flavonoids was present in the form of aglycones, some were present as esters (including the major compound - pinobanksin acetate), and interestingly, only one was identified as a glucoside (apigenin-7-*O*-glucoside). The identification of compounds was based on exact mass, MS^2^ fragmentation, UV spectra, or comparison with MS^2^ databases (Bruker MetaboBASE Personal Library 3.0, MS Dial library (MSMS-Public-VS15), and LC-MS/MS MoNA library) as well as reference compounds. Phenylpropanoid derivatives identity was confirmed based on observation of characteristic mass losses and diagnostic ions (caffeoyl: 179/161; *p*-coumaroyl: 163/145; feruloyl: 193/175) in MS^2^ spectra of their esters. The chemical profile of the investigated propolis corresponds to the characteristics of the orange subgroup of the European poplar type propolis that contains higher amounts of pinocembrin, galangin, and chrysin than the blue subgroup^29^. This also corresponds to the phytochemical profile of black poplar bud exudates that contain elevated amounts of these compounds and other components such as methyl-butenyl caffeates^30^. Nevertheless, propolis investigated in this study also contained glycerides (2-Acetyl-1,3-di-p-coumaroylglycerol, 2-Acetyl-3-p-coumaroyl-1-feruloylglycerol, 2-Acetyl-1,3-di-feruloylglycerol) characteristic of aspen bud exudates^30,31^, which suggests a mixed botanical origin with possible predominance of black poplar. The major compounds previously identified as relevant as potential markers related to the high antifungal activity of propolis extracts^32^ were quantified using UHPLC-DAD (Table 1). The most abundant compounds were pinocembrin (39.2 mg/g), chrysin (36.2 mg/g), followed by galangin (27.1 mg/g), pinobanksin acetate (22.7 mg/g), and pinobanksin (13.2 mg/g). These amounts are comparable to the highest values reported in our previous research (up to 45.7, 27.5, 28.6, 38.1, 12.22 mg/g, respectively). The mixture of equal parts of these compounds was found to exhibit elevated fungistatic and fungicidal potential due to synergistic effects^32^.

**Table 1.** The content of major compounds relevant as potential markers related to high antifungal activity of propolis extracts, determined by UHPLC-DAD (data expressed as mg/g of propolis extract).

| No. | Compound | RT<br>[min] | Content<br>[mg/g] |
| --- | --- | --- | --- |
| 1 | Pinobanksin | 26.613 | 13.2 |
| 2 | Caffeic acid methylbutenyl ester isomer <sup>a</sup> | 31.42 | 9.1 |
| 3 | Caffeic acid methylbutenyl ester isomer <sup>a</sup> | 32.15 | 8.1 |
| 4 | Chrysin | 33.23 | 36.2 |
| 5 | Pinocembrin | 34.16 | 39.2 |
| 6 | Isosakuranetin | 34.51 | 5.3 |
| 7 | Galangin | 34.82 | 27.1 |
| 8 | Pinobanksin acetate <sup>c</sup> | 35.97 | 22.7 |
| 9 | *p*-Coumaric acid methylbutenyl ester <sup>b</sup> | 37.81 | 2.3 |
| 10 | 2'6'-Dihydroxy-4'-methoxydihydrochalcone | 41.15 | 1.6 |
| 11 | *p*-Coumaric acid cinnamyl ester <sup>b</sup> | 41.91 | 4.4 |
| 12 | Techtochrysin | 42.2 | 6.3 |
| 13 | Pinostrobin | 42.39 | 8.0 |

<sup>a</sup> calculated as caffeic acid; <sup>b</sup> calculated as *p*-coumaric acid; <sup>c</sup> calculated as pinobanksin; equivalents were corrected considering molecular weight differences;

**Figure 1.**
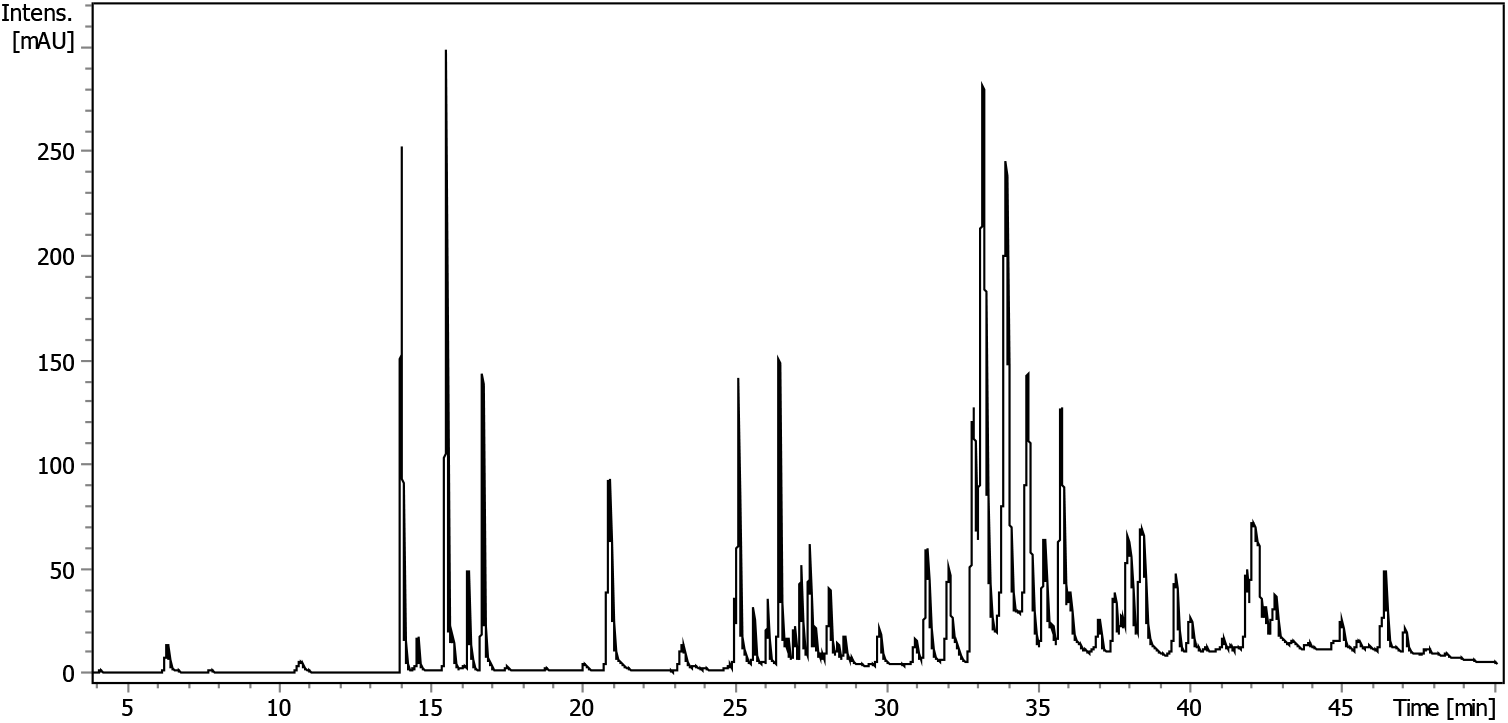
Chromatogram of the investigated propolis sample recorded at 280 nm.

### 2.2. Determination of MIC and MFC values

To evaluate the antifungal activity of ethanolic extract of propolis (EEP) in combination with conventional antifungal agents, the minimum inhibitory concentration (MIC) and minimum fungicidal concentration (MFC) were determined for four yeasts causing candidiasis. The results are summarized in Table 2. The susceptibility profiles for polyene antifungals were consistent across all tested species. MIC values for amphotericin B ranged from 0.5 to 2 µg/mL, for nystatin, this value equalled to 4 µg/mL, and natamycin exhibited MICs between 2 and 4 µg/mL. MFCs were generally equivalent to MICs or up to two-fold higher, except for *N. glabratus*, for which the MFC of amphotericin B was four times greater than the MIC. In contrast, susceptibility to echinocandins displayed greater variability. MICs for caspofungin ranged from 0.125 to 1 µg/mL, with the lowest susceptibility exhibited by *P. kudriavzevii*. Anidulafungin and micafungin demonstrated significantly lower MICs, ranging from 0.008 to 0.25 µg/mL, except in the case of *C. auris*, where the MIC for anidulafungin reached 1.563 µg/mL. MFCs for echinocandins were typically equal to or twice the MIC; however, *C. auris* showed markedly higher MFCs, with micafungin and caspofungin MFCs being 50- and 255-fold higher than the corresponding MICs, respectively. Due to the primarily fungistatic nature of azoles, MFCs could not be determined for this class. Fluconazole exhibited higher MICs compared to other azoles, with values of 0.125 µg/mL for *C. albicans* and 2 µg/mL for *C. auris*. In contrast, voriconazole, clotrimazole, and ketoconazole demonstrated MICs in the range of 0.004 to 0.063 µg/mL. Notably, *N. glabratus* and *P. kudriavzevii* required 32 and 64 µg/mL of fluconazole, respectively, whereas the MICs for the other azoles ranged from 0.125 to 4 µg/mL. The antifungal agent 5-fluorocytosine showed MICs between 0.031 and 0.063 µg/mL for most of the yeast species, except for *P. kudriavzevii* (12.5 µg/mL). MFCs values in this case were typically 2-to 4-fold higher than the values of MICs. For ciclopirox, both MIC and MFC values ranged between 0.25 and 0.5 µg/mL, although *C. auris* required a concentration of 2 µg/mL. Terbinafine exhibited activity only against *C. albicans* (MIC = 6.25 µg/mL). Chlorhexidine inhibited the growth at values ranging from 1.563 µg/mL (*P. kudriavzevii*) to 12.5 µg/mL (*C. auris*), with MFCs equal to MICs across all tested species. Octenidine dihydrochloride demonstrated MIC values from 0.195 to 0.781 µg/mL; MFCs were equal to or two-fold higher than the MICs. MICs for 2-phenoxyethanol ranged from 0.313% to 0.625%. While MFCs were equal to the MIC for *N. glabratus*, they were 4-to 8-fold higher for the remaining tested species. Hydrogen peroxide displayed relatively low MICs, ranging from 0.003% (*P. kudriavzevii*) to 0.006% (*C. albicans*), with corresponding MFCs four times higher. In contrast, *N. glabratus* and *C. auris* had MICs of 0.023%, with MFCs elevated 40-to 80-fold, respectively. MIC estimated for AgNPs ranged from 0.234 to 0.938 µg/mL, with MFCs two-to four-fold higher. Among the tested species, *C. albicans* turned out to be the least susceptible to AgNPs. Finally, EEP demonstrated MICs ranging from 32 to 128 µg/mL. MFCs were generally equivalent to or twice the MIC, except for *N. glabratus*, where the MFC was eight times higher than the MIC.

**Table 2.** Minimum inhibitory concentrations (MIC_90_) and minimum fungicidal concentrations (MFC) of different agents determined against selected yeast pathogens causing candidiasis. Numbers with a greater-than symbol (>) indicate values outside of the analysed concentration range. All values presented were rounded to three decimal places.

| Group of agents | Agent | MIC and MFC (µg/mL or %) |  |  |  |  |  |  |  |
| --- | --- | --- | --- | --- | --- | --- | --- | --- | --- |
|  |  | *Candida albicans*<br>SC 5314 |  | *Nakaseomyces glabratus*<br>DSM 11226 |  | *Pichia kudriavzevii*<br>DSM 6128 |  | *Candida auris*<br>B 8441 |  |
|  |  | MIC | MFC | MIC | MFC | MIC | MFC | MIC | MFC |
| Polyenes | Amphotericin B | 0.5 | 1 | 1 | 4 | 2 | 2 | 1 | 1 |
|  | Nystatin | 4 | 4 | 4 | 4 | 4 | 4 | 4 | 8 |
|  | Natamycin | 4 | 4 | 2 | 4 | 2 | 4 | 4 | 8 |
| Echinocandins | Caspofungin | 0.125 | 0.125 | 0.25 | 0.5 | 1 | 2 | 0.391 | 100 |
|  | Anidulafungin | 0.016 | 0.031 | 0.063 | 0.125 | 0.125 | 0.25 | 1.563 | 1.563 |
|  | Micafungin | 0.016 | 0.031 | 0.008 | 0.016 | 0.25 | 0.25 | 0.391 | 12.5 |
| Azoles | Fluconazole | 0.125 | >128 | 32 | >128 | 64 | >128 | 2 | >128 |
|  | Voriconazole | 0.004 | >64 | 2 | >64 | 0.5 | >64 | 0.031 | >64 |
|  | Clotrimazole | 0.016 | >64 | 4 | >64 | 0.125 | >64 | 0.016 | >64 |
|  | Ketoconazole | 0.008 | >64 | 4 | >64 | 1 | >64 | 0.063 | >64 |
| Pyrimidine analogues | 5-Fluorocytosine | 0.063 | 0.25 | 0.031 | 0.125 | 12.5 | 25 | 0.063 | 0.125 |
| N-hydroxypyridones | Ciclopirox | 0.5 | 0.5 | 0.25 | 0.5 | 0.5 | 0.5 | 2 | 2 |
| Allylamines | Terbinafine | 6.25 | >100 | >100 | >100 | >100 | >100 | >100 | >100 |
| Disinfectants | Chlorhexidine | 6.25 | 6.25 | 3.125 | 3.125 | 1.563 | 1.563 | 12.5 | 12.5 |
|  | Octenidine dihydrochloride | 0.195 | 0.391 | 0.391 | 0.391 | 0.391 | 0.391 | 0.781 | 0.781 |
|  | 2-Phenoxyethanol | 0.313% | 2.5% | 0.625% | 0.625% | 0.313% | 1.25% | 0.313% | 1.25% |
|  | Hydrogen peroxide | 0.006% | 0.023% | 0.023% | 0.938% | 0.003% | 0.012% | 0.023% | 1.875% |
| Nanoparticles | AgNPs | 0.938 | 3.75 | 0.938 | 1.875 | 0.234 | 0.469 | 0.469 | 0.938 |
| Extracts | Ethanollic extract of propolis (EEP) | 128 | 128 | 32 | 256 | 64 | 128 | 64 | 128 |

### 2.3. Assessment of Interactions Between EEP and Antifungal Agents via Checkerboard Titration Assay

The checkerboard titration assay revealed that amphotericin B exhibited synergistic activity with the ethanolic extract of propolis (EEP) across all tested pathogens (Table 3). Detailed MIC values for each compound, both alone and in combination, along with the corresponding fractional inhibitory concentration (FIC) indices, are provided in Supplementary Tables S2–S19. To facilitate interpretation, isobolograms were constructed for each interaction. Figure 2 presents isobolograms for fluconazole, illustrating characteristic curve shapes corresponding to indifference, additivity, and antagonism. Isobolograms for all additional compound combinations are available in Supplementary Figures S1– S17. In the case of polyenes, the combinations of EEP with nystatin and natamycin predominantly resulted in additive effects, except for *C. auris*, which demonstrated an indifferent response. Among echinocandins, combinations of EEP with anidulafungin and micafungin were largely indifferent, except for *C. auris*, where an additive interaction was observed. In contrast, the combination of caspofungin and EEP resulted in strong antagonism against *C. albicans* and *N. glabratus*, while the remaining strains exhibited indifferent responses. 5-Fluorocytosine displayed additive effects when combined with EEP against *N. glabratus* and *P. kudriavzevii*, whereas interactions were indifferent for *C. albicans* and *C. auris*. A similar trend was noted for ciclopirox, with indifference observed only for *C. auris*. For azole antifungals, fluconazole and voriconazole displayed additive interactions with EEP for *N. glabratus*, antagonistic effects for *C. auris*, and indifference for two other tested species. In contrast, clotrimazole and ketoconazole showed strong antagonism for *C. albicans* and *C. auris*, while interactions were indifferent in the remaining species. Terbinafine exhibited activity exclusively against *C. albicans*, where its combination with EEP resulted in an additive effect. Hydrogen peroxide also showed additive interactions with EEP, except for *N. glabratus*, where synergy was observed. Synergistic interactions were detected for combinations of EEP with chlorhexidine for *N. glabratus* and *C. auris*, while additivity and indifference were noted for *C. albicans* and *P. kudriavzevii*, respectively. The combination of EEP with octenidine dihydrochloride yielded antagonistic effects for *C. albicans* and *C. auris*, and indifference in the remaining species. Conversely, EEP combined with 2-phenoxyethanol resulted in additive effects for all tested species except *P. kudriavzevii*. Finally, the combination of EEP with AgNPs demonstrated consistent synergistic activity across all tested yeasts.

**Table 3.** Summary of the ∑FIC values and antifungal interaction types determined for selected agents against yeast pathogens causing candidiasis using the Checkerboard Titration method. Detailed MIC and FIC values for each compound are provided in Supplementary Materials: Tables S2-S19.

|  | *Candida albicans* |  | *Nakaseomyces glabratus* |  | *Pichia kudriavzevii* |  | *Candida auris* |  |
| --- | --- | --- | --- | --- | --- | --- | --- | --- |
|  | FIC | Int. type | FIC | Int. type | FIC | Int. type | FIC | Int. type |
| Amphotericin B | 0.375 | S | 0.5 | S | 0.5 | S | 0.5 | S |
| Nystatin | 1 | Add | 1 | Add | 0.75 | Add | 1.031 | I |
| Natamycin | 1 | Add | 0.75 | Add | 1 | Add | 1.031 | I |
| Caspofungin | >4.063 | Ant | 4.5 | Ant | 1.031 | I | 2.031 | I |
| Anidulafungin | 2.25 | I | 1.031 | I | 1.031 | I | 0.625 | Add |
| Micafungin | 2.031 | I | 1.031 | I | 1.031 | I | 1 | Add |
| 5-Fluorocytosine | 1.031 | I | 1 | Add | 0.75 | Add | 1.031 | I |
| Ciclopirox | 0.625 | Add | 1 | Add | 0.75 | Add | 1.031 | I |
| Fluconazole | 1.031 | I | 0.75 | Add | 1.031 | I | >4.03<br>1 | Ant |
| Voriconazole | 1.031 | I | 0.75 | Add | 1.031 | I | 4.25 | Ant |
| Ketoconazole | >4.031 | Ant | 1.031 | I | 1.031 | I | >4.06<br>3 | Ant |
| Clotrimazole | >4.031 | Ant | 1.031 | I | 1.031 | I | >4.06<br>3 | Ant |
| Terbinafine | 0.625 | Add | n/a | n/a | n/a | n/a | n/a | n/a |
| Hydrogen peroxide | 0.625 | Add | 0.313 | S | 0.625 | Add | 1 | Add |
| Chlorhexidine | 0.531 | Add | 0.5 | S | 1.031 | I | 0.5 | S |
| Octenidine dihydrochloride | >4.125 | Ant | 2.125 | I | 2.125 | I | >4.03<br>1 | Ant |
| 2-Phenoxyethanol | 0.75 | Add | 0.625 | Add | 1.031 | I | 1 | Add |
| AgNPs | 0.5 | S | 0.313 | S | 0.375 | S | 0.313 | S |

Int. type – interaction type; (S) – synergy; (Add) – additivity; (I) – indifference; (Ant) – antagonism; n/a – not analyzed

**Figure 2.**
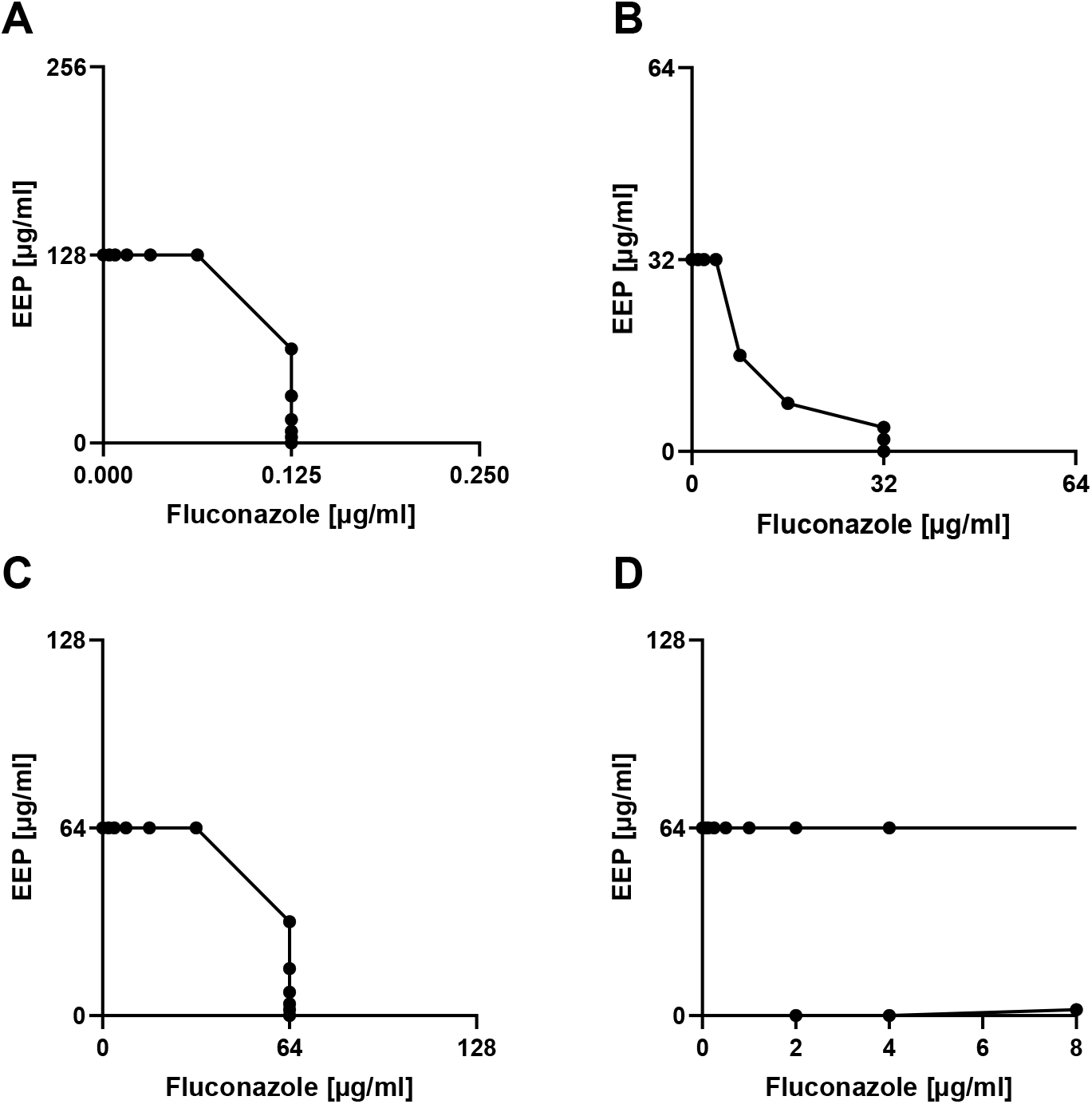
Isobolograms presenting the dependence between EEP and fluconazole concentrations regarding the MIC_90_ determined against A – *C. albicans;* B – *N. glabratus;* C – *P. kudriavzevii;* D – *C. auris*.

### 2.4. Growth kinetics analysis

Growth inhibition was assessed for the most important representatives of each group, for which the most promising results were observed. Figure 3 presents representative growth curves for four yeast species exposed to fluconazole, EEP, and their combinations, enabling dynamic assessment of antifungal activity over time. Growth inhibition curves for all remaining drug-EEP interactions are available in Supplementary Figures S18–S30. Interactions were interpreted based on standardized fractional MIC combinations and are presented in Table 4. Synergy was defined when 0.25×MIC of each agent in combination produced greater inhibition than 0.5×MIC of either agent alone, additivity was assigned when the combination of 0.5×MIC + 0.5×MIC was more effective than individual treatments, indifference was defined when the combination yielded effects equivalent to the single-agent 0.5×MIC treatment, and antagonism was recorded when the combination inhibited growth less effectively than either compound alone at 0.5×MIC. As in previous assays, amphotericin B consistently demonstrated synergy with EEP across all tested species. Nystatin similarly showed uniform additive effects. Caspofungin exhibited antagonism against most species, except for *P. kudriavzevii*, where an indifferent response was observed. Fluconazole displayed synergy with EEP for *C. albicans* and *N. glabratus*, indifference for *C. auris*, and antagonism for *P. kudriavzevii*. Voriconazole produced synergistic effects for *C. albicans*, antagonism for *C. auris*, and additive effects in the remaining species. Both clotrimazole and ketoconazole resulted in additive interactions for *C. albicans*, antagonism for *N. glabratus*, and either indifference or additivity in *P. kudriavzevii*, as well as antagonism and indifference for *C. auris*, respectively. The combination of EEP with 5-fluorocytosine produced additive effects across all tested pathogens. Chlorhexidine exhibited additive effects for *C. albicans* and *P. kudriavzevii*, and synergistic activity for *N. glabratus* and *C. auris*. For 2-phenoxyethanol, synergy was observed for *C. albicans*, while additive effects were seen in other species. AgNPs showed universal synergy across all tested yeasts. Hydrogen peroxide exhibited synergy for *C. albicans* and *N. glabratus*, and additivity for *P. kudriavzevii* and *C. auris*. Lastly, ciclopirox displayed additive interactions for *C. albicans* and *C. auris*, and synergy for *N. glabratus* and *P. kudriavzevii*.

**Table 4.** Interaction outcomes between EEP and antifungal agents based on growth curve analysis. Interactions were evaluated by comparing the inhibitory effects of compound combinations at fractional MICs. Synergy was defined when 0.25×MIC of each agent in combination produced a stronger inhibitory effect than either agent alone at 0.5×MIC. Additivity was assigned when the combination of 0.5×MIC + 0.5×MIC was more effective than either single treatment. Indifference was determined when the combination produced an effect equivalent to 0.5×MIC of either agent alone. Antagonism was concluded when the combined treatment resulted in weaker inhibition than one or both agents at 0.5×MIC.

|  | *Candida albicans* | *Nakaseomyces glabratus* | *Pichia kudriavzevii* | *Candida auris* |
| --- | --- | --- | --- | --- |
| Amphotericin B | S | S | S | S |
| Nystatin | Add | Add | Add | Add |
| Caspofungin | Ant | Ant | I | Ant |
| Fluconazole | S | S | Ant | I |
| Voriconazole | S | Add | Add | Ant |
| Clotrimazole | Add | Ant | I | Ant |
| Ketoconazole | Add | Ant | Add | I |
| 5-Fluorocytosine | Add | Add | Add | Add |
| Chlorhexidine | Add | S | Add | S |
| 2-Phenoxyethanol | S | Add | Add | Add |
| AgNPs | S | S | S | S |
| Hydrogen Peroxide | S | S | Add | Add |
| Ciclopirox | Add | S | S | Add |

(S) – synergy; (Add) – additivity; (I) – indifference; (Ant) – antagonism

**Figure 3.**
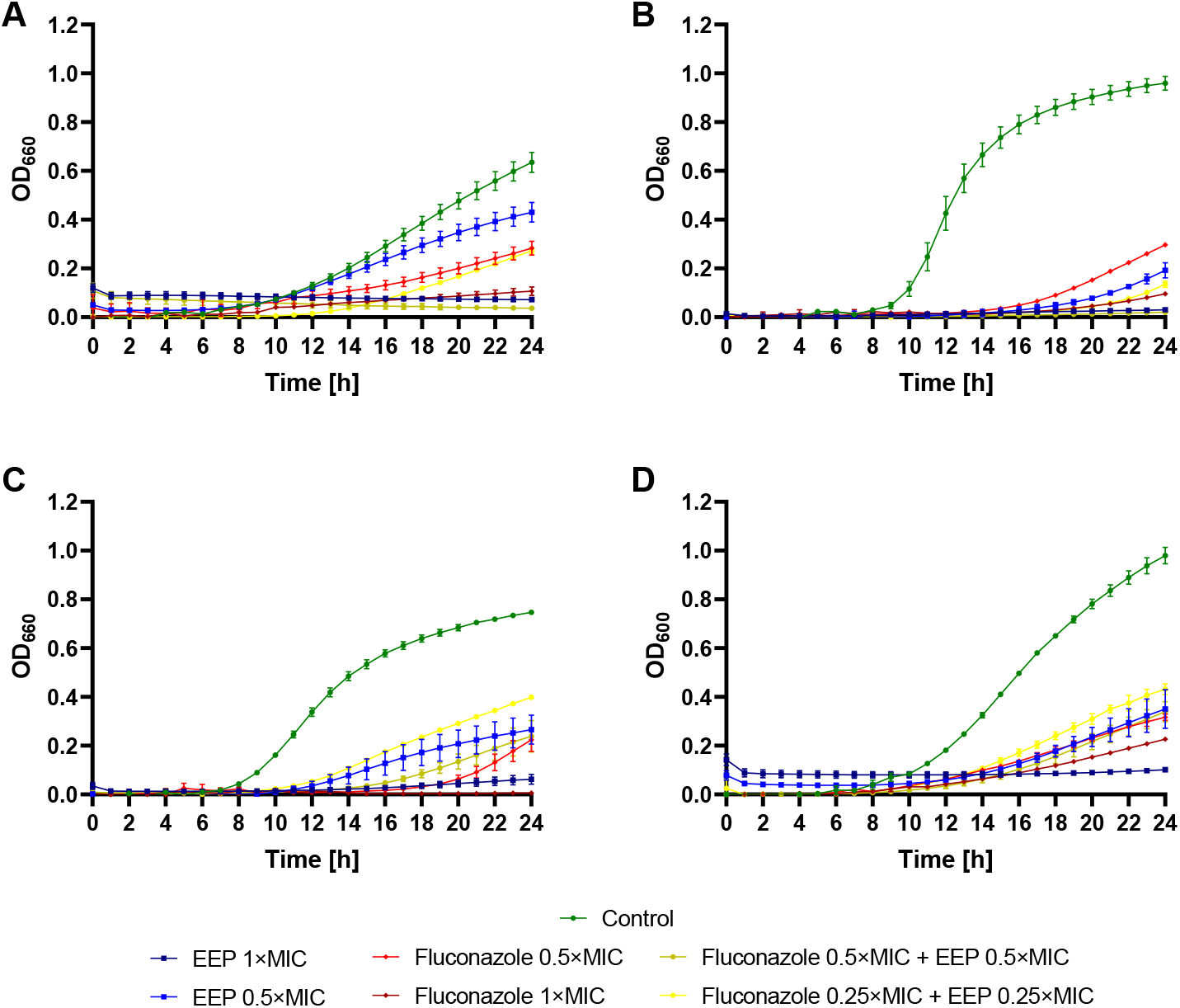
Antifungal activity of EEP, fluconazole, and their mixtures – growth curves (monitoring of the growth kinetics of the yeast cells measured as turbidity of the cultures at wavelength 660 nm (or 600 nm for *C. auris*) against A – *C. albicans*; B – *N. glabratus*; C – *P. kudriavzevii*; D – *C. auris*.

### 2.5. Disc diffusion assay

In the disc diffusion assay, propolis was applied to paper discs containing antifungal agents, while 70% ethanol was used as a control to account for solvent effects. An increase in the inhibition zone greater than 2 mm was interpreted as a synergistic interaction. An increase between 1 and 2 mm indicated additivity, while changes within ±1 mm were considered indicative of indifference. A decrease greater than 1 mm was classified as antagonism. The differences are summarized in Table 5, while inhibition zones diameters are available in supplementary data in Table S21. Synergy between amphotericin B and EEP was observed against *C. albicans* and *N. glabratus*, whereas additivity was noted for the remaining species. Similarly, nystatin showed synergy against *C. albicans* and *N. glabratus*, but exhibited indifference for *P. kudriavzevii* and antagonism for *C. auris*. The combination of caspofungin with EEP resulted in antagonism across all strains except *P. kudriavzevii*, where additivity was observed. All four tested azoles exhibited synergy with EEP against *C. albicans*. For *N. glabratus*, fluconazole and clotrimazole demonstrated synergy, while voriconazole showed indifference and ketoconazole antagonism. For *P. kudriavzevii*, combinations of EEP with fluconazole and ketoconazole resulted in antagonism, whereas the remaining azoles showed indifference. Notably, for *C. auris*, EEP exhibited synergy with fluconazole and voriconazole, but antagonism with clotrimazole and ketoconazole. For combinations of EEP with 5-fluorocytosine, synergy was observed against all tested species except *C. auris*, where an indifferent effect was noted. In contrast, combinations with chlorhexidine generally resulted in indifferent interaction, except for antagonism observed against *P. kudriavzevii*. The combination of EEP with 2-phenoxyethanol showed synergy for *C. albicans* and *N. glabratus*, additivity for *P. kudriavzevii*, and indifference for *C. auris*. AgNPs combined with EEP exhibited synergy for *C. auris*, antagonism for *N. glabratus*, and additivity for other yeasts. EEP combined with hydrogen peroxide resulted in synergy against *P. kudriavzevii* and additivity for the other pathogens. Finally, the combination of EEP with ciclopirox exhibited an indifferent effect against all tested species, except for *C. auris*.

**Table 5.** Mean inhibition zones difference (in mm) between control and EEP treatment and antifungal interactions estimated with disc diffusion assay against selected pathogens causing candidiasis.

|  | *Candida albicans* |  | *Nakaseomyces glabratus* |  | *Pichia kudriavzevii* |  | *Candida auris* |  |
| --- | --- | --- | --- | --- | --- | --- | --- | --- |
| Amphotericin B | 6.3 | S | 3.5 | S | 1.7 | Add | 1.7 | Add |
| Nystatin | 4.0 | S | 5.0 | S | -0.5 | I | -1.8 | Ant |
| Caspofungin | -1.7 | Ant | -1.3 | Ant | 2.0 | Add | -7.5 | Ant |
| Fluconazole | 10.0 | S | 3.0 | S | -2.3 | Ant | 2.2 | S |
| Voriconazole | 14.0 | S | 0.3 | I | 0.7 | I | 2.7 | S |
| Clotrimazole | 3.3 | S | 8.3 | S | 0.3 | I | -14.0 | Ant |
| Ketoconazole | 5.0 | S | -6.3 | Ant | -5.7 | Ant | -4.3 | Ant |
| 5-Fluorocytosine | 5.7 | S | 6.3 | S | 3.3 | S | 0.0 | I |
| Chlorhexidine | 0.2 | I | 0.0 | I | -2.2 | Ant | -0.8 | I |
| 2-Phenoxyethanol | 5.7 | S | 2.7 | S | 1.7 | Add | 0.3 | I |
| AgNPs | 1.3 | Add | -7.7 | Ant | 1.3 | Add | 6.0 | S |
| Hydrogen Peroxide | 1.3 | Add | 1.5 | Add | 2.3 | S | 1.2 | Add |
| Ciclopirox | -0.7 | I | -0.2 | I | -0.7 | I | -1.2 | Ant |

(S) – synergy; (Add) – additivity; (I) – indifference; (Ant) – antagonism

### 2.6. Simulated infection model

To evaluate the response of pathogens causing candidiasis to prolonged antifungal exposure and to further characterize interactions with azoles - the most commonly used agents in clinical antifungal therapy - a simulated infection model was developed by our team. In this model, yeast cultures were subjected to repeated treatment and regrowth cycles over five consecutive days. At the end of the experiment, colony-forming units were quantified to determine whether the combination of EEP with antifungal agents exerted a stronger inhibitory effect than the sum of CFU/mL reductions for both treatments alone. These differences are presented in Table 6 and Figures S31-34 in the supplementary materials, and the detailed data can be found in Table S22. Apart from azoles, amphotericin B, which previously demonstrated the most pronounced synergy or additivity in other assays, was also included for comparative analysis. In this prolonged exposure model, amphotericin B retained its synergistic interaction with EEP across all tested species, except for *C. auris*, where an additive effect was observed. Fluconazole, in combination with EEP, demonstrated synergy against *C. albicans* and *N. glabratus*. However, this combination showed indifference for *C. auris* and antagonism for *P. kudriavzevii*. Combinations of EEP with ketoconazole and clotrimazole were predominantly antagonistic, except for an indifferent interaction with ketoconazole for *N. glabratus*, and with clotrimazole for *C. auris*.

**Table 6.** CFU reduction difference between the combination (drug with EEP) and the drug alone combined with EEP alone, in a simulated infection model after 5 days of treatment.

|  | *Candida albicans* |  | *Nakaseomyces glabratus* |  | *Pichia kudriavzevii* |  | *Candida auris* |  |
| --- | --- | --- | --- | --- | --- | --- | --- | --- |
| Amphotericin B | 4.50 | S | 0.54 | S | 1.54 | S | 0.31 | Add |
| Fluconazole | 3.76 | S | 0.59 | S | -1.77 | Ant | -0.47 | I |
| Ketoconazole | -1.29 | Ant | -0.35 | I | -4.71 | Ant | -0.89 | Ant |
| Clotrimazole | -0.87 | Ant | -3.08 | Ant | -3.54 | Ant | -0.16 | I |

Types of interaction: (S) – synergy; (Add) – additivity; (I) – indifference; (Ant) – antagonism; n/a – not analyzed. ChT – Checkerboard Titration; GK – Growth Kinetics; D-D – Disc-diffusion; SI – Simulated Infection; AMB – amphotericin B; NS – nystatin; CAS – caspofungin; 5-FC – 5-fluorocytosine; CPX – ciclopirox; FLU – fluconazole; VOR – voriconazole; CLO – clotrimazole; KCA – ketoconazole; H_2_O_2_ – hydrogen peroxide; CHX – chlorhexidine; 2-PhOEtOH – 2-phenoxyethanol; AgNPs – silver nanoparticles.

## 3. Discussion

Depending on the origin of the propolis and the extraction method, EEP exhibit variable antifungal and antimicrobial efficacy. In this study, the major compounds identified in EEP included pinocembrin, chrysin, galangin, pinobanksin acetate, pinobanksin, and caffeic acid. These findings are consistent with our previous studies^32,33,34,35^, as well as with other independent studies on EEP^14,19^. There are relatively few reports detailing MIC values for multiple antifungal agents across different species formerly recognized as the *Candida* genus. Although the compounds tested in Table 2 are not novel, many have not yet been evaluated against *C. auris*. Our comprehensive analysis enables a direct comparison of several antifungal agents tested under consistent conditions against various fungal pathogens. The MIC values obtained in this study are consistent with previously published data on amphotericin B, 5-fluorocytosine, echinocandins, azoles, chlorhexidine, and AgNPs^36,37,38,39,40,41^. MIC values for propolis align with our earlier reports^32^. To the best of our knowledge, EEP has not been tested against *C. auris* before, and only a single study has demonstrated the activity of individual phenolic components against this pathogen^42^. Thus, this is the first report showing that EEP can effectively eradicate *C. auris in vitro*. The summary of interactions between EEP and antifungal agents using four different methods is shown in Table 7. Amphotericin B combined with EEP demonstrated clear synergistic effects across all tested yeast species in at least three out of four assays. For *C. auris*, two methods indicated additivity rather than full synergy. These observations align with prior studies showing synergy between amphotericin B and Tunisian propolis in *Leishmania* spp.^43^. Synergistic effects with Brazilian propolis have been reported for *C. albicans* and *Trichophyton rubrum*, and additive effects for *C. parapsilosis*^44^. Additionally, a combination of propolis and carnosic acid has been shown to synergize with amphotericin B in the treatment of *N. glabratus*^45^. Regarding nystatin, our results indicate additive interactions with EEP for all tested pathogens, except for *C. auris*, for which the results were inconclusive. Other groups have previously reported synergistic effects of nystatin with red or Serbian propolis against *C. albicans* using checkerboard and disc-diffusion assays^46,47^. Similarly, synergy between propolis and natamycin has been described for *C. albicans*, whereas our data suggest rather an additive interaction^48^. Although polyenes have been in clinical use for over 60 years, their precise mechanisms of action remain partially unresolved^49,50^. Synergy between polyene macrolides and EEP may result from combined membrane-disrupting activity or from enhanced oxidative stress due to reactive oxygen species (ROS) accumulation induced by both agents^51,52,53,54,55^. To date, there have been no reports examining the interaction between propolis and caspofungin. Our results show clear antagonism in all species except for *P. kudriavzevii*, where the interaction is at best described as indifferent. Prior study has reported indifference between Brazilian red propolis and anidulafungin for *C. parapsilosis* and *C. tropicalis*^56^, which is consistent with our findings. Echinocandins inhibit β-1,3-D-glucan synthase, a key component of the fungal cell wall. Although echinocandins share a common target, caspofungin exhibits distinct pharmacokinetics and pharmacodynamics compared to micafungin and anidulafungin^57^. The observed antagonism may be attributed to the fact that caspofungin is susceptible to oxidation, and EEP induces ROS accumulation^55,58^. Pre-exposure of *C. albicans* to hydrogen peroxide has also been shown to increase tolerance to caspofungin, suggesting that oxidative stress can modulate susceptibility^59^. Our findings also support previously reported additive interactions between EEP and 5-fluorocytosine in *C. albicans*^48^. In our study, this additivity extended to all tested species, except for *C. auris*, for which an indifferent interaction was observed. The primary mechanism of 5-fluorocytosine is inhibition of RNA and DNA synthesis^60^. EEP induces various cellular stress responses, including oxidative damage, which may overwhelm fungal cells when nucleic acid synthesis is simultaneously blocked. Additionally, EEP-induced membrane disruption could facilitate increased uptake of 5-fluorocytosine, enhancing its antifungal activity^52,61^. There are no previous reports on the interaction between propolis and ciclopirox. Our results suggest an additive effect across all tested species, except for *C. auris*, where an indifferent interaction was observed. Ciclopirox acts by chelating metal ions, disrupting the activity of iron-dependent enzymes, impairing mitochondrial function, and compromising membrane integrity^62^. Given the multifaceted mechanisms of action of both ciclopirox and EEP, it is difficult to pinpoint which pathways are primarily responsible for their interaction. The combination of EEP with fluconazole showed synergistic effects for *C. albicans* and *N. glabratus*, but antagonistic or indifferent effects for *P. kudriavzevii* and *C. auris*. Earlier studies reported synergy with Brazilian red propolis in *C. parapsilosis* and *C. tropicalis*^56^. Other reports have shown additive effects with fluconazole in *C. albicans* and *C. parapsilosis*, and synergy in *T. rubrum*^44^. In this work, similar interaction profiles were observed with voriconazole, showing additivity for *C. albicans* and *N. glabratus*, but indifference or antagonism for *P. kudriavzevii* and *C. auris*. Our previous studies using the disc diffusion method also indicated synergy between EEP and both fluconazole and voriconazole for *C. albicans*^63^. For clotrimazole and ketoconazole, results varied considerably across methods, preventing a definitive conclusion for *C. albicans*. In order to address this issue, further studies using multiple strains and methodologies are necessary. For other tested yeast species, interactions with these imidazoles were predominantly indifferent or antagonistic. Triazoles, including fluconazole and voriconazole, exhibit higher binding affinity to the heme iron of Erg11p compared to imidazoles like clotrimazole and ketoconazole, contributing to differences in their pharmacokinetics and antifungal spectra^64,65,66^. Imidazoles, typically used for superficial infections due to a stronger inhibitory effect against the human CYP450 system, act via inhibition of several membrane-bound enzymes and membrane lipid biosynthesis, whereas triazoles mainly inhibit ERG11^67,68^. These pharmacodynamic differences may explain the more favourable interactions observed with triazoles. The variation in EEP interactions with triazoles among tested species may be attributed to differences in resistance gene regulation and efflux pump expression^68^. Moreover, the observed innate fluconazole resistance in *P. kudriavzevii* and potentially *C. auris* is often due to reduced Erg11p affinity, widespread ERG11 mutations, or changes in the regulation of ergosterol biosynthesis pathways^69,70^. Interactions between EEP and hydrogen peroxide were classified as additive or weakly synergistic for all examined species. This may result from the accumulation of ROS induced by both agents, along with the reported inhibition of catalase by flavonoids present in EEP, limiting cellular detoxification capacity^71^. Combination treatment with chlorhexidine and EEP yielded additive or synergistic interactions for all tested species except *P. kudriavzevii*, for which results were inconclusive. Chlorhexidine disrupts fungal membrane lipids and induces ROS accumulation, while also reducing surface hydrophobicity^72,73,74^. These effects complement those of EEP, explaining the observed synergy. In contrast, prior work on *Staphylococcus aureus* revealed antagonism, potentially due to chlorhexidine efflux pump activity enhanced under stress conditions induced by EEP^35,75,76^. Results for 2-phenoxyethanol indicated additivity or synergy with EEP across all tested species, consistent with earlier findings in *S. aureus*. In contrast, EEP combined with octenidine dihydrochloride showed antagonism or indifference, which is also in agreement with our previous bacterial studies^35^. The antifungal mechanism of 2-phenoxyethanol remains poorly understood, although studies on *Escherichia coli* suggest disruption of the membranes, inhibition of nucleic acid synthesis and malate dehydrogenase activity, and the disruption of proton gradient^77^. Given EEP’s similar cellular targets, pinpointing the precise mechanism of synergy is challenging. EEP combined with AgNPs exhibited synergy for at least two of the tested methods across all tested yeast species. AgNPs exert antifungal activity by disrupting cell walls and membranes, and inducing oxidative and osmotic stress^78,79,80,81^. These abilities overlap with those exhibited by components of EEP. Moreover, membrane damage induced by EEP may facilitate AgNPs penetration^52^. While synergy between EEP and AgNPs has not been previously reported, multiple studies have shown that AgNPs synthesized using propolis exhibit stronger antifungal activity than either compound alone^82,83,84^.

**Table 7.** Summary of the antifungal interaction types determined for selected agents against *Candida* genus representatives using different methods.

|  | *Candida albicans* |  |  |  | *Nakaseomyces glabratus* |  |  |  | *Pichia kudriavzevii* |  |  |  | *Candida auris* |  |  |  |
| --- | --- | --- | --- | --- | --- | --- | --- | --- | --- | --- | --- | --- | --- | --- | --- | --- |
|  | ChT | GK | D-D | SI | ChT | GK | D-D | SI | ChT | GK | D-D | SI | ChT | GK | D-D | SI |
| AMB | S | S | S | S | S | S | S | S | S | S | Add | S | S | S | Add | Add |
| NS | Add | Add | S | n/a | Add | Add | S | n/a | Add | Add | I | n/a | I | Add | Ant | n/a |
| CAS | Ant | Ant | Ant | n/a | Ant | Ant | Ant | n/a | I | I | Add | n/a | I | Ant | Ant | n/a |
| 5-FC | I | Add | S | n/a | Add | Add | S | n/a | Add | Add | S | n/a | I | Add | I | n/a |
| CPX | Add | Add | I | n/a | Add | S | I | n/a | Add | S | I | n/a | I | Add | I | n/a |
| FLU | I | S | S | S | Add | S | S | S | I | Ant | Ant | Ant | Ant | I | S | I |
| VOR | I | Add | S | n/a | Add | Add | I | n/a | I | Add | I | n/a | Ant | Ant | S | n/a |
| CLO | I | Add | S | Ant | I | Ant | S | Ant | I | I | I | Ant | Ant | Ant | Ant | I |
| KCA | Ant | Add | S | Ant | I | Ant | Ant | I | I | Add | Ant | Ant | Ant | I | Ant | Ant |
| H<sub>2</sub>O<sub>2</sub> | Add | S | Add | n/a | S | S | Add | n/a | Add | Add | S | n/a | Add | Add | Add | n/a |
| CHX | Add | Add | I | n/a | S | S | I | n/a | I | Add | Ant | n/a | S | S | I | n/a |
| 2-PhO-EtOH | Add | S | S | n/a | Add | Add | S | n/a | I | Add | Add | n/a | Add | Add | I | n/a |
| AgNPs | S | S | Add | n/a | S | S | Ant | n/a | S | S | Add | n/a | S | S | S | n/a |

## 4. Conclusions

The escalating burden of antifungal resistance demands the exploration of novel therapeutic strategies. Our findings suggest that the combination of propolis with polyene macrolides such as amphotericin B or nystatin holds significant promise and could lead to the future use of this combination in clinical practice. The positive interactions observed between propolis and fluconazole or voriconazole suggest clinical utility. However, careful species identification is essential prior to treatment, given the potential antagonism observed for *C. auris*. Moreover, the antagonistic interactions identified between propolis and caspofungin, clotrimazole, and ketoconazole must be widely disseminated to prevent negative therapeutic outcomes when these agents are co-administered with propolis-based products. Beyond clinical contexts, the synergistic effects of propolis with 2-phenoxyethanol and AgNPs suggest promising applications in the cosmetic and wound care industries. Harnessing the synergistic properties of natural products like propolis may contribute to the expansion of the antifungal arsenal and could inspire future approaches aimed at multi-targeted, resistance-mitigating therapies. Future work should focus on understanding the molecular mechanisms underpinning these interactions and validating their efficacy in preclinical infection models.

## 5. Materials and methods

### 5.1. Strains and cultivation

The antifungal activity of agents was investigated using four reference strains, formerly or currently recognized as *Candida* species: *Candida albicans* SC 5314, *Nakaseomyces glabratus* DSM 11226 (formerly *C. glabrata*), *Pichia kudriavzevii* (formerly *Candida krusei)* DSM 6128, and *Candida auris* B8441 (AR0387), a representative of clade I. Yeasts were routinely grown on Sabouraud Dextrose Agar (Merck KGaA, Germany). The Minimum Inhibitory Concentration (MIC) and Minimum Fungicidal Concentration (MFC) values were determined using RPMI 1640 medium (Sigma-Aldrich, Germany) with pH 7.0, supplemented with 35 g/L 3-*N*-morpholinopropanesulfonic acid (MOPS; Sigma-Aldrich, Germany), and 2% of glucose. The pH of the medium was adjusted with solid NaOH (POCH, Poland). Silver nanoparticles (AgNPs) were provided by Rafał Banasiuk and analyzed previously^85^.

### 5.2. Preparation of ethanolic extract of propolis

*Apis mellifera* propolis sample was collected at the turn of September and October 2022 and derived from an apiary located in Pruszcz Gdański, Poland. The preparation of the ethanolic extract of propolis was done as previously described^33^. In short, the extraction was carried out in the absence of light for 100 hours at ambient temperature, with continuous agitation at 50 RPM. Upon completion, the ethanolic extracts were subjected to centrifugation at 9000 RPM, followed by filtration through Millipore membranes with a pore size of 0.22 μm. The resulting filtrates were concentrated to dryness at 40 °C using a rotary vacuum evaporator. The dry, resin-like residue was subsequently weighed, and stock solutions were prepared by dissolving the extract in 70% ethanol to achieve a final concentration of 81.92 mg/mL.

### 5.3. Chemicals

Acetonitrile (gradient and LC/MS grade), water (LC/MS grade), absolute ethanol, formic acid, 2′-6′-Dihydroxy-4′-methoxydihydrochalcone were purchased from Merck (Sigma-Aldrich, Steinheim, Germany). Analytical standards of apigenin, caffeic acid, chrysin, *p*-coumaric acid, ferulic acid, galangin, isoferulic acid, kaempferol, pinobanksin, pinocembrin, pinostrobin, sakuranetin, techtochrysin vanillin were purchased from Extrasynthese. Ultrapure water (<0.06 μS/cm) was produced using Hydrolab HLP20UV (Hydrolab, Straszyn, Poland) purification system.

### 5.4. UHPLC-DAD and UHPLC-DAD-QqTOF-MS Analyses

Chromatographic analyses were performed as previously^32^. Following hardware setting was used: Thermo Scientific™ UltiMate™ 3000 system (Thermo Scientific™ Dionex™, Sunnyvale, CA, USA) connected to an autosampler, DAD detector and Compact QqTOF-MS detector (Bruker, Darmstadt, Germany). Separation was obtained on Kinetex® C18 polar 2.6 µm, 100 Å, 150 × 2.1 mm analytical column equipped with guard-column (Phenomenex, Torrence, CA, USA) and thermostated at 20 ± 1°C.

For analyses, 1 μL of the sample was injected. The mobile phase consisted of 0.1% formic acid in water (solvent A) and acetonitrile (solvent B). The mobile phase flow was set at 0.4 mL/min and following gradient was used: 95% of solvent A isocratic for 10 min, decreasing to 80% A within 1 min and isocratic for another 10 min, decreasing to 65% A within 1 min and isocratic for 8 min, decreasing to reach 40% A within 15 minutes and isocratic for another 15 min. Afterwards, the solvent composition increased to 100% B, the column was washed, and then the solvent returned to 95% A. Before each analysis, the system was stabilized. Data from DAD detector was recorded in the 200–600 nm range and at 280, 320, 360 nm. MS detector was used in both ESI negative and positive modes. The following settings were used: ion source temperature 100 °C; nebulizer gas pressure 2.0 bar; dry gas flow 0.8 L/min, 210 °C; capillary voltage 2.20 kV (negative mode) or 4.50 kV (positive mode); and collision energy 8.0 eV. Internal calibration was achieved using 10 mM solution of sodium formate clusters injected during the analysis. The collision energy for ESI-MS/MS experiments was set at 35 eV, and nitrogen was used as collision gas. The sample was filtered through CHROMAFIL® 0.2 µm, Ø13mm, H-PTFE syringe filter (Macherey-Nagel, Düren, Germany). Solutions of standard compounds were prepared in absolute ethanol and the working solutions were diluted in ethanol or ultrapure water to obtain calibration curves in a concentration range of 6.25 – 200 µg/mL. The obtained correlation values were 0.999–1.000. The derivatives of caffeic acid, *p*-coumaric, and pinobanksin were quantified as equivalents of these compounds, and the results were corrected considering the difference in molecular mass.

### 5.5. Determination of MIC and MFC values

MIC and MFC assays were performed in accordance with the Clinical Laboratory Standard Institute (CLSI) M27-A2 guidelines^86^. Prior to the experiments, yeasts were plated on Sabouraud solid medium and incubated for 24 h at 37 °C. One loop of pure culture was taken directly from the plate and transferred into a PBS (phosphate-buffered saline; Sigma-Aldrich, Germany) solution. The fungal suspensions were adjusted to an optical density of 0.1 (OD, λ=600 nm for *C. auris;* λ=660 nm for *C. albicans, N. glabratus*, and *P. kudriavzevii*) and further diluted in RPMI 1640 medium at a ratio of 1:50 (v/v) to achieve a cell count of approximately 1.0 × 10^4^ CFU/mL. Then, 100 µl of such suspension was transferred to wells of a 96-well microtiter plate (Nest Biotechnology, China) containing 100 µl of two-fold dilutions of analyzed agents in RPMI 1640 medium. Compounds were studied against pathogenic species in the following concentration ranges: 0.125 – 8 µg/mL for polyenes; 0.002 – 8 µg/mL (0.098 – 100 µg/mL against *C. auris*) for echinocandins; 0.002 – 64 µg/mL for azoles; 0.016 – 4 µg/mL (1.56 – 100 µg/mL against *P. kudriavzevii*) for 5-fluorocystosine; 0.125 – 8 µg/mL for ciclopirox; 1.563 – 100 µg/mL for terbinafine; 0.391 – 50 µg/mL for chlorhexidine; 0.098 – 25 µg/mL for octenidine dihydrochloride; 0.039 – 2.5% for 2-phenoxyethanol; 0.001 – 3.75% for hydrogen peroxide; 0.0586 – 7.5 µg/mL for AgNPs, and 4 – 256 µg/mL for EEP. Agent-free wells served as a growth control, and agent- and cell-free wells as sterility controls. Plates were incubated stationary for 24 h at 37 °C. After the incubation, the absorbance in the wells was measured at a wavelength of either 600 nm (for *C. auris)* or 660 nm (for the remaining species) using a Victor3 microplate reader (Perkin Elmer, USA). In all the experiments, the MIC value was considered to be the concentration that inhibited fungal growth by approximately 90% compared to the untreated control. The MFC value was determined by transferring each dilution used for the MIC assay on Sabouraud solid medium using a sterile 48-well microtiter plate replicator. The plates were then incubated for 24 h at 37 °C. Concentrations for which no growth was indicated were recognized as MFC. The lack of growth was also verified after 48 h of incubation. All experiments were performed in triplicate.

### 5.6. Checkerboard titration method

The possible interactions occurring between EEP and conventional/unconventional antimycotics were analyzed using the Checkerboard Titration Method^87,88^. 100 µl of inoculum of the tested yeasts (1.0 × 10^4^ CFU/mL) was added to each well on prepared 8×8 matrices (on 96-well plates) containing combinations of decreasing concentrations in RPMI 1640 medium (from 2 × MIC_90_ to 0.03125 × MIC_90_) of extracts and tested antimycotics. The plates were then incubated for 24 h at 37 °C. After incubation, the absorbance in the wells was measured at a wavelength of 600/660 nm using a Victor3 microplate reader (Perkin Elmer, USA). Once again, the MIC value was considered to be the concentration that inhibited fungal growth by approximately 90% compared to the untreated control. Next, for a given combination, the fractional inhibitory concentration index (FIC index) was calculated according to the following formula:

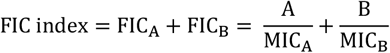

To graphically illustrate the results of the checkerboard assay, isobolograms were prepared^89^. The graph is the geometric counterpart of the equation mentioned above. If the FIC index ≤ 0.5 – the interaction is considered synergistic; if 0.5 < FIC index ≤ 1.0 – the interaction is additive; if 1 < FIC index ≤ 4 – the interaction is indifferent; if 4 < FIC index – the interaction is antagonistic^24^. All experiments were performed in triplicate.

### 5.7. Growth kinetics analysis

To determine the growth kinetics of *C. albicans, N. glabratus, P. kudriavzevii*, and *C. auris* exposed to different concentrations of antifungal agents or their mixtures with EEP, a microtiter plate-based assay was used based on the procedure described previously^32^. Briefly, two-fold dilutions of those agents were prepared in the RPMI 1640 medium. Subsequently, to 100 µL of each dilution present in the wells, an inoculum containing approximately 1.0 × 10^4^ CFU/mL was added, resulting in a final volume of 200 μL. Microbial growth kinetics were monitored over 24 hours using the SPARK® multimode microplate reader (Tecan, Switzerland). At hourly intervals, the turbidity of the culture was read by measuring absorbance at 600/660 nm, with agitation for 10 seconds preceding each optical density measurement. XY graphs were constructed using GraphPad Prism® 8.0.2 (GraphPad Software, Inc., USA). All experiments were performed in triplicate.

### 5.8. Disc-diffusion assay

Overnight cultures in Sabouraud broth were adjusted to 0.5 McFarland units and spread on Mueller-Hinton plates with 2% glucose. Then, on agar were placed antibiotic discs (Liofilchem, Italy) or blank discs, onto which 10 µL of antifungal agent was added. After 15 minutes, 10 µL of 70% ethanol as a control or EEP was added onto the discs. After overnight incubation at 37 °C, inhibition zones were measured. The decrease of the zone higher than −1 mm was considered antagonism, the difference between −1 mm or equal and +1 mm or equal was considered indifference, the increase higher than 1 mm up to 2 mm was considered additivity, and the increase higher than 2 mm was considered synergy. The list of antifungal concentrations is in the supplementary data - Table S20. All experiments were performed in triplicate. Interactions were interpreted based on the difference in inhibition zone diameter (mm) between the combination and the antifungal alone. An increase in the inhibition zone greater than 2 mm was interpreted as a synergistic interaction. An increase between 1 and 2 mm indicated additivity, while changes within ±1 mm were considered indicative of indifference. A decrease greater than 1 mm was classified as antagonism. This approach is consistent with previously described disc diffusion-based methods used to assess antimicrobial interactions in both bacterial and fungal models by applying additional compounds to discs and evaluating changes in inhibition zone diameter ^90,91,92^.

### 5.9. Simulated infection model

Overnight cultures were centrifuged (5 min, 10 000 RCF) and suspended in RPMI medium. Cultures were adjusted to 0.5 McFarland units in tubes with 5 ml of RPMI, then 2xMIC concentrations of amphotericin B, fluconazole, clotrimazole, ketoconazole, or EEP were added alone or as a combination of antifungal plus EEP. The tubes were incubated overnight at 37 °C with shaking at 180 RPM. The next day, cultures were centrifuged, washed with PBS, and resuspended in RPMI. The cultures were again adjusted to 0.5 McFarland units, and the antifungals or EEP were added. The cycle was repeated for 5 days. The cultures were then serially diluted in PBS and spread onto Sabouraud Agar. The plates were incubated at 37 °C overnight, and then the CFU/mL value was determined. From the mean CFU/mL value estimated for the control (no treatment), the mean CFU/mL values obtained for drug alone, EEP alone, or a combination of drug plus EEP, respectively, were subtracted. Then, the differences for the drug alone and for EEP alone were subtracted from the difference for the combination. If the CFU/mL difference of the combination was greater than 0.5, then the differences for drug and EEP together, the interaction was considered synergistic. The difference between 0 and 0.5 was considered additivity, the difference between −0.5 and 0 was considered indifference, and the difference lower than −0.5 was considered antagonism. For *C. albicans*, YNB (Yeast Nitrogen Base; Sigma-Aldrich, Germany) medium with 2% glucose was used due to intensive yeast-to-hyphae transition and problems with CFU/mL determination in RPMI medium. All experiments were performed in triplicate.

## 6. Acknowledgments

The authors would like to express gratitude to Rafał Banasiuk for providing silver nanoparticles. The language of the manuscript was improved with the use of Grammarly software.

## 7. Authors’ contributions

P.B. did the experimental work on MIC/MFC, checkerboard, and growth kinetics, analysed the data, participated in the conception of the study, and participated in writing the manuscript. M.K.P. did the experimental work on disc diffusion and simulated infection, analysed the data, participated in the conception of the study, and wrote the manuscript. P.M.K. did the experimental work on chemical characterisation of EEP, analysed the data, and participated in writing the manuscript. P.V.D. participated in the conception of the study, analysed the data, and revised the manuscript. P.S. led the design and conception of the study, coordinated the work, analysed the data, substantially revised the manuscript, and provided funding. All the authors have read and approved the final manuscript.

## 8. Data availability

All data supporting the findings of this study are available within the paper and its Supplementary Information. Should any raw data files be needed, they are available from the corresponding authors (MKP, PS) upon reasonable request.

## 9. Declaration of competing interests

The authors report no conflict of interest.

## 10. Funding

The research was financed by the grant UMO-2020/39/B/NZ7/02901 from the National Science Centre, Poland.

## Notes

### Competing Interest Statement

The authors have declared no competing interest.

