## Supplement for "Ethanolic Extract of Polish Propolis exhibits synergy with selected antifungal agents against yeast pathogens causing candidiasis"

Table S1. Compounds detected in the investigated propolis by UHPLC-QqTOF-DAD-MS.

| No. | Compound | RT [min.] | Molecular formula | UV max [nm] | $[M-H^+]^-$ | [M+H+]+ / [M+Na+]+ | $MS^2$ Base peak (ESI-) | $MS^2$ Base peak (ESI+) | Error (ESI-) [ppm] | Error (ESI+) [ppm] | Abundance (ESI-) | Abundance (ESI+) | References |
|---|---|---|---|---|---|---|---|---|---|---|---|---|---|
| **1** | Protocatechuic acid [a, b, c] | 4.30 | $C_7H_6O_4$ | 260, 294 | 153.0193 | 155.0342 | 108.0206 | - | -0.2 | 2.0 | +++ | ++ | 1 |
| **2** | Protocatechuic aldehyde (3,4-dihydroxybenzaldehyde) [b,c] | 6.53 | $C_7H_6O_3$ | 202, 229, 280, 311 | 137.0244 | 139.0391 | 108.0206 | 125.9902 | -0.1 | 0.9 | ++++ | +++ | 2 |
| **3** | 4-Hydroxybenzoic acid [a,b,c] | 7.90 | $C_7H_6O_3$ | 256 | 137.0241 | 139.0390 | 137.0984 | 121.0287 | -2.3 | 0.2 | +++ | ++ | 3, 4, 5 |
| **4** | 4-Hydroxybenzaldehyde [b,c] | 10.88 | $C_7H_6O_2$ | 284 | 121.0293 | 123.0441 | - | 95.0465 | -1.7 | 0.4 | +++ | ++ | 3, 5, 6 |
| **5** | Caffeic acid [a,b,c] | 14.20 | $C_9H_8O_4$ | 323 | 179.0351 | 181.0500 | 135.0453 | 117.0333 | 0.6 | 2.6 | ++++ | ++++ | 3, 4, 7, 8 |
| **6** | Vanillin (4-hydroxy-3-methoxybenzaldehyde) [a, b] | 14.74 | $C_8H_8O_3$ | 309, 280 | 151.0396 | 153.0549 | 108.0192 | 110.0353 | -3.1 | 1.8 | +++ | +++ | 9 |
| **7** | *p*-Coumaric acid [a,b,c] | 15.66 | $C_9H_8O_3$ | 310 | 163.0402 | 165.0553 | 119.0498 | 119.0486 | 0.8 | 4.1 | ++++ | ++++ | 3, 4, 7, 8 |
| **8** | Benzoic acid a, b, c | 15.87 | $C_7H_6O_2$ | 274 | 121.0292 | 123.0441 | - | 105.034 | -2.5 | 0.4 | +++ | +++ | 10 |
| **9** | Ferulic acid [a,b,c] | 16.38 | $C_{10}H_{10}O_4$ | 322, 298 | 193.0507 | 195.0659 | 134.0288 | 117.0331 | 0.3 | 3.7 | ++++ | +++ | 3, 4, 7, 8 |
| **10** | Isoferulic acid [a,b,c] | 16.86 | $C_{10}H_{10}O_4$ | 323, 298 | 193.0508 | 195.066 | 134.0292 | 134.00362 | 0.9 | 4.2 | ++++ | ++++ | 3, 4, 7, 8 |
| **11** | Dihydrokaempferol (Aromadendrin) [b,c] | 20.20 | $C_{15}H_{12}O_6$ | 292 | 287.0561 | 289.0711 | 125.0243 | 215.0192 | -0.1 | 1.5 | ++++ | +++ | 3 |
| **12** | Methyl ferulate [b,c] | 21.00 | $C_{11}H_{12}O_4$ | 322, 289 | 207.0669 | 209.0818 | 132.0198 | 148.0521 | 3.0 | 4.6 | ++++ | ++++ | 3 |
| **13** | Cinnamic acid [a,b] | 23.38 | $C_9H_8O_2$ | 280 | 147.0450 | 149.0594 | 117.0334 | 103.0542 | -1.1 | -2.0 | ++ | ++ | 3, 4, 7, 8 |
| **14** | Apigetrin (Apigenin 7-*O*-glucoside) [b,c] | 23.78 | $C_{21}H_{20}O_{10}$ | 265 | 431.0994 | 433.1137 | 268.0384 | 271.0599 | 2.4 | 1.8 | +++ | +++ | 3, 7, 8 |
| **15** | Pinobanksin 5-methylether [b] | 25.24 | $C_{16}H_{14}O_5$ | 288 | 285.0777 | 287.0921 | 252.0444 | 91.0546 | 3.0 | 2.4 | ++++ | ++++ | 3, 4, 7, 8 |
| **16** | Quercetin [a,b,c] | 25.74 | $C_{15}H_{10}O_7$ | 369, 270sh, 255 | 301.0359 | 303.0509 | 151.004 | 303.0509 | 1.7 | 3.2 | ++++ | ++++ | 3, 4, 7, 8 |
| **17** | Luteolin [a, b, c] | 25.80 | $C_{15}H_{10}O_6$ | 352, 290sh,255 | 285.0420 | 287.0558 | 133.0295 | 287.0558 | 5.4 | 2.7 | ++++ | +++ | 12 |
| **18** | Quercetin 3-methyl ether [b,c] | 26.19 | $C_{16}H_{12}O_7$ | 255, 269sh, 356 | 315.0518 | 317.0665 | 271.0252 | 302.0427 | 2.4 | 2.9 | ++++ | ++++ | 3, 4, 7, 8 |
| **19** | Pinobanksin [a,b,c] | 26.57 | $C_{15}H_{12}O_5$ | 292 | 271.0623 | 273.0766 | 197.0618 | 199.0761 | 4.1 | 3.1 | ++++ | ++++ | 3, 4, 7, 8 |
| **20** | Naringenin [a,b,c] | 26.86 | $C_{15}H_{12}O_5$ | 288 | 271.0614 | 273.0765 | 119.0501 | 153.0188 | 0.7 | 2.7 | ++++ | +++ | 3, 4, 7, 8 |
| **21** | Alpinetin (Pinocembrin-5-methyl-ether) [b,c] | 27.09 | $C_{16}H_{14}O_4$ | 286 | 269.0826 | 271.0974 | 149.996 | 167.0345 | 2.5 | 3.4 | ++++ | ++++ | 13 |
| **22** | Apigenin [a,b,c] | 27.30 | $C_{15}H_{10}O_5$ | 335, 290sh, 268 | 269.0463 | 271.0608 | 117.0349 | 271.0608 | 2.8 | 2.6 | ++++ | ++++ | 3, 4, 7, 8 |
| **23** | Kaempferol [a,b,c] | 27.58 | $C_{15}H_{10}O_6$ | 365, 295sh, 266 | 285.0412 | 287.0559 | 285.0413 | 287.0559 | 2.6 | 3.1 | ++++ | ++++ | 3, 4, 7, 8 |
| **24** | Isorhamnetin [a,b,c] | 27.75 | $C_{16}H_{12}O_7$ | 369, 300sh,255 | 315.0523 | 317.0665 | 300.0284 | 317.0665 | 4.0 | 2.9 | ++++ | ++++ | 14 |

| | | | | | | | | | | | | | |
|---|---|---|---|---|---|---|---|---|---|---|---|---|---|
| **25** | Rhamnetin [a,b,c] | 27.94 | $C_{16}H_{12}O_7$ | 369, 300sh,256 | 315.052 | 317.0662 | 300.0288 | 317.0662 | 3.1 | 2.0 | ++++ | +++ | 15 |
| **26** | 1,2-di-*p*-Coumaroylglycerol isomer I [b] | 28.14 | $C_{21}H_{20}O_7$ | 311, 300sh | 383.1135 | 385.1285 | 163.0398 | - | -0.3 | 0.8 | +++ | ++ | 3, 4, 7, 8 |
| **27** | Luteolin-5-methyl ether [b] | 28.24 | $C_{16}H_{12}O_6$ | 346, 298sh, 267 | 299.0570 | 301.0713 | 255.0307 | 286.048 | 3.0 | 2.1 | ++++ | ++++ | 3, 4, 7, 8 |
| **28** | Quercetin-dimethyl-ether [b] | 28.50 | $C_{17}H_{14}O_7$ | 256, 352 | 329.0671 | 331.0817 | 299.0201 | 316.0582 | 1.2 | 1.4 | ++++ | ++++ | 3, 4, 7, 8 |
| **29** | Galangin-5-methyl-ether [b,c] | 28.73 | $C_{16}H_{12}O_5$ | 352, 300sh, 260 | 283.0618 | 285.0765 | 211.0407 | 270.0526 | 2.1 | 2.6 | ++++ | ++++ | 3, 4, 7, 8 |
| **30** | 5-Methyl-pinobanksin-3- acetate [b] | 28.93 | $C_{18}H_{16}O_6$ | 286 | 327.0880 | 329.1025 | 224.048 | 213.0907 | 1.8 | 1.6 | ++++ | +++ | 7, 8 |
| **31** | Quercetin-methyl-ether [b] | 29.86 | $C_{16}H_{12}O_7$ | 255, 272sh, 369 | 315.0518 | 317.0665 | 165.0198 | 317.0665 | 2.4 | 2.9 | ++++ | ++++ | 3, 4, 7, 8 |
| **32** | Caffeic acid butyl or isobutyl ester isomer isomer I [b] | 30.54 | $C_{13}H_{16}O_4$ | 320 | 235.0979 | 237.1128 | 133.0297 | - | 1.3 | 2.8 | +++ | + | 3, 4, 7, 8 |
| **33** | Caffeic acid prenyl ester isomer [b] | 30.75 | $C_{14}H_{16}O_4$ | 320 | 247.0972 | 249.1121 | 133.029 | - | -1.6 | -0.1 | +++ | ++ | 3, 4, 7, 8 |
| **34** | Quercetin-dimethyl-ether [b] | 31.04 | $C_{17}H_{14}O_7$ | 254, 270sh, 356 | 329.0674 | 331.0818 | 299.0204 | 316.0582 | 2.2 | 1.7 | ++++ | ++++ | 3, 4, 7, 8 |
| **35** | Caffeic acid 2-methyl-2-butenyl ester [b] | 31.43 | $C_{14}H_{16}O_4$ | 325, 296sh | 247.0982 | 249.1127 | 135.0451 | 163.0396 | 2.5 | 2.3 | ++++ | ++++ | 3, 4, 7, 8 |
| **36** | Caffeic acid 3-methyl-2-butenyl ester (Basic prenyl ester) [b] | 32.12 | $C_{14}H_{16}O_4$ | 325, 298 sh | 247.0989 | 249.1125 | 133.0299 | 163.0395 | 5.3 | 1.5 | ++++ | +++ | 3, 4, 7, 8 |
| **37** | Caffeic acid 3-methyl-3-butenyl ester [b] | 32.41 | $C_{14}H_{16}O_4$ | 325, 298 sh | 247.0986 | 249.1124 | 133.0298 | 163.0394 | 4.1 | 1.1 | ++++ | ++ | 3, 4, 7, 8 |
| **38** | Caffeic acid benzyl ester [b] | 32.94 | $C_{16}H_{14}O_4$ | 327, 296sh | 269.0834 | 271.0975 | 133.0297 | 117.0698 | 5.4 | 3.7 | ++++ | ++++ | 3, 4, 7, 8 |
| **39** | Chrysin [a, b, c] | 33.27 | $C_{15}H_{10}O_4$ | 312sh, 268 | 253.0520 | 255.0657 | 253.0513 | 255.0657 | 5.4 | 2.0 | ++++ | ++++ | 3, 4, 7, 8 |
| **40** | Sakuranetin a,b,[c] | 33.77 | $C_{16}H_{14}O_5$ | 289 | 285.0779 | 287.0919 | 119.0503 | 167.0341 | 3.7 | 1.7 | ++++ | +++ | 16 |
| **41** | Pinocembrin a,[b, c] | 33.99 | $C_{15}H_{12}O_4$ | 290 | 255.0676 | 257.0812 | 171.0463 | 153.0185 | 5.2 | 1.4 | ++++ | ++++ | 3, 4, 7, 8 |
| **42** | Isosakuranetin a,[b, c] | 34.42 | $C_{16}H_{14}O_5$ | 292 | 285.0784 | 287.0919 | 164.0112 | 153.0185 | 5.4 | 1.7 | ++++ | ++++ | 3, 4, 7, 8 |
| **43** | Pinobanksin acetate isomer*[, b] | 34.52 | $C_{17}H_{14}O_6$ | 293 | 313.0733 | 315.0862 | 253.0514 | 153.0184 | 4.9 | -0.4 | ++++ | ++++ | 3, 4, 7, 8 |
| **44** | Galangin [a,b] | 34.71 | $C_{15}H_{10}O_5$ | 358, 266 | 269.0469 | 271.0605 | 269.0472 | 271.0605 | 5.0 | 1.5 | ++++ | ++++ | 3, 4, 7, 8 |
| **45** | Caffeic acid phenethyl ester (CAPE) [b,c] | 35.26 | $C_{17}H_{16}O_4$ | 326 | 283.0991 | 285.1123 | 135.0459 | 105.0698 | 5.4 | 0.6 | ++++ | ++++ | 3, 4, 7, 8 |
| **46** | Pinobanksin 3-*O*-acetate [b] | 35.83 | $C_{17}H_{14}O_6$ | 294 | 313.0733 | 315.0864 | 253.0523 | 199.0759 | 4.9 | 0.3 | ++++ | ++++ | 3, 4, 7, 8 |
| **47** | 2-Acetyl-1,3-di-*p*-coumaroylglycerol [b] | 36.11 | $C_{23}H_{22}O_8$ | 312 (296) | 425.1252 | -/449.1211 | 163.0404 | 147.0442 | 2.4 | 0.9 | ++++ | +++ | 3, 4, 7, 8 |
| **48** | 2-Acetyl-3-*p*-coumaroyl-1-feruloylglycerol [b] | 36.42 | $C_{24}H_{24}O_9$ | 316 | 455.1360 | -/479.1323 | 163.0408 | 177.0549 | 2.7 | 2.2 | +++ | +++ | 3, 4, 7, 8 |
| **49** | 2-Acetyl-1,3-di-feruloylglycerol [b] | 36.77 | $C_{25}H_{26}O_{10}$ | 324 | 485.1458 | -/509.1437 | 193.051 | 177.0546 | 1.0 | 3.7 | +++ | ++ | 3, 4, 7, 8 |
| **50** | Ermanin (Kaempferol-3,4'-dimethyleter) [b] | 36.84 | $C_{17}H_{14}O_6$ | 350, 267 | 313.0731 | 315.0867 | 283.0256 | 300.0627 | 4.3 | 1.2 | ++++ | +++ | 3, 4, 5, 8 |
| **51** | Methoxychrysin [b] | 36.89 | $C_{15}H_{10}O_5$ | 310sh, 268 | 269.0475 | 271.0612 | 211.1827 | - | 7.2 | 4.1 | +++ | ++ | 3, 4, 7, 8 |
| **52** | *p*-Coumaric acid 3-methyl-3-butenyl ester [b] | 37.07 | $C_{14}H_{16}O_3$ | 311 | 231.1030 | 233.1175 | 117.0347 | 147.0441 | 1.5 | 1.2 | ++++ | +++ | 3, 4, 7, 8 |
| **53** | *p*-Coumaric acid 3-methyl-3-butenyl ester [b] | 37.58 | $C_{14}H_{16}O_3$ | 316 | 231.1031 | 233.1179 | 117.0348 | - | 1.9 | 2.9 | ++++ | ++ | 3, 4, 7, 8 |

| | | | | | | | | | | | | | |
|---|---|---|---|---|---|---|---|---|---|---|---|---|---|
| **54** | *p*-Coumaric acid 3-methyl-3-butenyl ester [b] | 37.80 | $C_{14}H_{16}O_3$ | 311 | 231.1038 | 233.1182 | 117.0349 | - | 4.9 | 4.2 | ++++ | ++ | 3, 4, 7, 8 |
| **55** | *p*-Coumaric acid 3-methyl-3-butenyl ester [b] | 37.94 | $C_{14}H_{16}O_3$ | 312 | 231.1037 | 233.1179 | 117.0345 | - | 4.5 | 2.9 | ++++ | ++ | 3, 4, 7, 8 |
| **56** | *p*-Coumaric acid benzyl ester [b] | 38.01 | $C_{16}H_{14}O_3$ | 320 | 253.0883 | 255.1018 | 117.0348 | - | 5.1 | 0.9 | ++++ | +++ | 3, 4, 7, 8 |
| **57** | Caffeic acid cinnamyl ester [b] | 38.42 | $C_{18}H_{16}O_4$ | 325 | 295.0982 | 297.1129 | 134.0379 | 117.0699 | 2.2 | 2.6 | ++++ | ++ | 3, 4, 7, 8 |
| **58** | Ferulic acid benzyl ester [b] | 38.55 | $C_{17}H_{16}O_4$ | 325 | 283.0989 | 285.1126 | 133.0298 | - | 4.6 | 1.6 | +++ | +++ | 3, 4, 7, 8 |
| **59** | Pinobanksin-3-*O*-propanoate [b] | 39.57 | $C_{18}H_{16}O_6$ | 295 | 327.0891 | 329.1025 | 253.0521 | 199.0762 | 5.2 | 1.6 | ++++ | ++++ | 3, 4, 7, 8 |
| **60** | *p*-Coumaric acid phenethyl ester [b] | 39.61 | $C_{17}H_{16}O_3$ | 310 | 267.1039 | 269.1169 | 119.0503 | - | 4.6 | -1.2 | ++++ | +++ | 3, 4, 7, 8 |
| **61** | *p*-Coumaric acid pentyl or isopentyl ester [b] | 40.11 | $C_{14}H_{18}O_3$ | 292, 309 | 233.1191 | 235.1338 | 117.0336 | - | 3.3 | 4.0 | ++++ | +++ | 15, 17 |
| **62** | 2′-6′-Dihydroxy-4′-methoxydihydrochalcone [a, b] | 41.12 | $C_{16}H_{16}O_4$ | 286 | 271.0984 | 273.1121 | 152.0114 | 91.0547 | 3.0 | -0.1 | ++++ | +++ | 18 |
| **63** | Pinobanksin 3-*O*-butanoate or isobutanoate [b] | 41.24 | $C_{19}H_{18}O_6$ | 286 | 341.1039 | 343.1183 | 253.0515 | 153.0184 | 2.4 | 2.0 | +++ | ++ | 3, 4, 5, 7 |
| **64** | Pinobanksin 3-*O*-pentenoate or isopentenoate isomer I [b] | 41.84 | $C_{20}H_{18}O_6$ | 292 | 353.1027 | 355.1153 | 253.0488 | - | -1.0 | -6.5 | +++ | ++ | 3, 4, 5, 7 |
| **65** | *p*-Coumaric acid cinnamyl ester [b] | 41.91 | $C_{18}H_{16}O_3$ | 309 | 279.1037 | - | 117.0342 | - | 3.7 | - | ++++ | − | 3, 4, 5, 7 |
| **66** | Tectochrysin [a, b] | 42.10 | $C_{16}H_{12}O_4$ | 269 | - | 269.0801 | 226.0635 | 269.0801 | - | -2.6 | − | ++++ | 8 |
| **67** | Pinostrobin [a, b] | 42.27 | $C_{16}H_{14}O_4$ | 291 | - | 271.0970 | 167.035 | - | - | 1.9 | − | ++++ | 3, 8 |
| **68** | Pinobanksin 3-*O*-pentenoate or isopentenoate isomer II [b] | 42.87 | $C_{20}H_{18}O_6$ | 292 | 353.1048 | 355.1182 | 253.0516 | - | 4.9 | 1.6 | ++++ | +++ | 3, 4, 5, 7 |
| **69** | Pinobanksin 3-*O*-benzoate [b] | 43.85 | $C_{22}H_{16}O_6$ | 286 | 375.0877 | 377.1030 | 253.0514 | 105.0337 | 0.8 | 2.7 | ++++ | +++ | 19 |
| **70** | Pinobanksin 3-*O*-pentanoate or isopentenoate isomer I [b] | 44.75 | $C_{20}H_{20}O_6$ | 290 | 355.1196 | 357.1340 | 253.0505 | 227.0713 | 2.5 | 2.1 | ++++ | ++ | 3, 4, 7, 8 |
| **71** | Pinobanksin 3-*O*-pentanoate or isopentenoate isomer II [b] | 45.01 | $C_{20}H_{20}O_6$ | 293 | 355.1197 | 357.1339 | 253.0515 | 199.0764 | 2.8 | 1.8 | ++++ | +++ | 3, 4, 7, 8 |
| **72** | Pinobanksin 3-*O*-hexenoate or isohexenoate [b] | 45.63 | $C_{21}H_{20}O_6$ | 292 | 367.1177 | - | 2530502 | - | -2.8 | - | +++ | − | 20 |
| **73** | Pinobanksin-3-*O*-cinnamate [b] | 45.94 | $C_{24}H_{18}O_6$ | 292 | 401.1035 | 403.1196 | 253.0509 | - | 1.1 | 4.9 | +++ | ++ | 21 |
| **74** | Pinobanksin-3-*O*-hydroxycinnamate [b] | 46.31 | $C_{24}H_{20}O_6$ | 292 | 403.1207 | 405.1345 | 253.0516 | 105.0698 | 4.9 | 3.0 | ++++ | ++++ | 19 |
| **75** | Metoxycinnamic acid cinnamyl ester [b] | 46.42 | $C_{18}H_{30}O_3$ | 282 | 293.2130 | 295.2277 | 293.2121 | 165.1279 | 2.7 | 3.2 | ++++ | ++++ | 3, 4, 7, 8 |
| **76** | Pinobanksin 3-*O*-hexanoate or isohexanoate isomer I [b] | 46.76 | $C_{21}H_{22}O_6$ | 286 | 369.1349 | 371.1484 | 253.0506 | 227.0697 | 1.4 | -1.4 | ++++ | ++ | 3, 4, 7, 8 |
| **77** | Pinobanksin 3-*O*-hexanoate or isohexanoate isomer II [b] | 47.07 | $C_{21}H_{22}O_6$ | 282 | 369.1350 | 371.1489 | 253.0509 | - | 1.7 | 0.0 | ++++ | +++ | 3, 4, 7, 8 |

[a] identified by comparison with standard compound; [b] identified by exact mass, $MS^2$ and/or UV, comparison with literature data; [c] identified using $MS^2$ databases ; * tentatively identified; + / ++ / +++ / ++++ compound detected <1000 / >1000 / >10000 / >100000 counts, respectively; − compound not detected.

### References to table S1

14. Pobłocka-Olech, L.; Głód, D.; Jesionek, A.; Łuczkiewicz, M.; Krauze-Baranowska, M. Studies on the

Polyphenolic Composition and the Antioxidant Properties of the Leaves of Poplar (*Populus* Spp.) Various Species and Hybrids. *Chem. Biodivers.* **2021**, *18*, e2100227, doi:10.1002/CBDV.202100227;PAGE:STRING:ARTICLE/CHAPTER.

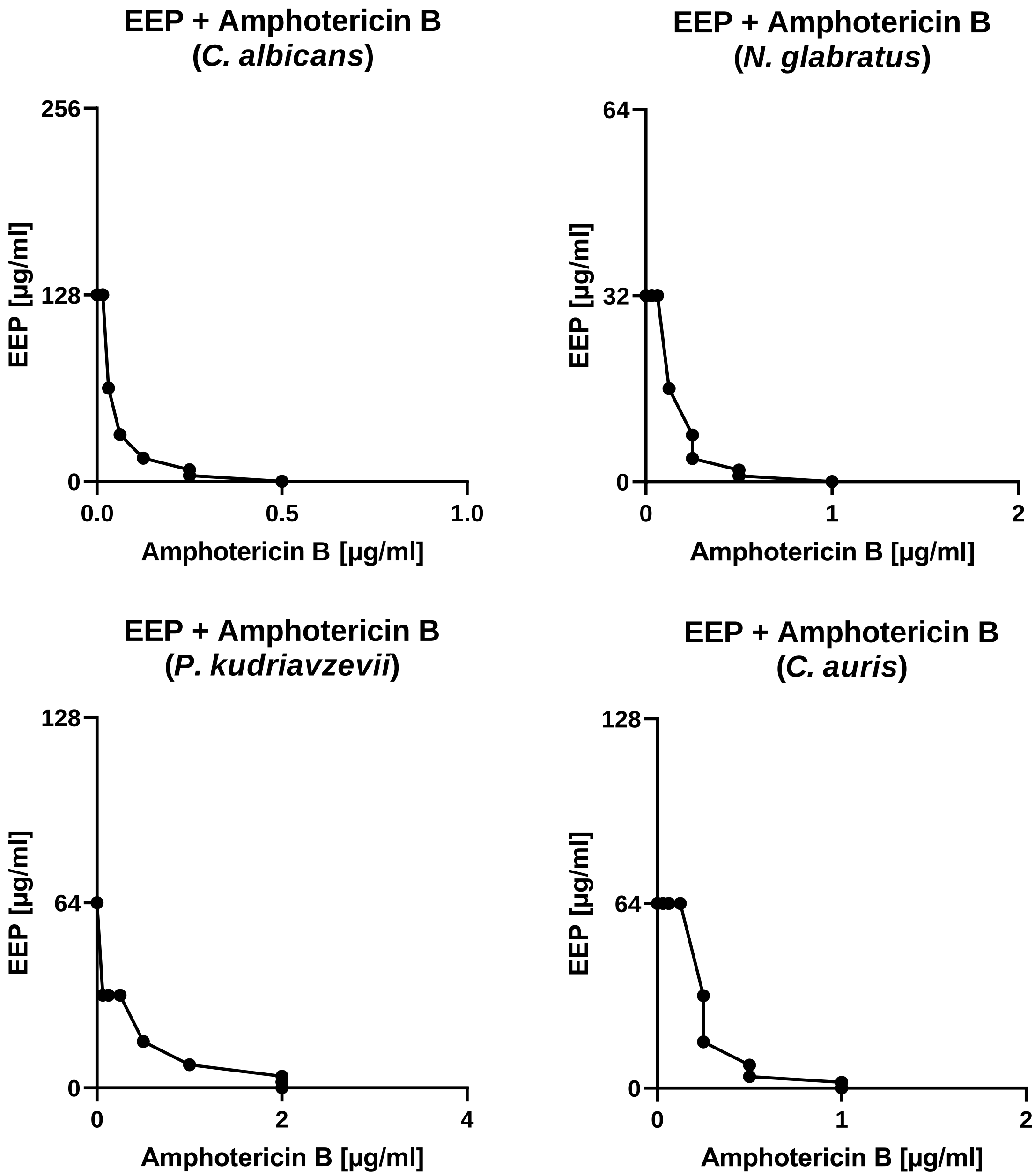


Figure S1 – Isobolograms presenting the dependence between EEP and amphotericin B concentrations regarding $MIC_{90}$ values determined against selected yeast pathogens causing candidiasis.

Table S2 – Summary of estimated fractional inhibitory concentrations and antifungal interaction types between amphotericin B and EEP against selected yeast pathogens causing candidiasis.

| Microorganism | Agent | MIC (μg/mL) | | FIC | ∑FIC | Effect |
|---|---|---|---|---|---|---|
| | | Alone | In combination | | | |
| *Candida albicans* SC 5314 | Amphotericin B | 0.5 | 0.0625 | 0.125 | 0.375 | Synergy |
| | EEP | 128 | 32 | 0.25 | | |
| *Nakaseomyces glabratus* DSM 11226 | Amphotericin B | 1 | 0.25 | 0.25 | 0.5 | Synergy |
| | EEP | 32 | 8 | 0.25 | | |
| *Pichia kudriavzevii* DSM 5784 | Amphotericin B | 2 | 0.5 | 0.25 | 0.5 | Synergy |
| | EEP | 64 | 16 | 0.25 | | |
| *Candida auris* B 8441 | Amphotericin B | 0.5 | 0.25 | 0.25 | 0.5 | Synergy |
| | EEP | 64 | 16 | 0.25 | | |

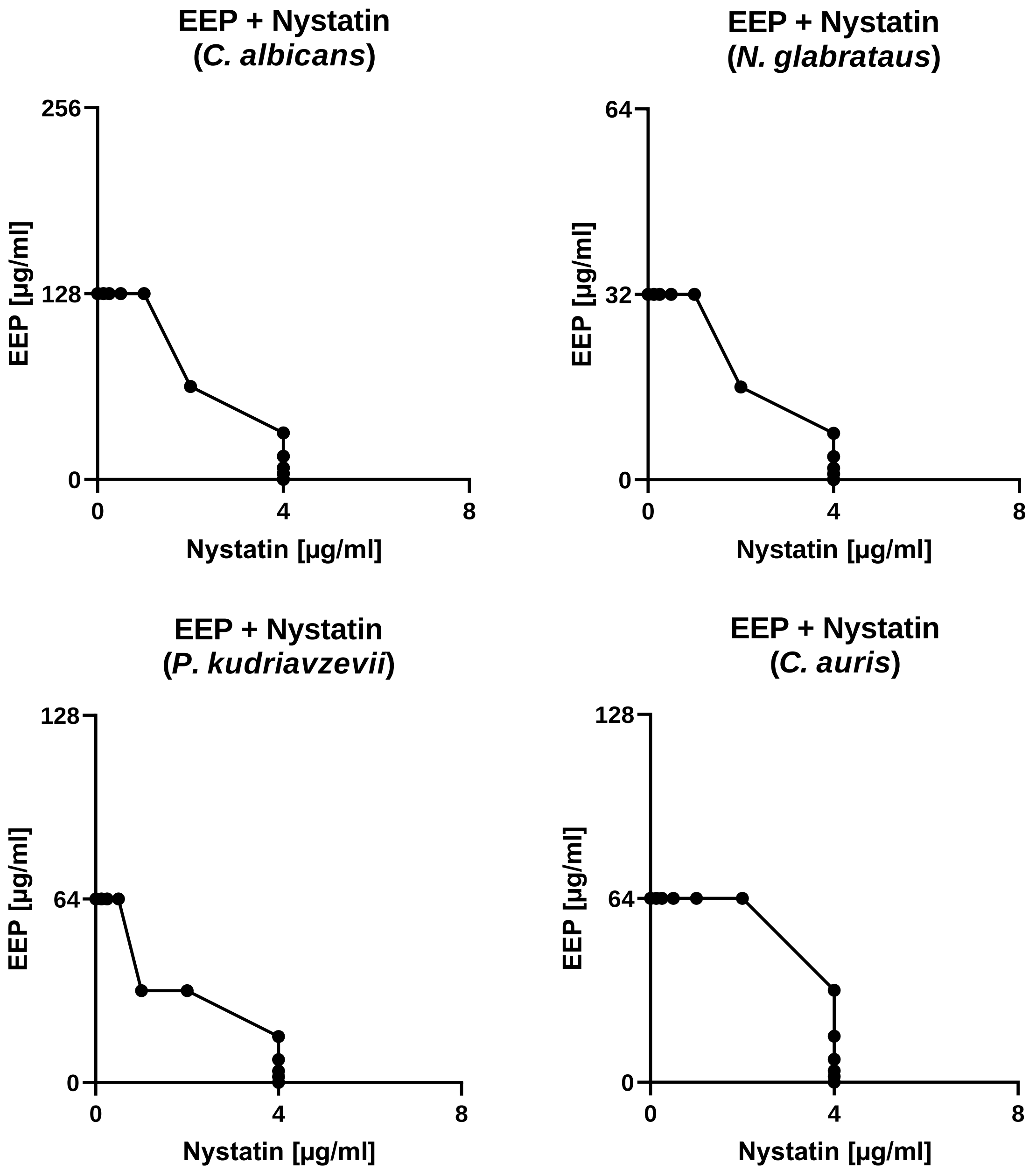


Figure S2 – Isobolograms presenting the dependence between EEP and nystatin concentrations regarding $MIC_{90}$ values determined against selected yeast pathogens causing candidiasis.

Table S3 – Summary of estimated fractional inhibitory concentrations and antifungal interaction types between nystatin and EEP against selected yeast pathogens causing candidiasis.

| Microorganism | Agent | MIC (µg/mL) | | FIC | ∑FIC | Effect |
|---|---|---|---|---|---|---|
| | | Alone | In combination | | | |
| *Candida albicans* SC 5314 | Nystatin | 4 | 2 | 0. 5 | 1 | Additivity |
| | EEP | 128 | 64 | 0.5 | | |
| *Nakaseomyces glabratus* DSM 11226 | Nystatin | 4 | 2 | 0.5 | 1 | Additivity |
| | EEP | 32 | 16 | 0.5 | | |
| *Pichia kudriavzevii* DSM 5784 | Nystatin | 4 | 1 | 0.25 | 0.75 | Additivity |
| | EEP | 64 | 32 | 0.5 | | |
| *Candida auris* B 8441 | Nystatin | 4 | 4 | 1 | 1.03125 | Indifference |
| | EEP | 64 | 2 | 0.03125 | | |

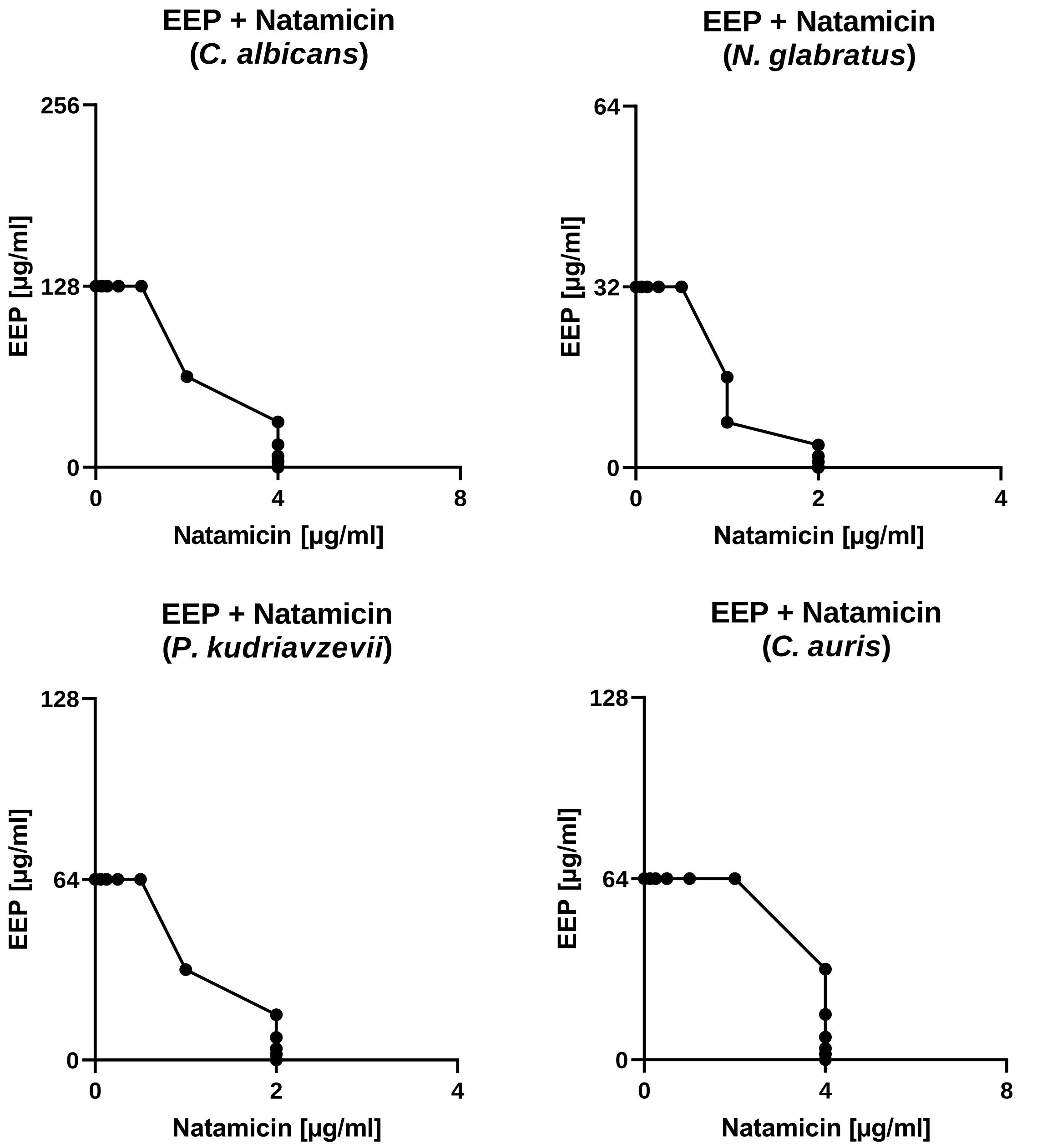


Figure S3 – Isobolograms presenting the dependence between EEP and natamycin concentrations regarding $MIC_{90}$ values determined against selected yeast pathogens causing candidiasis.

Table S4 – Summary of estimated fractional inhibitory concentrations and antifungal interaction types between natamycin and EEP against selected yeast pathogens causing candidiasis.

| Microorganism | Agent | MIC (μg/mL) | | FIC | ∑FIC | Effect |
|---|---|---|---|---|---|---|
| | | Alone | In combination | | | |
| *Candida albicans* SC 5314 | Natamycin | 4 | 2 | 0.5 | 1 | Additivity |
| | EEP | 128 | 64 | 0.5 | | |
| *Nakaseomyces glabratus* DSM 11226 | Natamycin | 2 | 1 | 0.5 | 0.75 | Additivity |
| | EEP | 32 | 8 | 0.25 | | |
| *Pichia kudriavzevii* DSM 5784 | Natamycin | 2 | 1 | 0.5 | 1 | Additivity |
| | EEP | 64 | 32 | 0.5 | | |
| *Candida auris* B 8441 | Natamycin | 4 | 4 | 1 | 1.03125 | Indifference |
| | EEP | 64 | 2 | 0.03125 | | |

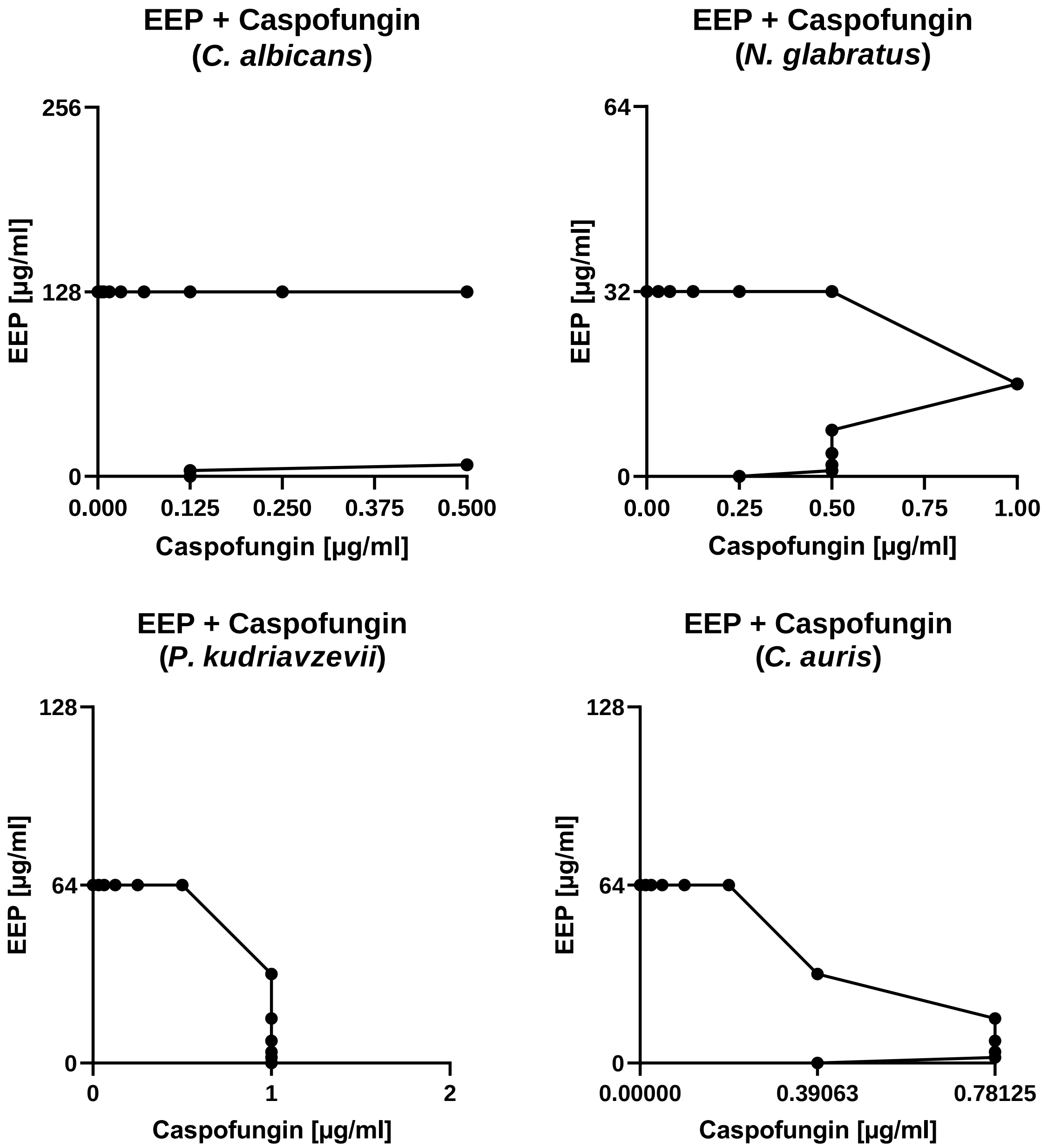


Figure S4 – Isobolograms presenting the dependence between EEP and caspofungin concentrations regarding $MIC_{90}$ values determined against selected yeast pathogens causing candidiasis.

Table S5 – Summary of estimated fractional inhibitory concentrations and antifungal interaction types between caspofungin and EEP against selected yeast pathogens causing candidiasis.

| Microorganism | Agent | MIC (µg/mL) | | FIC | ∑FIC | Effect |
|---|---|---|---|---|---|---|
| | | Alone | In combination | | | |
| *Candida albicans* SC 5314 | Caspofungin | 0.125 | >0.5 | >4 | >4.0625 | Antagonism |
| | EEP | 128 | 8 | 0.0625 | | |
| *Nakaseomyces glabratus* DSM 11226 | Caspofungin | 0.25 | 1 | 4 | 4.5 | Antagonism |
| | EEP | 32 | 16 | 0.5 | | |
| *Pichia kudriavzevii* DSM 5784 | Caspofungin | 1 | 1 | 1 | 1.03125 | Indifference |
| | EEP | 64 | 2 | 0.03125 | | |
| *Candida auris* B 8441 | Caspofungin | 0.390625 | 0.78125 | 2 | 2.03125 | Indifference |
| | EEP | 64 | 2 | 0.03125 | | |

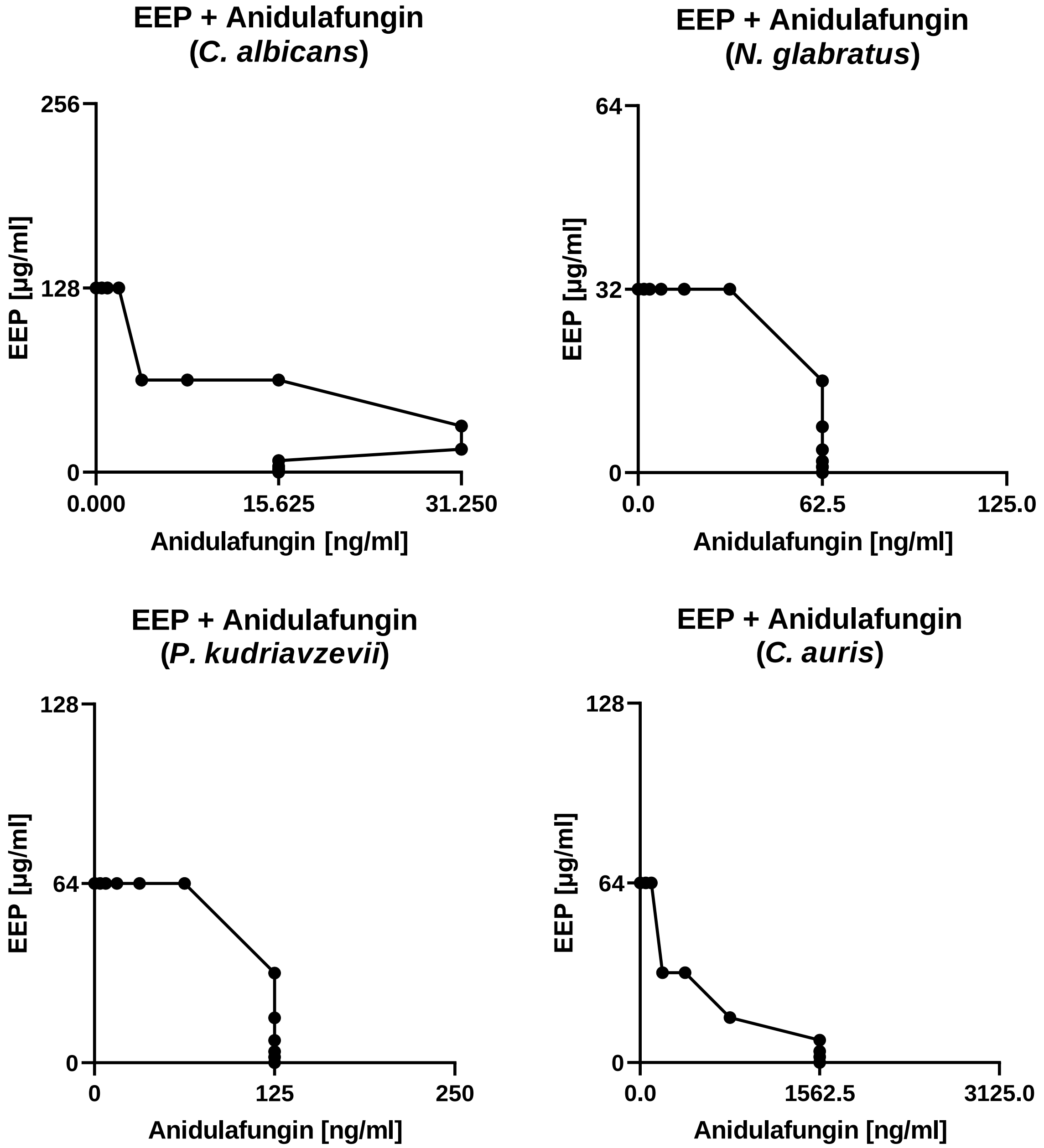


Figure S5 – Isobolograms presenting the dependence between EEP and anidulafungin concentrations regarding $MIC_{90}$ values determined against selected yeast pathogens causing candidiasis.

Table S6 – Summary of estimated fractional inhibitory concentrations and antifungal interaction types between anidulafungin and EEP against selected yeast pathogens causing candidiasis.

| Microorganism | Agent | MIC (ng/mL) | | FIC | ∑FIC | Effect |
|---|---|---|---|---|---|---|
| | | Alone | In combination | | | |
| *Candida albicans* SC 5314 | Anidulafungin | 15.625 | 31.25 | 2 | 2.25 | Indifference |
| | EEP | 128 | 32 | 0.25 | | |
| *Nakaseomyces glabratus* DSM 11226 | Anidulafungin | 62.5 | 62.5 | 1 | 1.03125 | Indifference |
| | EEP | 32 | 1 | 0.03125 | | |
| *Pichia kudriavzevii* DSM 5784 | Anidulafungin | 125 | 125 | 1 | 1.03125 | Indifference |
| | EEP | 64 | 2 | 0.03125 | | |
| *Candida auris* B 8441 | Anidulafungin | 1562.5 | 195.3125 | 0.125 | 0.625 | Additivity |
| | EEP | 64 | 32 | 0.5 | | |

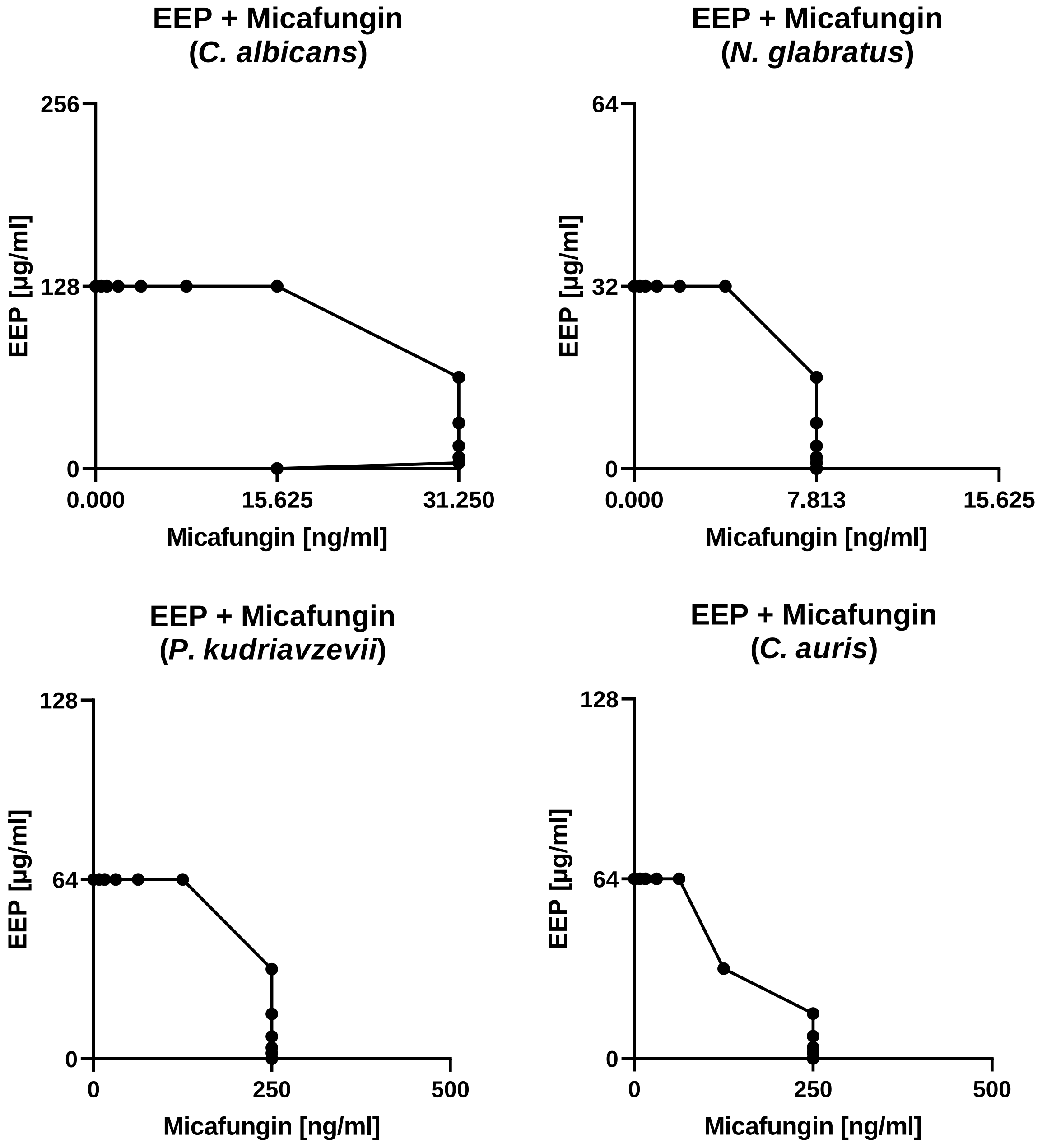


Figure S6 – Isobolograms presenting the dependence between EEP and micafungin concentrations regarding $MIC_{90}$ values determined against selected yeast pathogens causing candidiasis.

Table S7 – Summary of estimated fractional inhibitory concentrations and antifungal interaction types between micafungin and EEP against selected yeast pathogens causing candidiasis.

| Microorganism | Agent | MIC (ng/mL) | | FIC | ∑FIC | Effect |
|---|---|---|---|---|---|---|
| | | Alone | In combination | | | |
| *Candida albicans* SC 5314 | Micafungin | 15.625 | 31.25 | 2 | 2.03125 | Indifference |
| | EEP | 128 | 4 | 0.03125 | | |
| *Nakaseomyces glabratus* DSM 11226 | Micafungin | 7.8125 | 7.8125 | 1 | 1.03125 | Indifference |
| | EEP | 32 | 1 | 0.03125 | | |
| *Pichia kudriavzevii* DSM 5784 | Micafungin | 250 | 250 | 1 | 1.03125 | Indifference |
| | EEP | 64 | 2 | 0.03125 | | |
| *Candida auris* B 8441 | Micafungin | 250 | 125 | 0.5 | 1 | Additivity |
| | EEP | 64 | 32 | 0.5 | | |

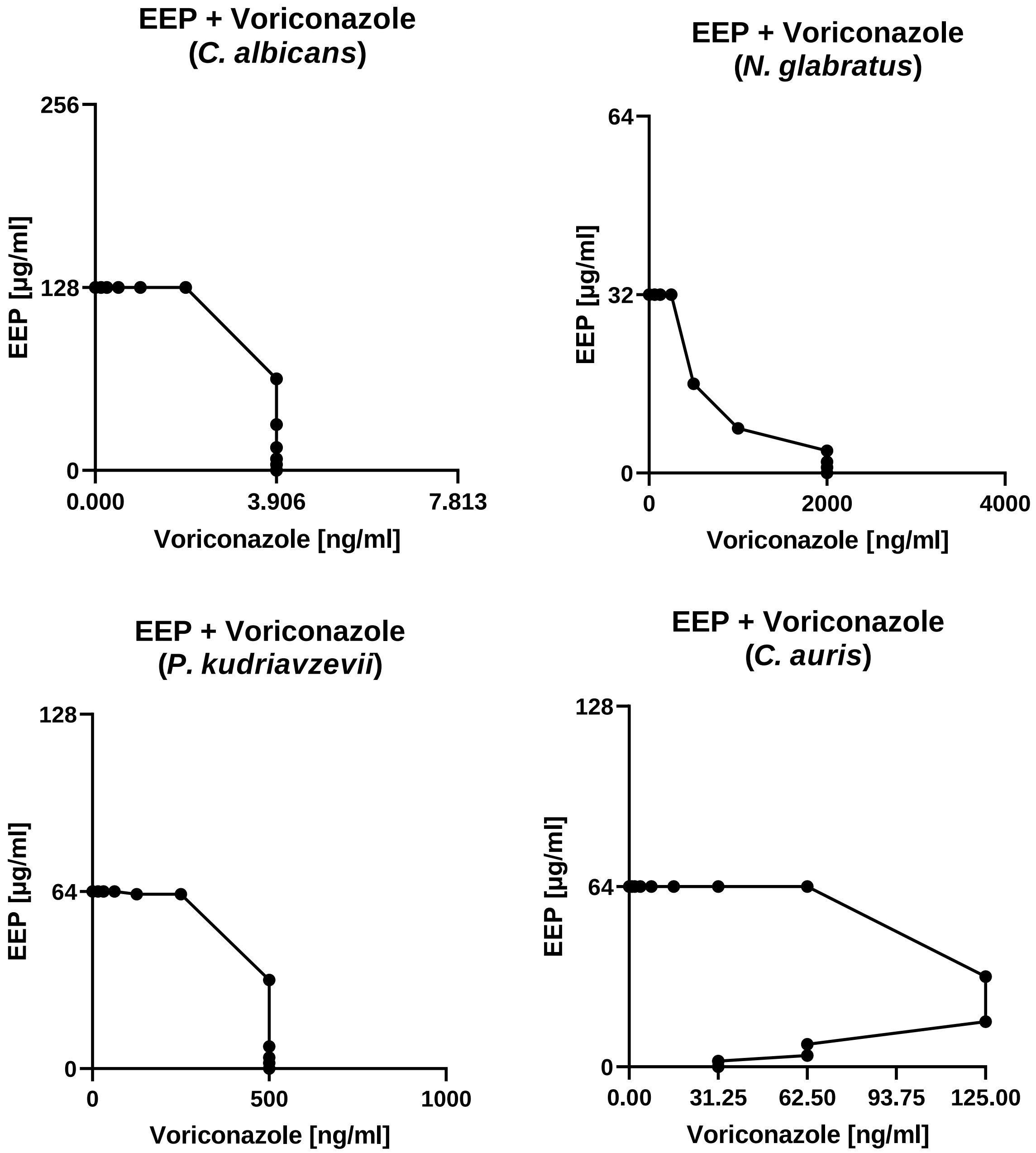


Figure S7 – Isobolograms presenting the dependence between EEP and voriconazole concentrations regarding $MIC_{90}$ values determined against selected yeast pathogens causing candidiasis.

Table S8 – Summary of estimated fractional inhibitory concentrations and antifungal interaction types between voriconazole and EEP against selected yeast pathogens causing candidiasis.

| Microorganism | Agent | MIC (ng/mL) | | FIC | ∑FIC | Effect |
|---|---|---|---|---|---|---|
| | | Alone | In combination | | | |
| *Candida albicans* SC 5314 | Voriconazole | 3.90625 | 3.90625 | 1 | 1.03125 | Indifference |
| | EEP | 128 | 4 | 0.03125 | | |
| *Nakaseomyces glabratus* DSM 11226 | Voriconazole | 2000 | 500 | 0.25 | 0.75 | Additivity |
| | EEP | 32 | 16 | 0.5 | | |
| *Pichia kudriavzevii* DSM 5784 | Voriconazole | 500 | 500 | 1 | 1.03125 | Indifference |
| | EEP | 64 | 2 | 0.03125 | | |
| *Candida auris* B 8441 | Voriconazole | 31.25 | 125 | 4 | 4.25 | Antagonism |
| | EEP | 64 | 16 | 0.25 | | |

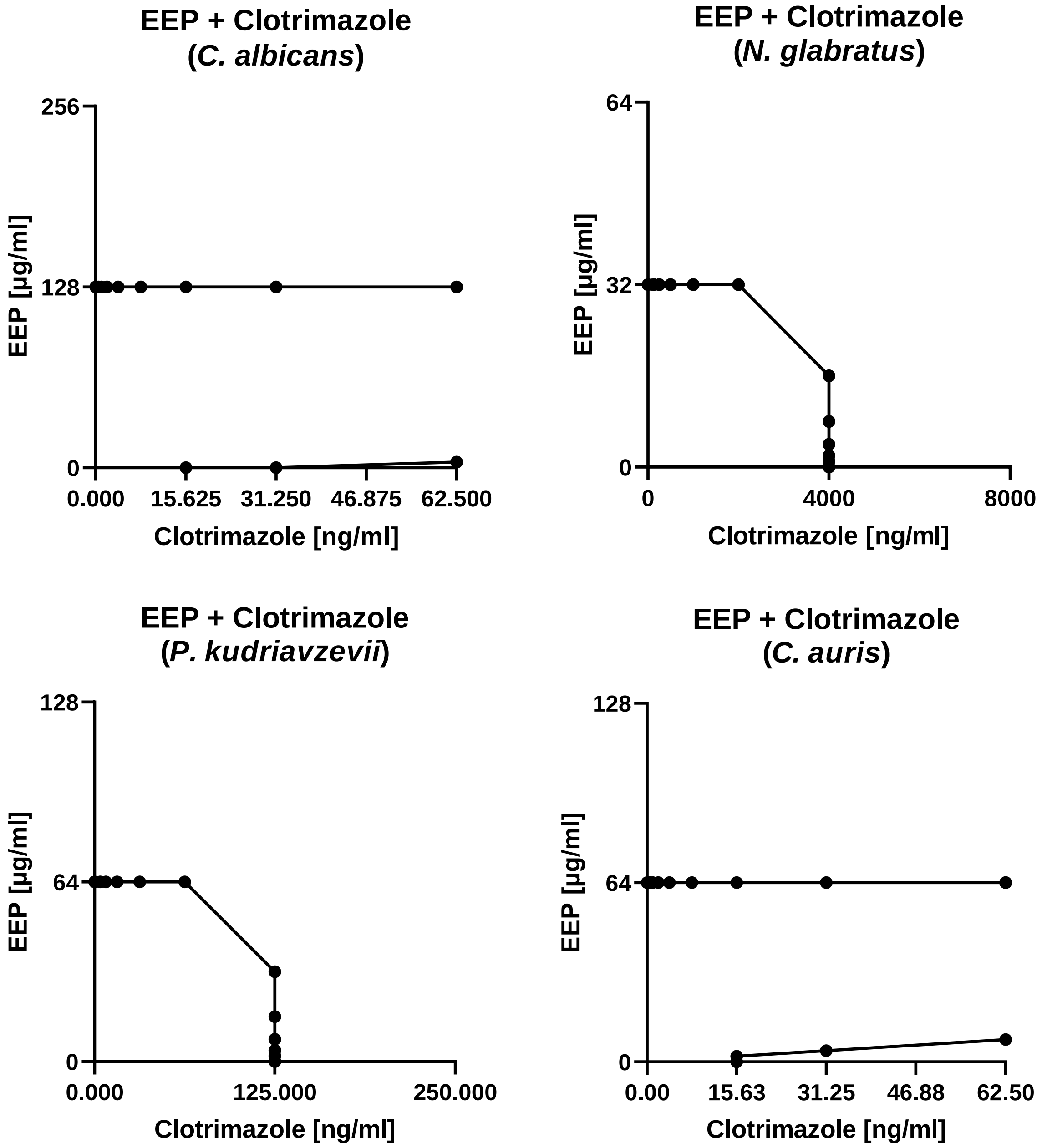


Figure S8 – Isobolograms presenting the dependence between EEP and clotrimazole concentrations regarding $MIC_{90}$ values determined against selected yeast pathogens causing candidiasis.

Table S9 – Summary of estimated fractional inhibitory concentrations and antifungal interaction types between clotrimazole and EEP against selected yeast pathogens causing candidiasis.

| Microorganism | Agent | MIC (ng/mL) | | FIC | ∑FIC | Effect |
|---|---|---|---|---|---|---|
| | | Alone | In combination | | | |
| *Candida albicans* SC 5314 | Clotrimazole | 15.625 | >62.5 | >4 | >4.03125 | Antagonism |
| | EEP | 128 | 4 | 0.03125 | | |
| *Nakaseomyces glabratus* DSM 11226 | Clotrimazole | 4000 | 4000 | 1 | 1.03125 | Indifference |
| | EEP | 32 | 1 | 0.03125 | | |
| *Pichia kudriavzevii* DSM 5784 | Clotrimazole | 125 | 125 | 1 | 1.03125 | Indifference |
| | EEP | 64 | 2 | 0.03125 | | |
| *Candida auris* B 8441 | Clotrimazole | 15.625 | >125 | >4 | >4.0625 | Antagonism |
| | EEP | 64 | 4 | 0.0625 | | |

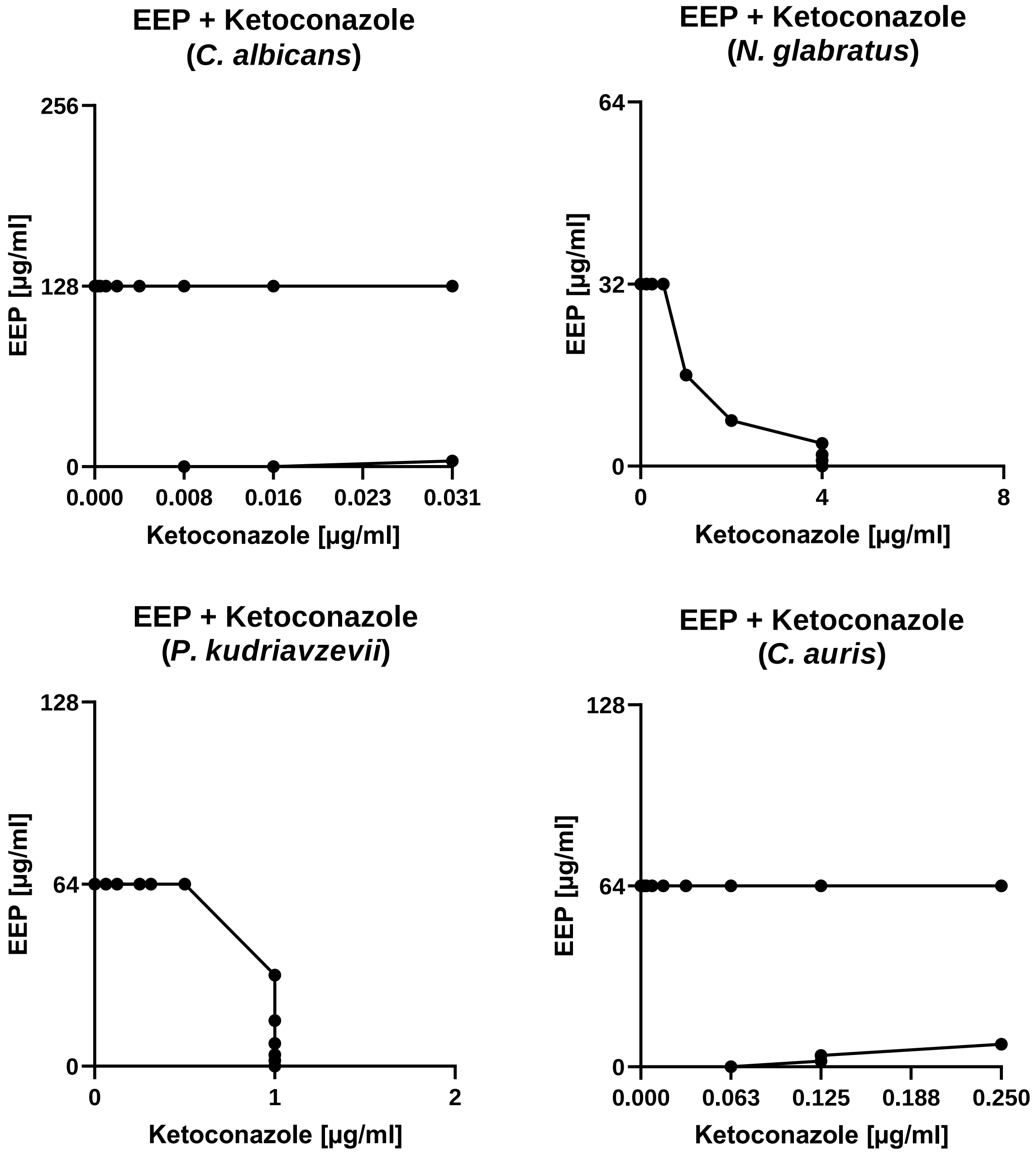


Figure S9 – Isobolograms presenting the dependence between EEP and ketoconazole concentrations regarding $MIC_{90}$ values determined against selected yeast pathogens causing candidiasis.

Table S10 – Summary of estimated fractional inhibitory concentrations and antifungal interaction types between ketoconazole and EEP against selected yeast pathogens causing candidiasis.

| Microorganism | Agent | MIC (µg/mL) | | FIC | ∑FIC | Effect |
|---|---|---|---|---|---|---|
| | | Alone | In combination | | | |
| *Candida albicans* SC 5314 | Ketoconazole | 0.0078125 | >0.03125 | >4 | >4.03125 | Antagonism |
| | EEP | 128 | 4 | 0.03125 | | |
| *Nakaseomyces glabratus* DSM 11226 | Ketoconazole | 4 | 4 | 1 | 1.03125 | Indifference |
| | EEP | 32 | 1 | 0.03125 | | |
| *Pichia kudriavzevii* DSM 5784 | Ketoconazole | 1 | 1 | 1 | 1.03125 | Indifference |
| | EEP | 64 | 2 | 0.03125 | | |
| *Candida auris* B 8441 | Ketoconazole | 0.0625 | >0.25 | >4 | >4.0625 | Antagonism |
| | EEP | 64 | 4 | 0.0625 | | |

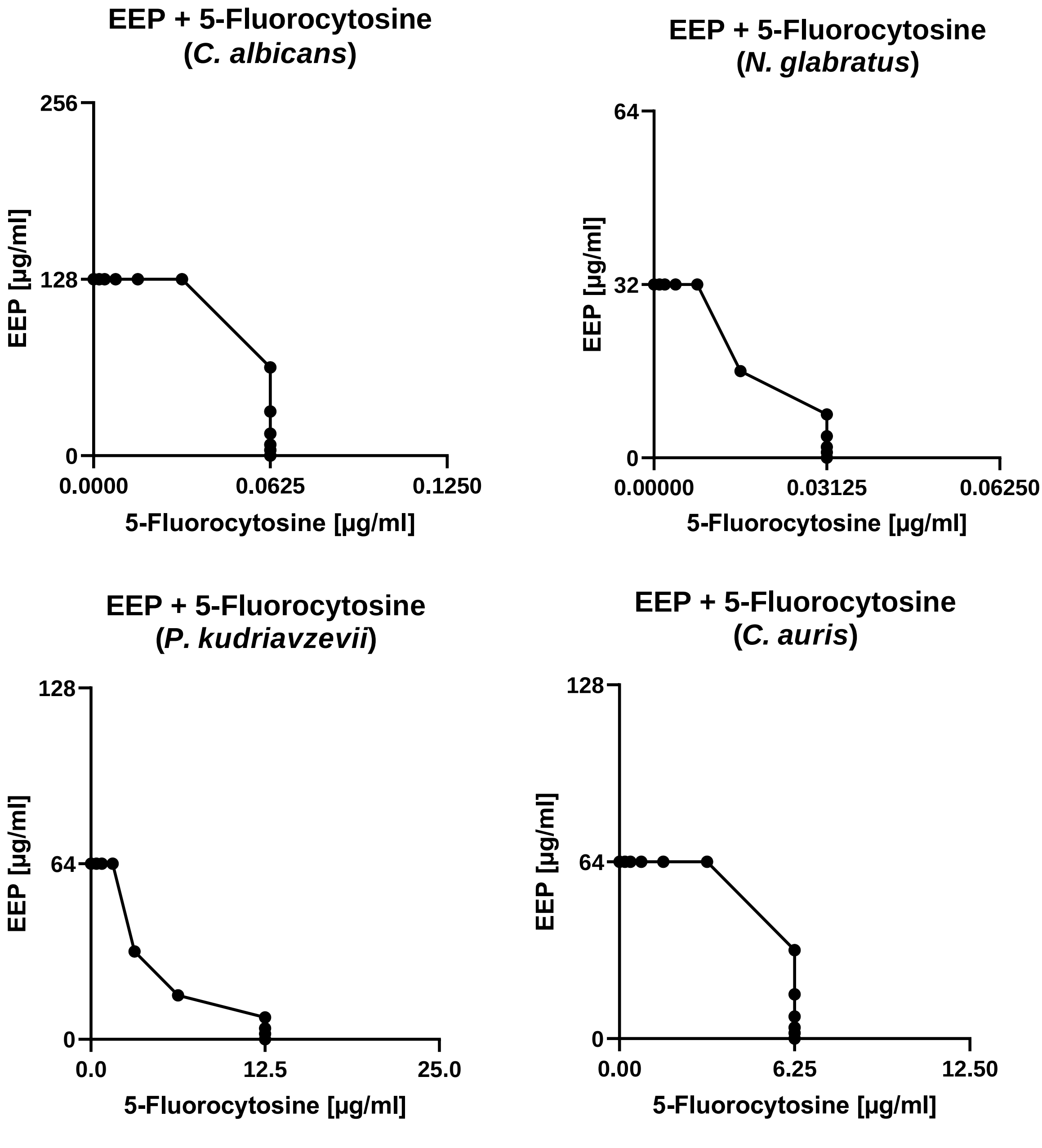


Figure S10 – Isobolograms presenting the dependence between EEP and 5-fluorocytosine concentrations regarding $MIC_{90}$ values determined against selected yeast pathogens causing candidiasis.

Table S11 – Summary of estimated fractional inhibitory concentrations and antifungal interaction types between 5-fluorocytosine and EEP against selected yeast pathogens causing candidiasis.

| Microorganism | Agent | MIC (µg/mL) | | FIC | ∑FIC | Effect |
|---|---|---|---|---|---|---|
| | | Alone | In combination | | | |
| *Candida albicans* SC 5314 | 5-Fluorocytosine | 0.0625 | 0.0625 | 1 | 1.03125 | Indifference |
| | EEP | 128 | 4 | 0.03125 | | |
| *Nakaseomyces glabratus* DSM 11226 | 5-Fluorocytosine | 0.03125 | 0.015625 | 0.5 | 1 | Additivity |
| | EEP | 32 | 16 | 0.5 | | |
| *Pichia kudriavzevii* DSM 5784 | 5-Fluorocytosine | 12.5 | 3.125 | 0.25 | 0.75 | Additivity |
| | EEP | 64 | 32 | 0.5 | | |
| *Candida auris* B 8441 | 5-Fluorocytosine | 6.25 | 6.25 | 1 | 1.03125 | Indifference |
| | EEP | 64 | 2 | 0.03125 | | |

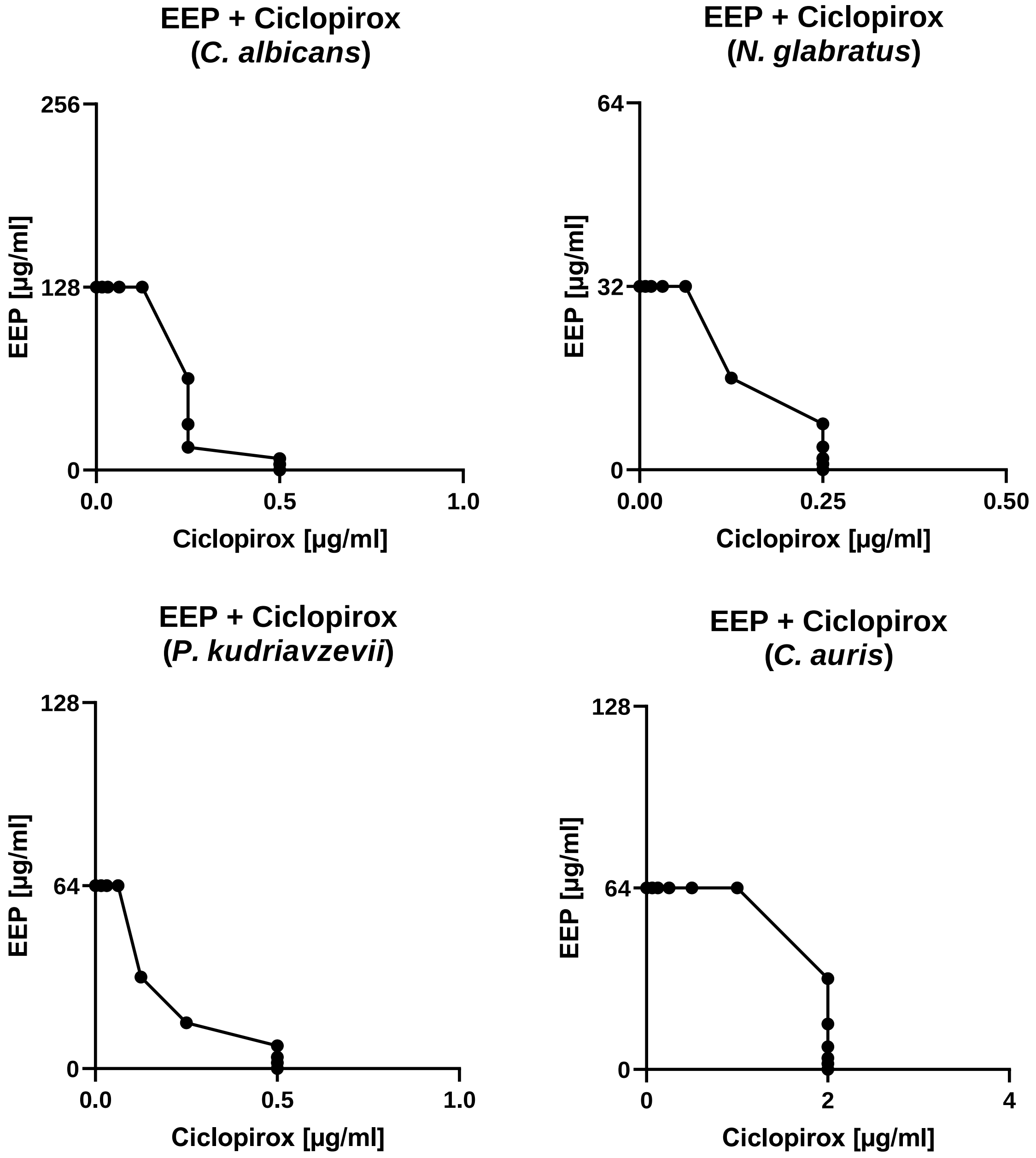


Figure S11 – Isobolograms presenting the dependence between EEP and ciclopirox concentrations regarding $MIC_{90}$ values determined against selected yeast pathogens causing candidiasis.

Table S12 – Summary of estimated fractional inhibitory concentrations and antifungal interaction types between ciclopirox and EEP against selected yeast pathogens causing candidiasis.

| Microorganism | Agent | MIC (µg/mL) | | FIC | ∑FIC | Effect |
|---|---|---|---|---|---|---|
| | | Alone | In combination | | | |
| *Candida albicans* SC 5314 | Ciclopirox | 0.5 | 0.25 | 0.5 | 0.625 | Additivity |
| | EEP | 128 | 16 | 0.125 | | |
| *Nakaseomyces glabratus* DSM 11226 | Ciclopirox | 0.25 | 0.125 | 0.5 | 1 | Additivity |
| | EEP | 32 | 16 | 0.5 | | |
| *Pichia kudriavzevii* DSM 5784 | Ciclopirox | 0.5 | 0.125 | 0.25 | 0.75 | Additivity |
| | EEP | 64 | 32 | 0.5 | | |
| *Candida auris* B 8441 | Ciclopirox | 2 | 2 | 1 | 1.03125 | Indifference |
| | EEP | 64 | 2 | 0.03125 | | |

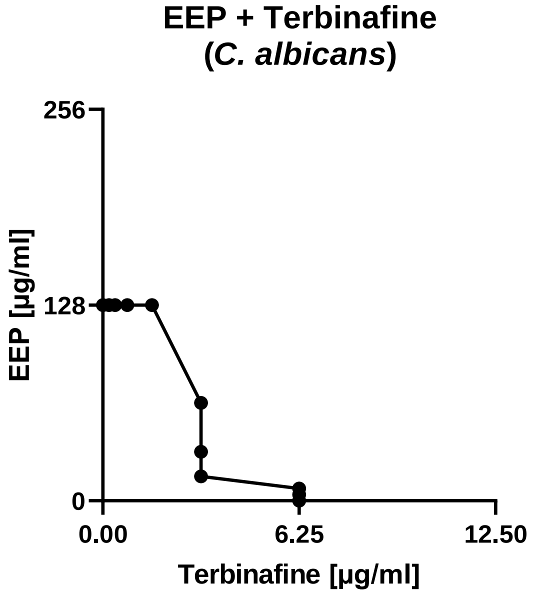


Figure S12 – Isobologram presenting the dependence between EEP and terbinafine concentrations regarding $MIC_{90}$ values determined against *Candida albicans*.

Table S13 – Estimated fractional inhibitory concentration and antifungal interaction type between terbinafine and EEP against *Candida albicans* SC 5314.

| Microorganism | Agent | MIC (µg/mL) | | FIC | ∑FIC | Effect |
|---|---|---|---|---|---|---|
| | | Alone | In combination | | | |
| *Candida albicans* SC 5314 | Terbinafine | 6.25 | 3.125 | 0.5 | 0.625 | Additivity |
| | EEP | 128 | 16 | 0.125 | | |

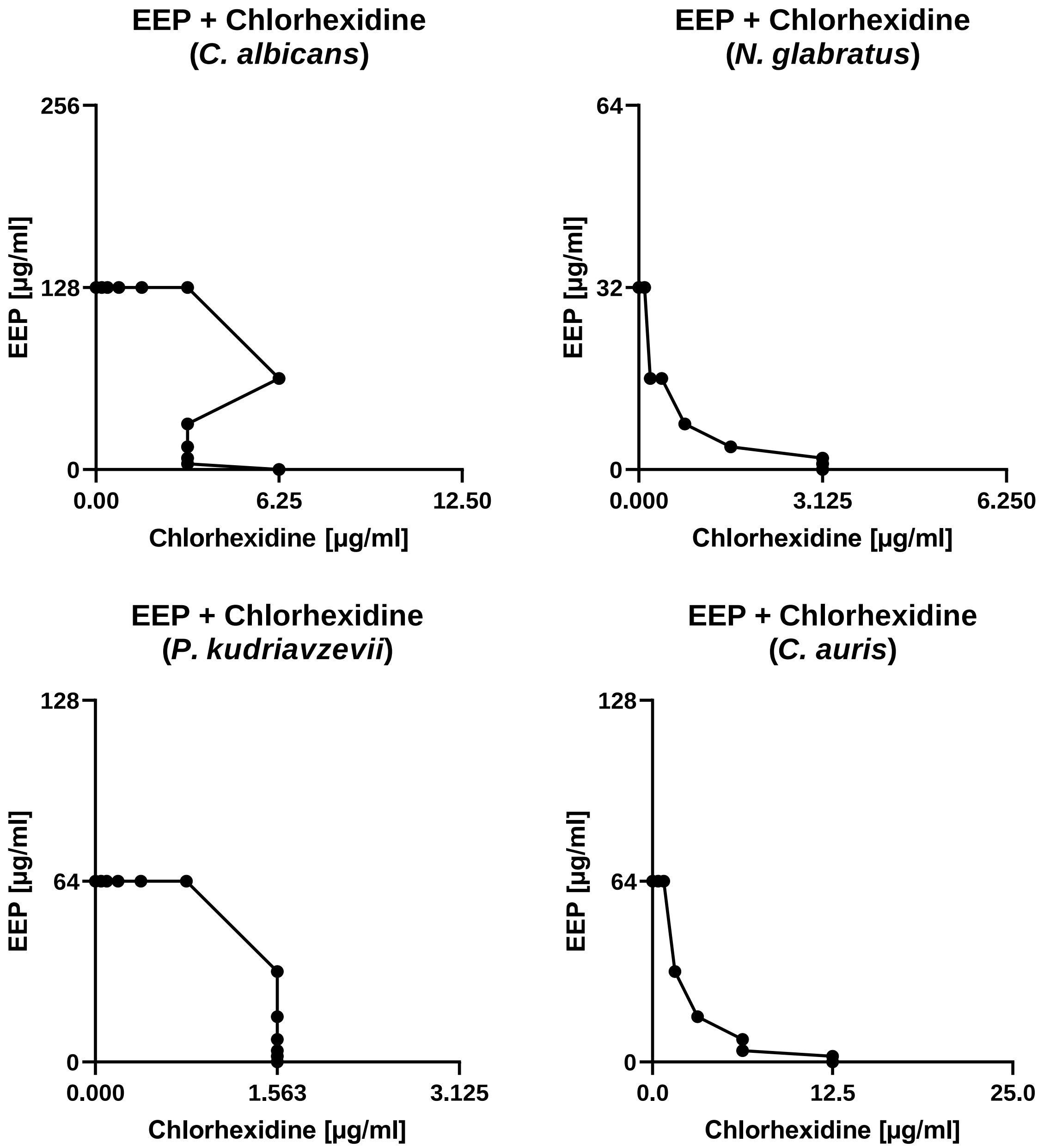


Figure S13 – Isobolograms presenting the dependence between EEP and chlorhexidine concentrations regarding $MIC_{90}$ values determined against selected yeast pathogens causing candidiasis.

Table S14 – Summary of estimated fractional inhibitory concentrations and antifungal interaction types between chlorhexidine and EEP against selected yeast pathogens causing candidiasis.

| Microorganism | Agent | MIC (µg/mL) | | FIC | ∑FIC | Effect |
|---|---|---|---|---|---|---|
| | | Alone | In combination | | | |
| *Candida albicans* SC 5314 | Chlorhexidine | 6.25 | 3.125 | 0.5 | 0.53125 | Additivity |
| | EEP | 128 | 4 | 0.03125 | | |
| *Nakaseomyces glabratus* DSM 11226 | Chlorhexidine | 3.125 | 0.78125 | 0.25 | 0.5 | Synergy |
| | EEP | 32 | 8 | 0.25 | | |
| *Pichia kudriavzevii* DSM 5784 | Chlorhexidine | 1.5625 | 1.5625 | 1 | 1.03125 | Indifference |
| | EEP | 64 | 2 | 0.03125 | | |
| *Candida auris* B 8441 | Chlorhexidine | 12.5 | 3.125 | 0.25 | 0.5 | Synergy |
| | EEP | 64 | 16 | 0.25 | | |

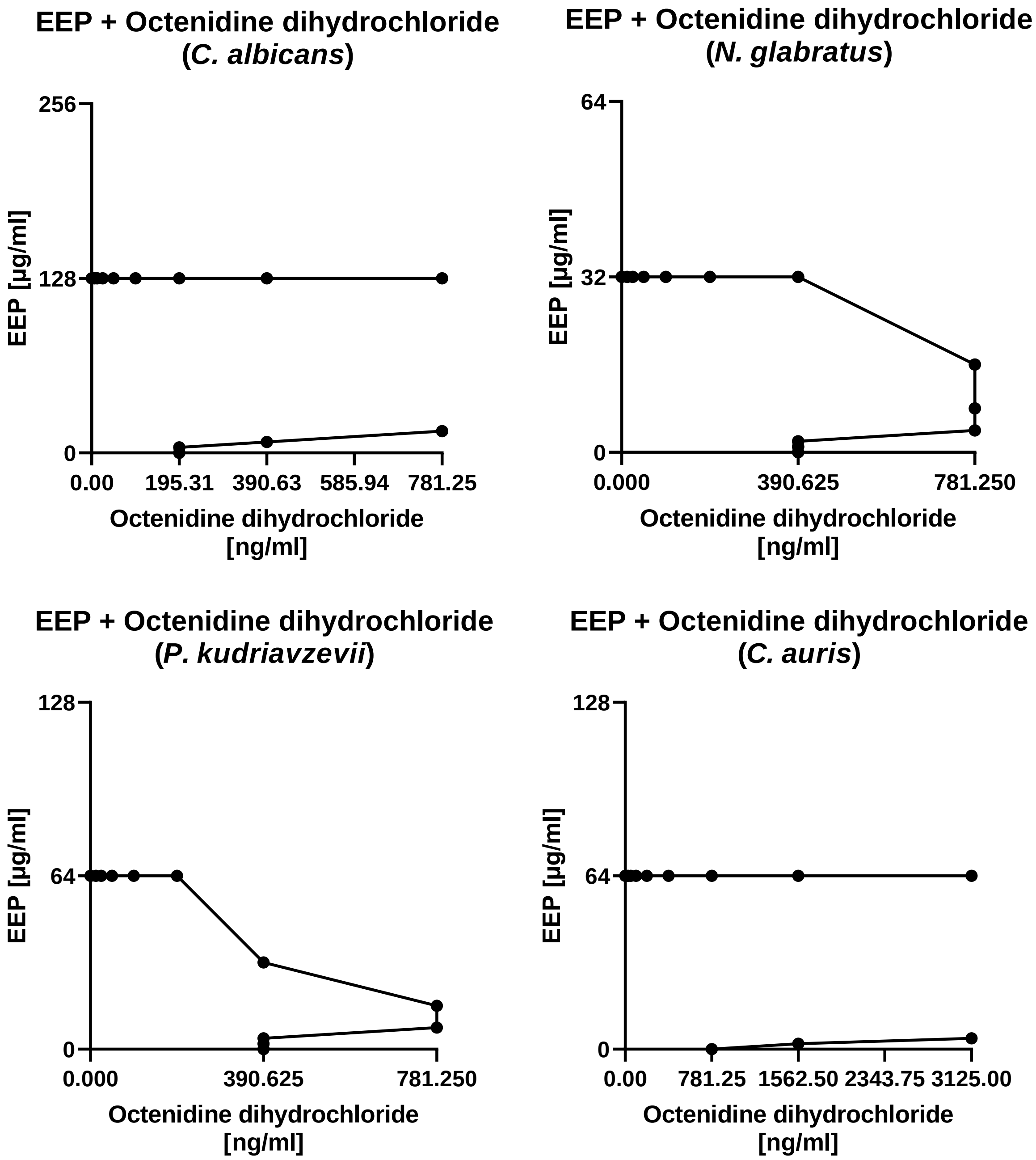


Figure S14 – Isobolograms presenting the dependence between EEP and octenidine dihydrochloride concentrations regarding $MIC_{90}$ values determined against selected yeast pathogens causing candidiasis.

Table S15 – Summary of estimated fractional inhibitory concentrations and antifungal interaction types between octenidine dihydrochloride and EEP against selected yeast pathogens causing candidiasis.

| Microorganism | Agent | MIC (ng/mL) | | FIC | ∑FIC | Effect |
|---|---|---|---|---|---|---|
| | | Alone | In combination | | | |
| *Candida albicans* SC 5314 | Octenidine dihydrochloride | 195.31 | >781.24 | >4 | >4.125 | Antagonism |
| | EEP | 128 | 16 | 0.125 | | |
| *Nakaseomyces glabratus* DSM 11226 | Octenidine dihydrochloride | 390.625 | 781.25 | 2 | 2.125 | Indifference |
| | EEP | 32 | 4 | 0.125 | | |
| *Pichia kudriavzevii* DSM 5784 | Octenidine dihydrochloride | 390.625 | 781.25 | 2 | 2.125 | Indifference |
| | EEP | 64 | 8 | 0.125 | | |
| *Candida auris* B 8441 | Octenidine dihydrochloride | 781.25 | >3125 | >4 | >4.03125 | Antagonism |
| | EEP | 64 | 2 | 0.03125 | | |

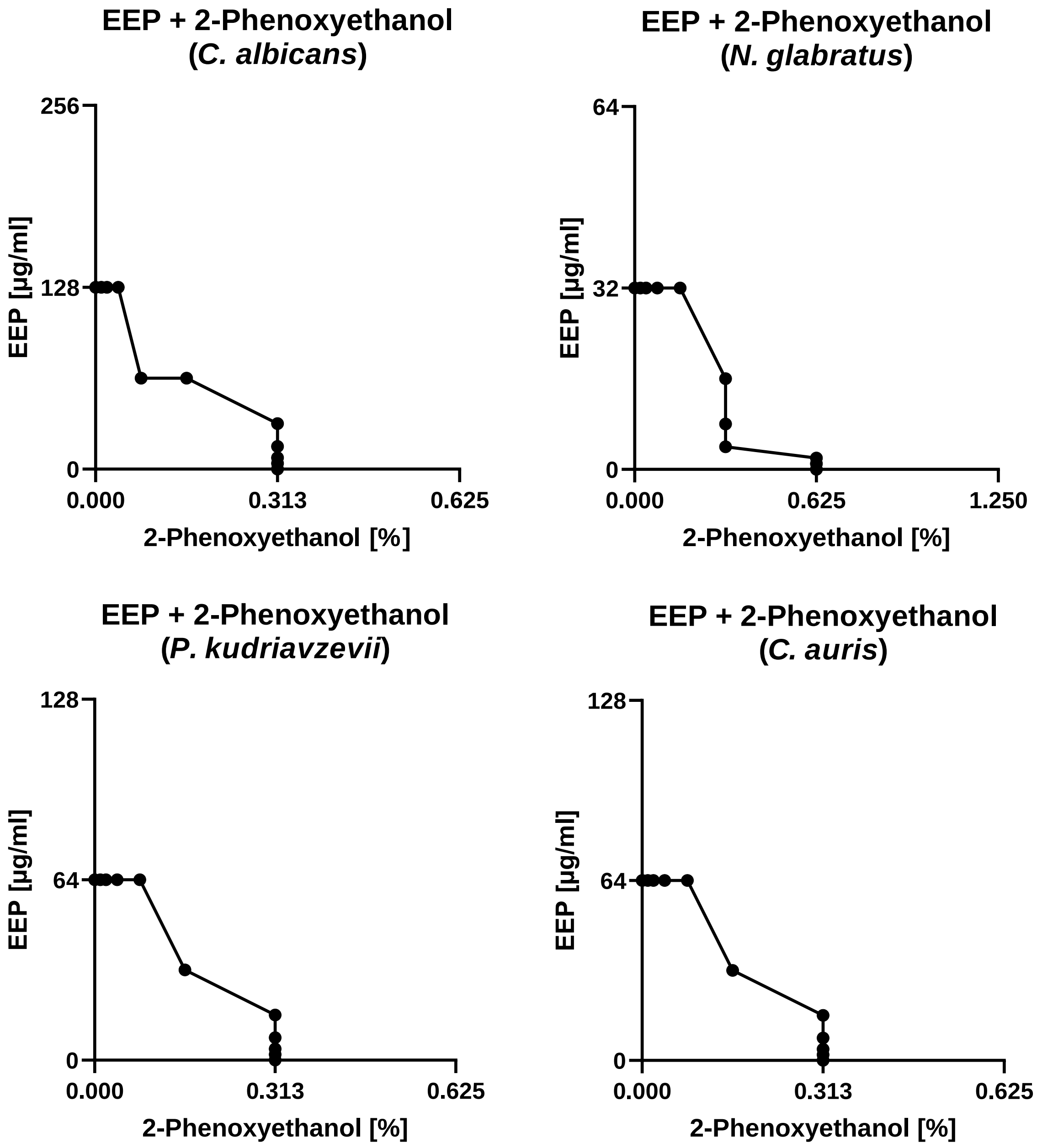


Figure S15 – Isobolograms presenting the dependence between EEP and 2-phenoxyethanol concentrations regarding $MIC_{90}$ values determined against selected yeast pathogens causing candidiasis.

Table S16 – Summary of estimated fractional inhibitory concentrations and antifungal interaction types between 2-phenoxyethanol and EEP against selected yeast pathogens causing candidiasis.

| Microorganism | Agent | MIC (% \| µg/mL) | | FIC | ∑FIC | Effect |
|---|---|---|---|---|---|---|
| | | Alone | In combination | | | |
| *Candida albicans* SC 5314 | 2-Phenoxyethanol | 0.3125 | 0.078125 | 0.25 | 0.75 | Additivity |
| | EEP | 128 | 64 | 0.5 | | |
| *Nakaseomyces glabratus* DSM 11226 | 2-Phenoxyethanol | 0.625 | 0.3125 | 0.5 | 0.625 | Additivity |
| | EEP | 32 | 4 | 0.125 | | |
| *Pichia kudriavzevii* DSM 5784 | 2-Phenoxyethanol | 0.3125 | 0.3125 | 1 | 1.03125 | Indifference |
| | EEP | 64 | 2 | 0.03125 | | |
| *Candida auris* B 8441 | 2-Phenoxyethanol | 0.3125 | 0.15625 | 0.5 | 1 | Additivity |
| | EEP | 64 | 32 | 0.5 | | |

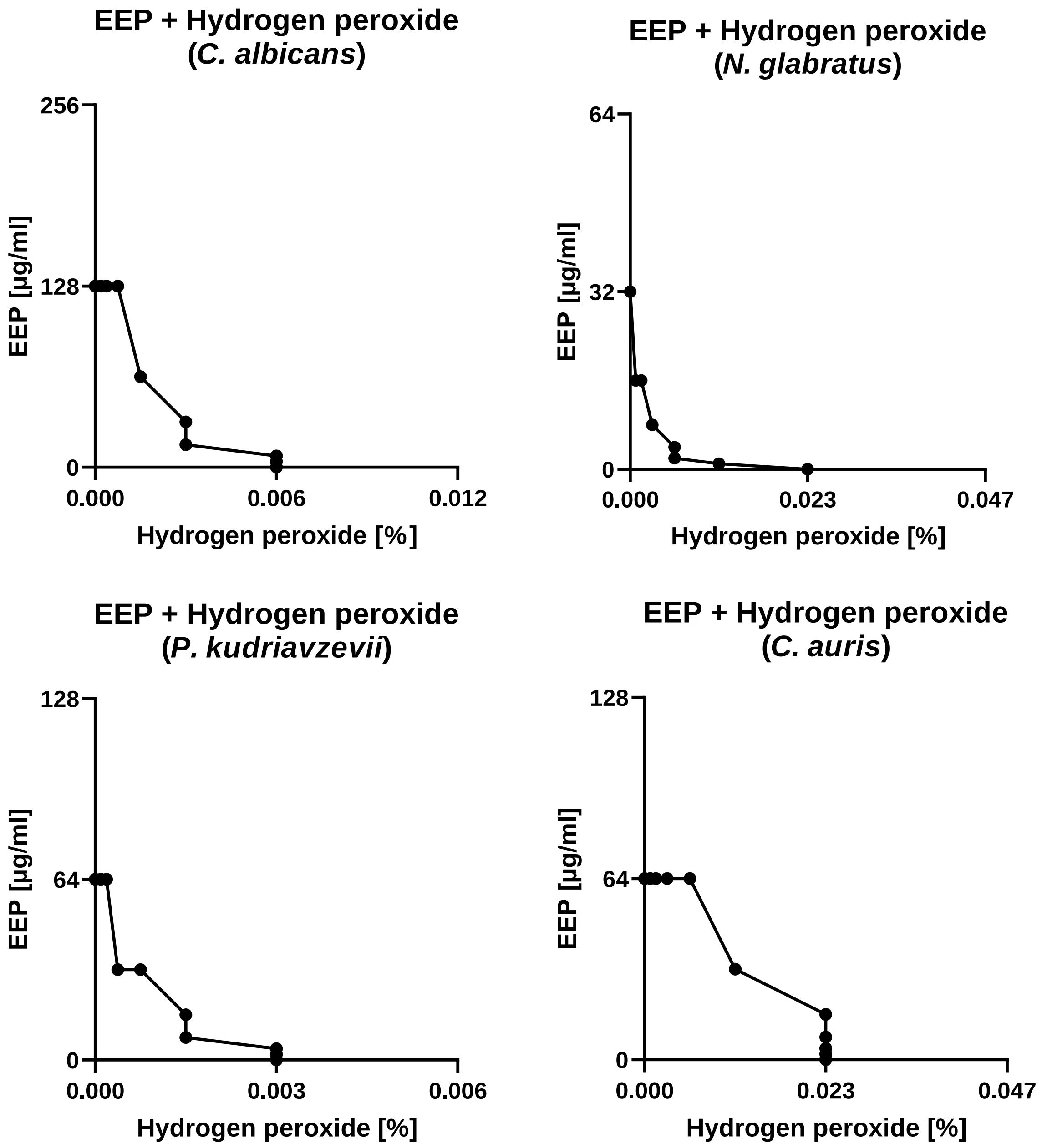


Figure S16 – Isobolograms presenting the dependence between EEP and hydrogen peroxide concentrations regarding $MIC_{90}$ values determined against selected yeast pathogens causing candidiasis.

Table S17 – Summary of estimated fractional inhibitory concentrations and antifungal interaction types between hydrogen peroxide and EEP against selected yeast pathogens causing candidiasis.

| Microorganism | Agent | MIC (% \| µg/mL) | | FIC | ∑FIC | Effect |
|---|---|---|---|---|---|---|
| | | Alone | In combination | | | |
| *Candida albicans* SC 5314 | Hydrogen peroxide | 0.005859375 | 0.002929687 | 0.5 | 0.625 | Additivity |
| | EEP | 128 | 16 | 0.125 | | |
| *Nakaseomyces glabratus* DSM 11226 | Hydrogen peroxide | 0.0234375 | 0.005859375 | 0.25 | 0.3125 | Synergy |
| | EEP | 32 | 2 | 0.0625 | | |
| *Pichia kudriavzevii* DSM 5784 | Hydrogen peroxide | 0.002929687 | 0.000366211 | 0.125 | 0.625 | Additivity |
| | EEP | 64 | 32 | 0.5 | | |
| *Candida auris* B 8441 | Hydrogen peroxide | 0.0234375 | 0.01171875 | 0.5 | 1 | Additivity |
| | EEP | 64 | 32 | 0.5 | | |

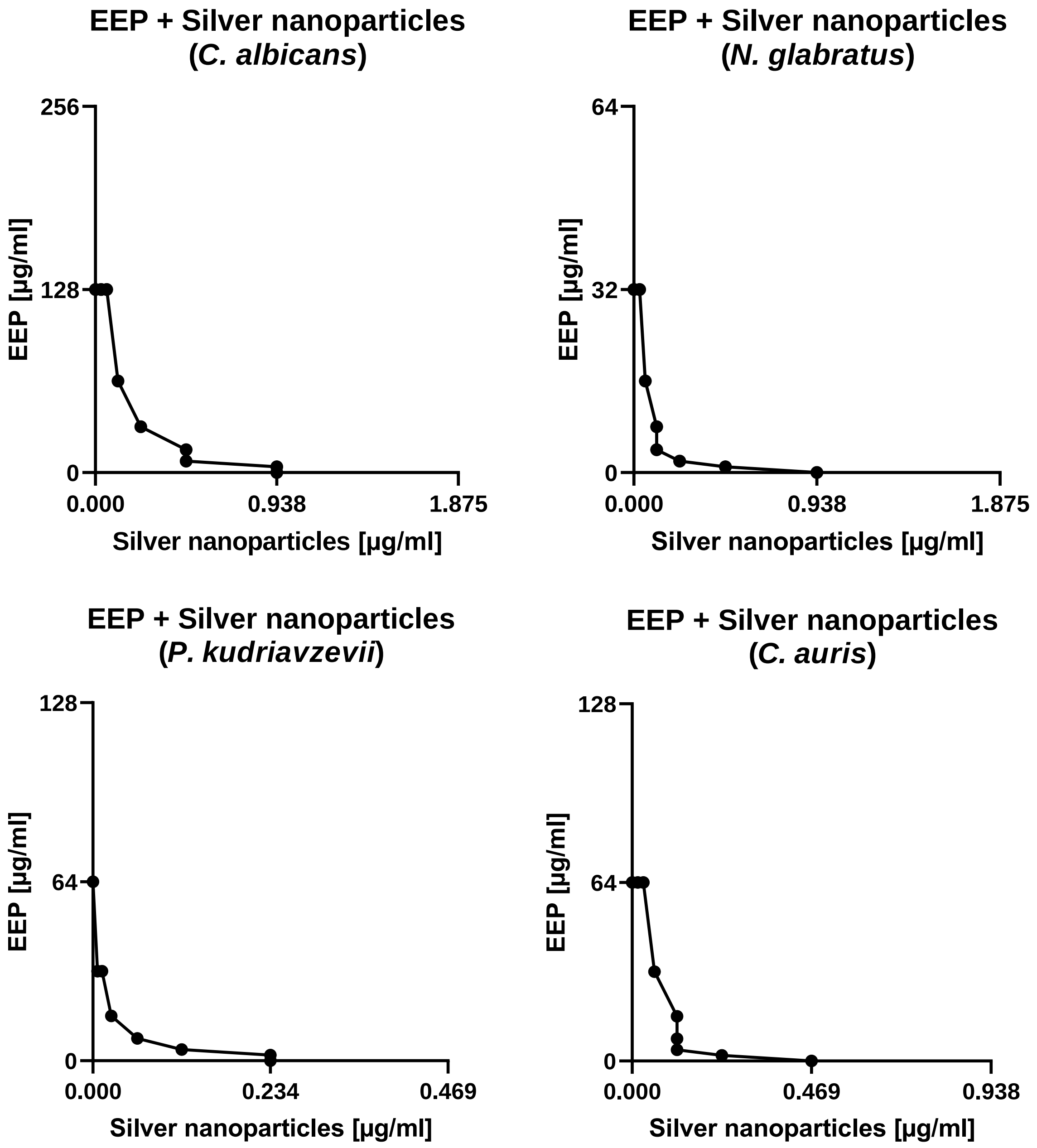


Figure S17 – Isobolograms presenting the dependence between EEP and silver nanoparticles concentrations regarding $MIC_{90}$ values determined against selected yeast pathogens causing candidiasis.

Table S18 – Summary of estimated fractional inhibitory concentrations and antifungal interaction types between silver nanoparticles and EEP against selected yeast pathogens causing candidiasis.

| Microorganism | Agent | MIC (µg/mL) | | FIC | ∑FIC | Effect |
|---|---|---|---|---|---|---|
| | | Alone | In combination | | | |
| *Candida albicans* SC 5314 | Silver nanoparticles | 0.9375 | 0.234375 | 0.25 | 0.5 | Synergy |
| | EEP | 128 | 64 | 0.25 | | |
| *Nakaseomyces glabratus* DSM 11226 | Silver nanoparticles | 0.9375 | 0.234375 | 0.25 | 0.3125 | Synergy |
| | EEP | 32 | 2 | 0.0625 | | |
| *Pichia kudriavzevii* DSM 5784 | Silver nanoparticles | 0.234375 | 0.029296875 | 0.125 | 0.375 | Synergy |
| | EEP | 64 | 8 | 0.25 | | |
| *Candida auris* B 8441 | Silver nanoparticles | 0.46875 | 0.1171875 | 0.25 | 0.3125 | Synergy |
| | EEP | 64 | 4 | 0.0625 | | |

Table S19 – Summary of estimated fractional inhibitory concentrations and antifungal interaction types between fluconazole and EEP against selected yeast pathogens causing candidiasis.

| Microorganism | Agent | MIC (µg/mL) | | FIC | ∑FIC | Effect |
|---|---|---|---|---|---|---|
| | | Alone | In combination | | | |
| *Candida albicans* SC 5314 | Fluconazole | 0.125 | 0.125 | 1 | 1.03125 | Indifference |
| | EEP | 128 | 4 | 0.03125 | | |
| *Nakaseomyces glabratus* DSM 11226 | Fluconazole | 32 | 8 | 0.25 | 0.75 | Additivity |
| | EEP | 32 | 16 | 0.5 | | |
| *Pichia kudriavzevii* DSM 5784 | Fluconazole | 64 | 64 | 1 | 1.03125 | Indifference |
| | EEP | 64 | 2 | 0.03125 | | |
| *Candida auris* B 8441 | Fluconazole | 2 | >8 | >4 | >4.03125 | Antagonism |
| | EEP | 64 | 2 | 0.03125 | | |

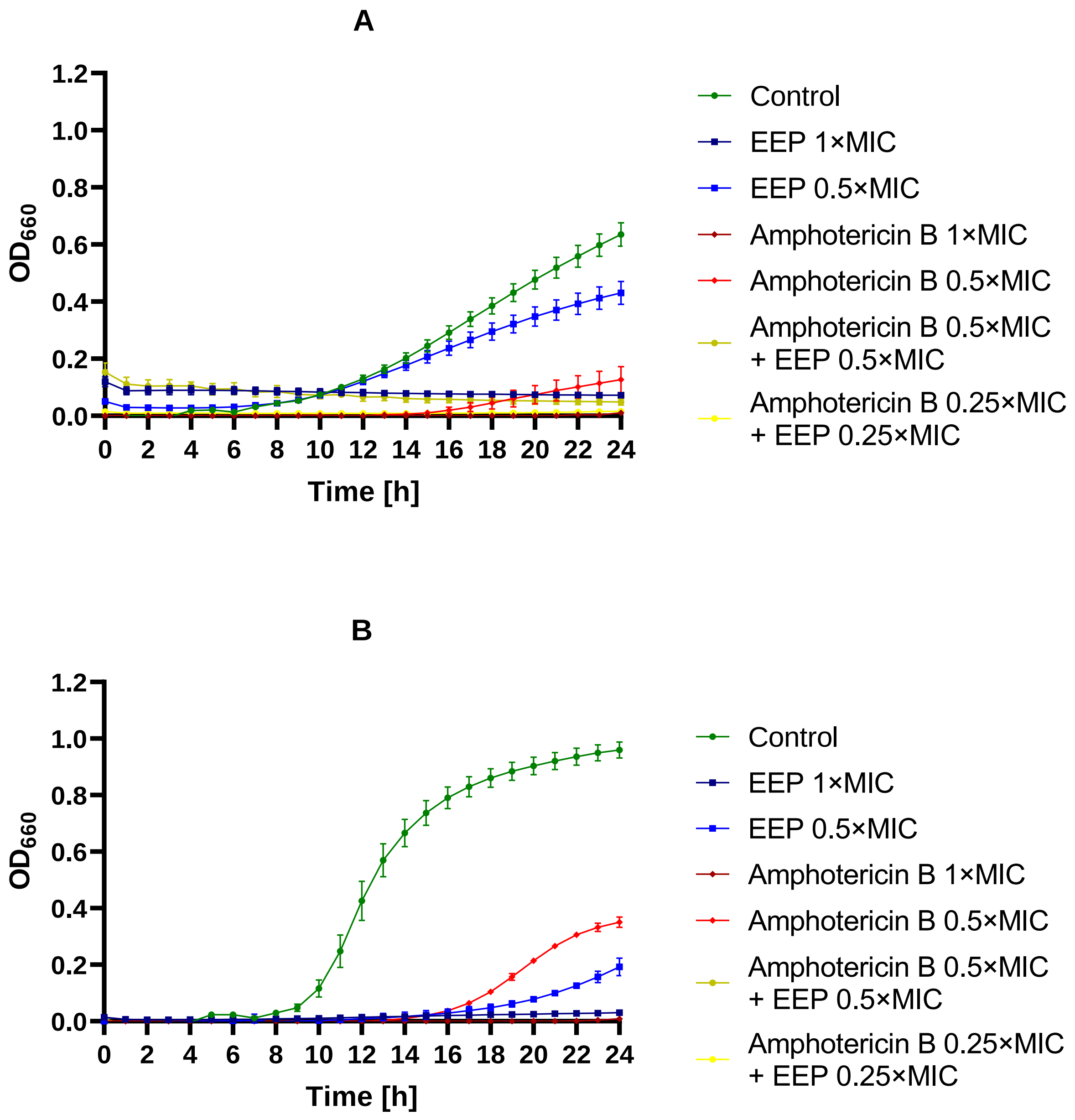
A
1.2
1.0
0.8
0.6
0.4
0.2
0.0
$OD_{660}$
0 2 4 6 8 10 12 14 16 18 20 22 24
Time [h]
Control
EEP 1×MIC
EEP 0.5×MIC
Amphotericin B 1×MIC
Amphotericin B 0.5×MIC
Amphotericin B 0.5×MIC + EEP 0.5×MIC
Amphotericin B 0.25×MIC + EEP 0.25×MIC
B
1.2
1.0
0.8
0.6
0.4
0.2
0.0
$OD_{660}$
0 2 4 6 8 10 12 14 16 18 20 22 24
Time [h]
Control
EEP 1×MIC
EEP 0.5×MIC
Amphotericin B 1×MIC
Amphotericin B 0.5×MIC
Amphotericin B 0.5×MIC + EEP 0.5×MIC
Amphotericin B 0.25×MIC + EEP 0.25×MIC

**C**

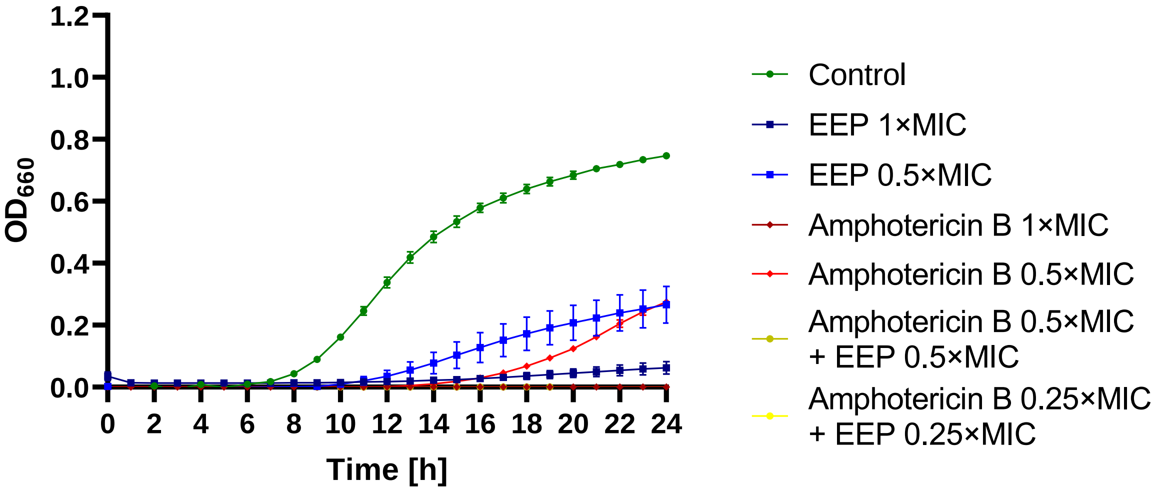


**D**

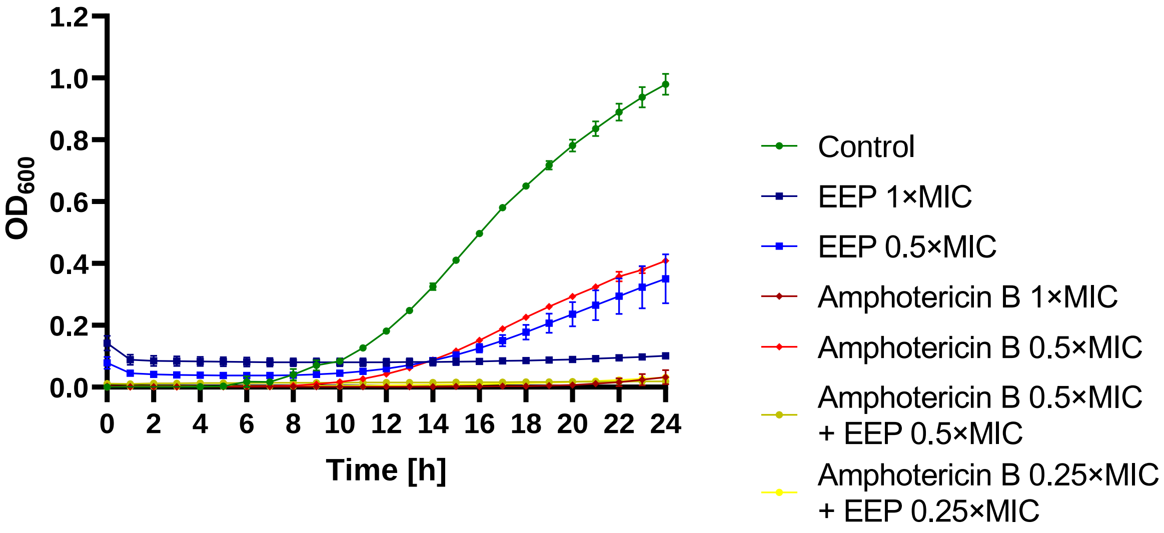


Figure S18 – Antifungal activity of EEP, amphotericin B, and their mixtures – growth curves (monitoring of the growth kinetics of the yeast cells measured as turbidity of the cultures at wavelength 660 nm or 600 nm for *C. auris*) against A – *C. albicans*, B – *N. glabratus*, C – *P. kudriavzevii*, and D – *C. auris*.

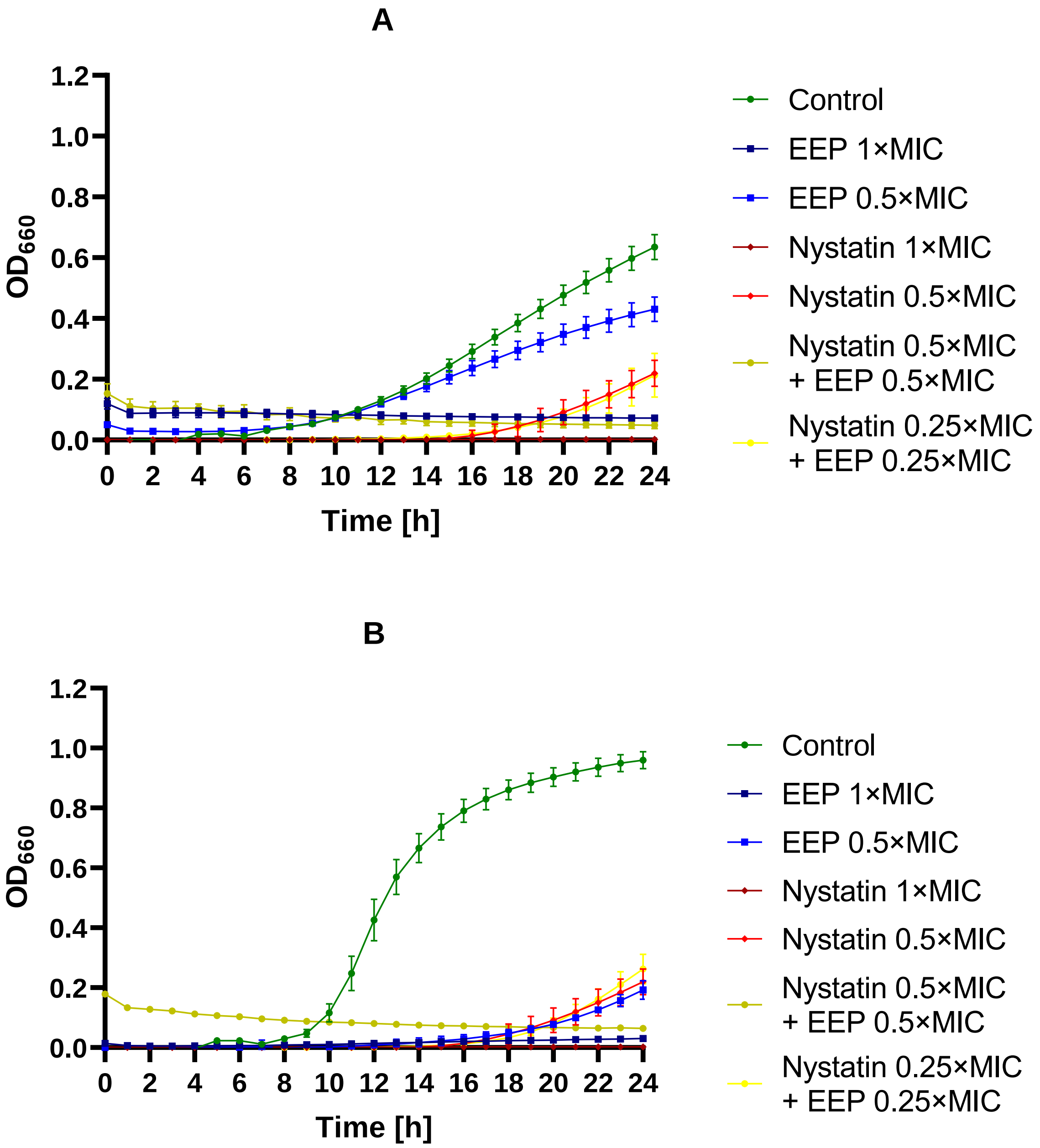

A
OD660
Time [h]
Control
EEP 1×MIC
EEP 0.5×MIC
Nystatin 1×MIC
Nystatin 0.5×MIC
Nystatin 0.5×MIC + EEP 0.5×MIC
Nystatin 0.25×MIC + EEP 0.25×MIC
B
OD660
Time [h]
Control
EEP 1×MIC
EEP 0.5×MIC
Nystatin 1×MIC
Nystatin 0.5×MIC
Nystatin 0.5×MIC + EEP 0.5×MIC
Nystatin 0.25×MIC + EEP 0.25×MIC

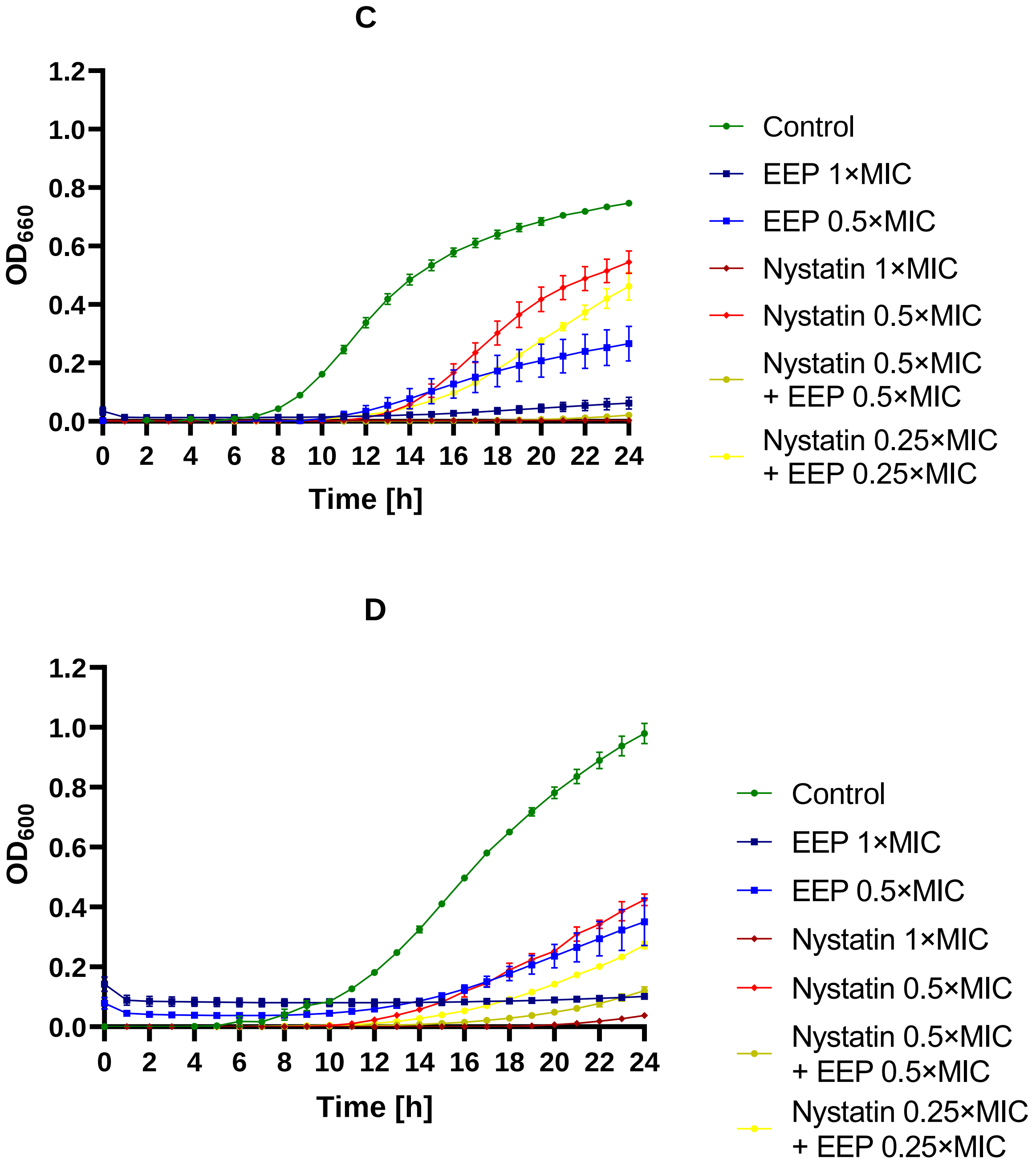


Figure S19 – Antifungal activity of EEP, nystatin, and their mixtures – growth curves (monitoring of the growth kinetics of the yeast cells measured as turbidity of the cultures at wavelength 660 nm or 600 nm for *C. auris*) against A – *C. albicans*, B – *N. glabratus*, C – *P. kudriavzevii*, and D – *C. auris*.

A

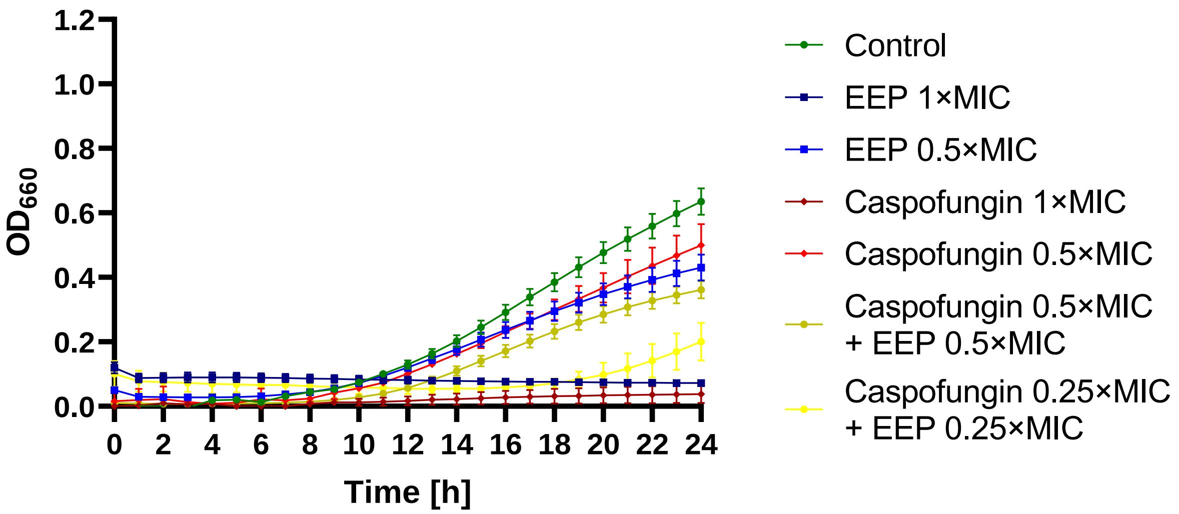

1.2
1.0
0.8
0.6
0.4
0.2
0.0
$OD_{660}$
0
2
4
6
8
10
12
14
16
18
20
22
24
Time [h]
Control
EEP 1×MIC
EEP 0.5×MIC
Caspofungin 1×MIC
Caspofungin 0.5×MIC
Caspofungin 0.5×MIC + EEP 0.5×MIC
Caspofungin 0.25×MIC + EEP 0.25×MIC


B

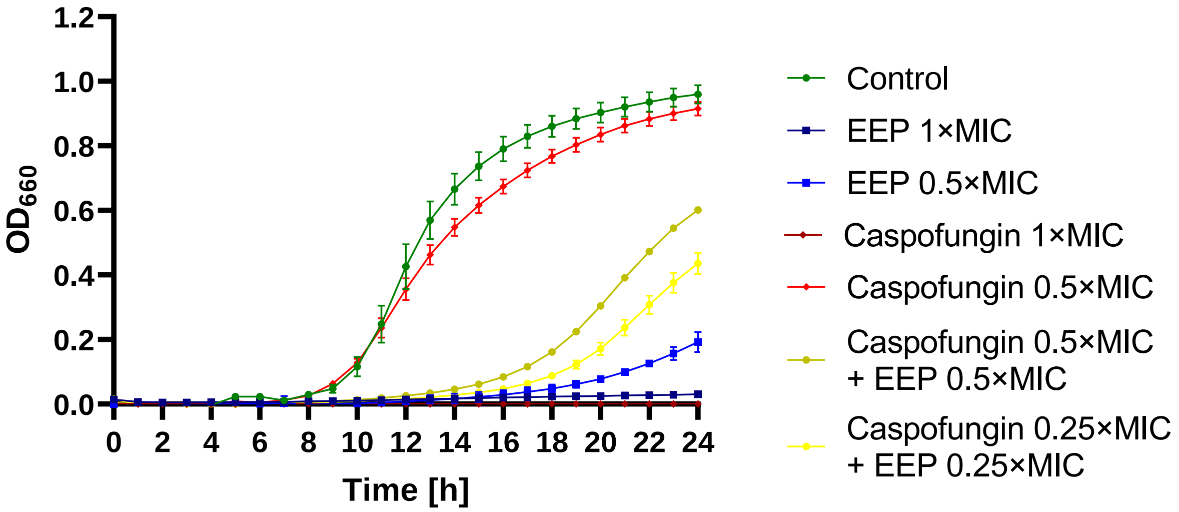

1.2
1.0
0.8
0.6
0.4
0.2
0.0
$OD_{660}$
0
2
4
6
8
10
12
14
16
18
20
22
24
Time [h]
Control
EEP 1×MIC
EEP 0.5×MIC
Caspofungin 1×MIC
Caspofungin 0.5×MIC
Caspofungin 0.5×MIC + EEP 0.5×MIC
Caspofungin 0.25×MIC + EEP 0.25×MIC

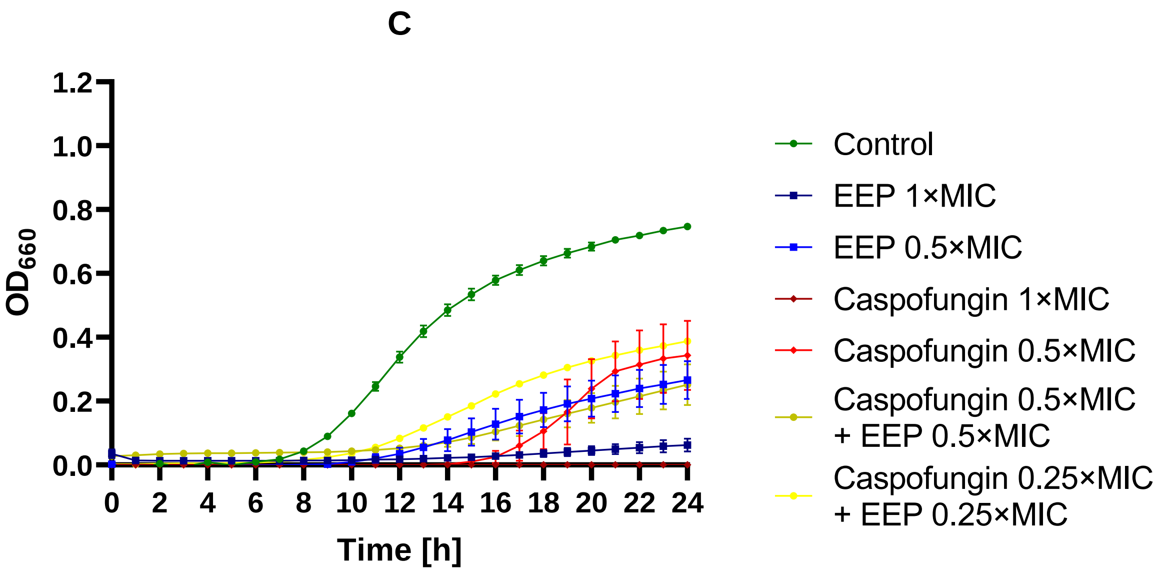


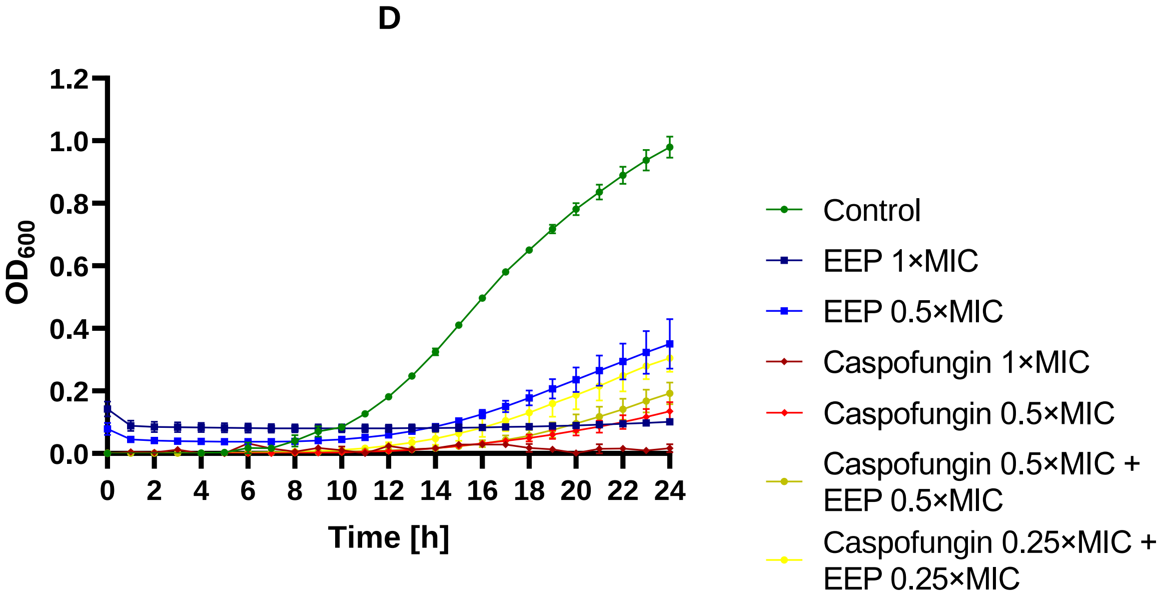


Figure S20 – Antifungal activity of EEP, caspofungin, and their mixtures – growth curves (monitoring of the growth kinetics of the yeast cells measured as turbidity of the cultures at wavelength 660 nm or 600 nm for *C. auris*) against A – *C. albicans*, B – *N. glabratus*, C – *P. kudriavzevii*, and D – *C. auris*.

A

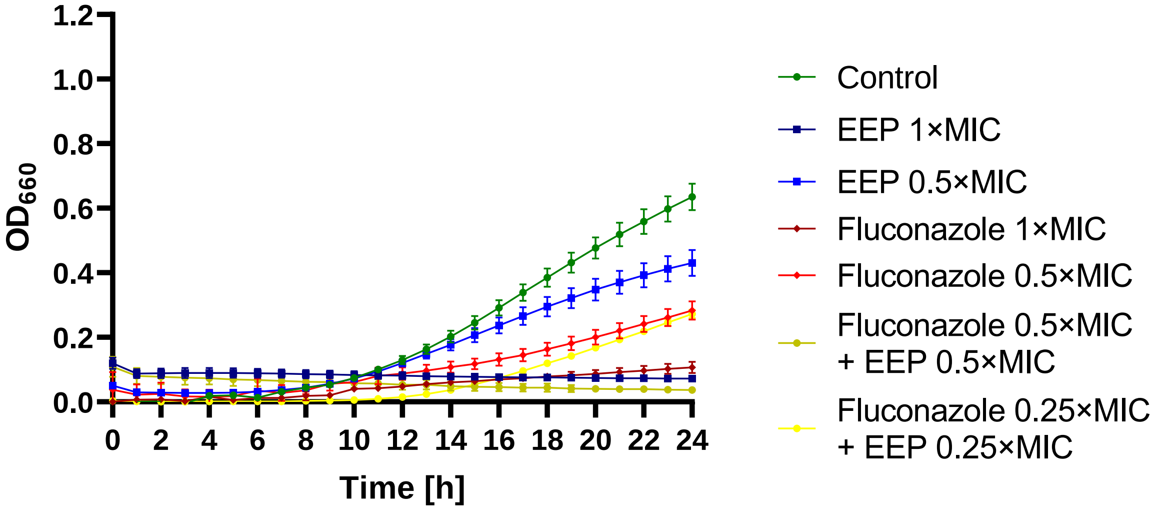
1.2
1.0
0.8
0.6
0.4
0.2
0.0
$OD_{660}$
0 2 4 6 8 10 12 14 16 18 20 22 24
Time [h]
Control
EEP 1×MIC
EEP 0.5×MIC
Fluconazole 1×MIC
Fluconazole 0.5×MIC
Fluconazole 0.5×MIC + EEP 0.5×MIC
Fluconazole 0.25×MIC + EEP 0.25×MIC

B

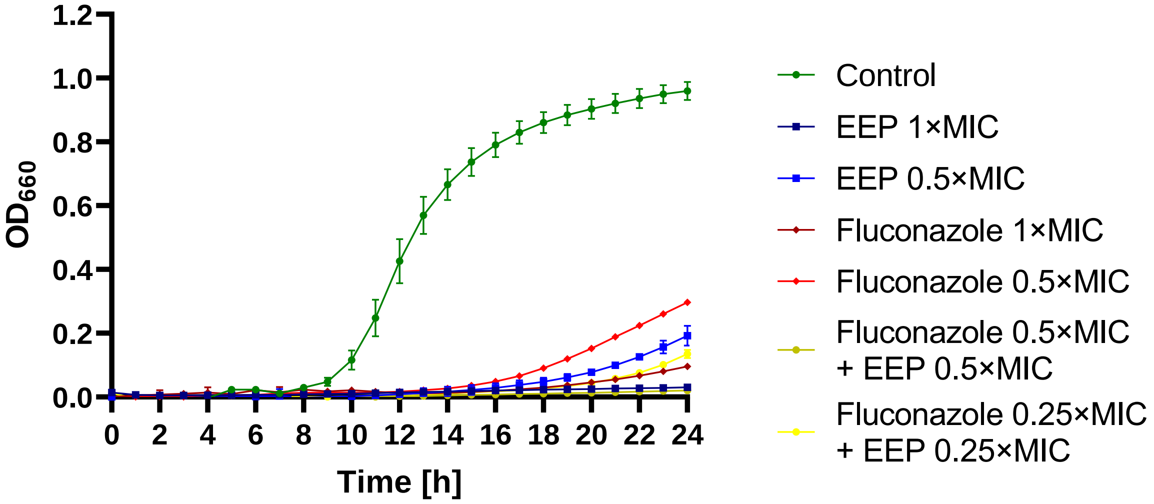
1.2
1.0
0.8
0.6
0.4
0.2
0.0
$OD_{660}$
0 2 4 6 8 10 12 14 16 18 20 22 24
Time [h]
Control
EEP 1×MIC
EEP 0.5×MIC
Fluconazole 1×MIC
Fluconazole 0.5×MIC
Fluconazole 0.5×MIC + EEP 0.5×MIC
Fluconazole 0.25×MIC + EEP 0.25×MIC

C

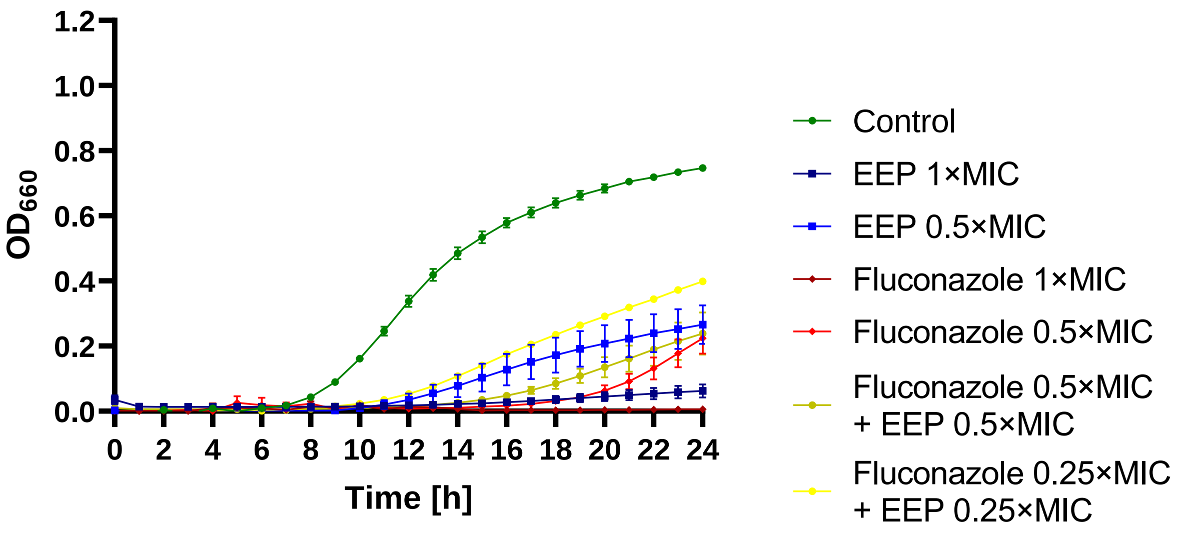


D

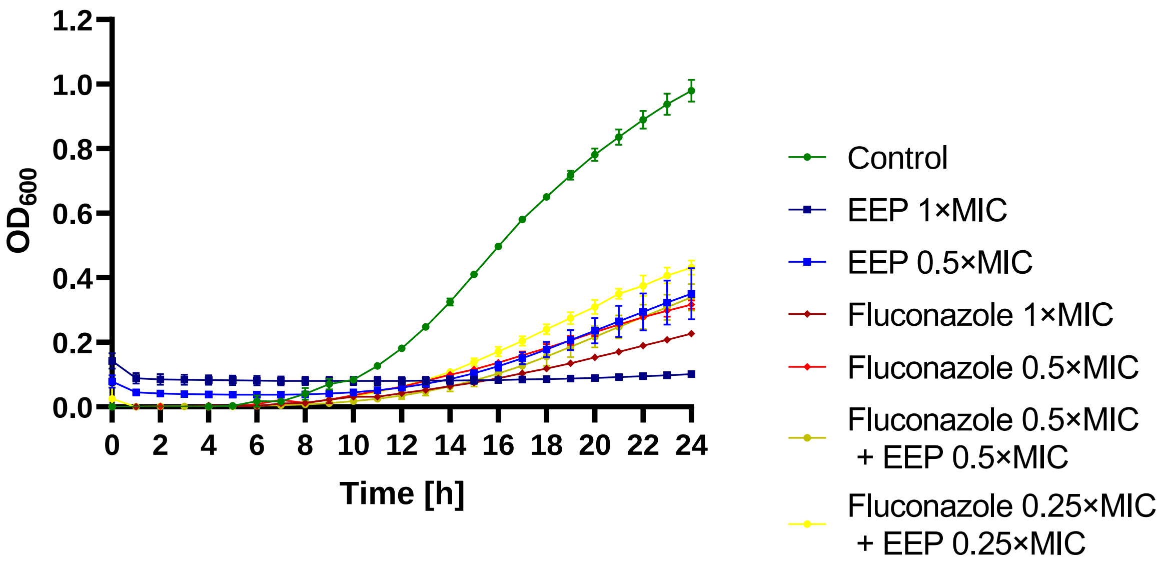


Figure S21 – Antifungal activity of EEP, fluconazole, and their mixtures – growth curves (monitoring of the growth kinetics of the yeast cells measured as turbidity of the cultures at wavelength 660 nm or 600 nm for *C. auris*) against A – *C. albicans*, B – *N. glabratus*, C – *P. kudriavzevii*, and D – *C. auris*.

A

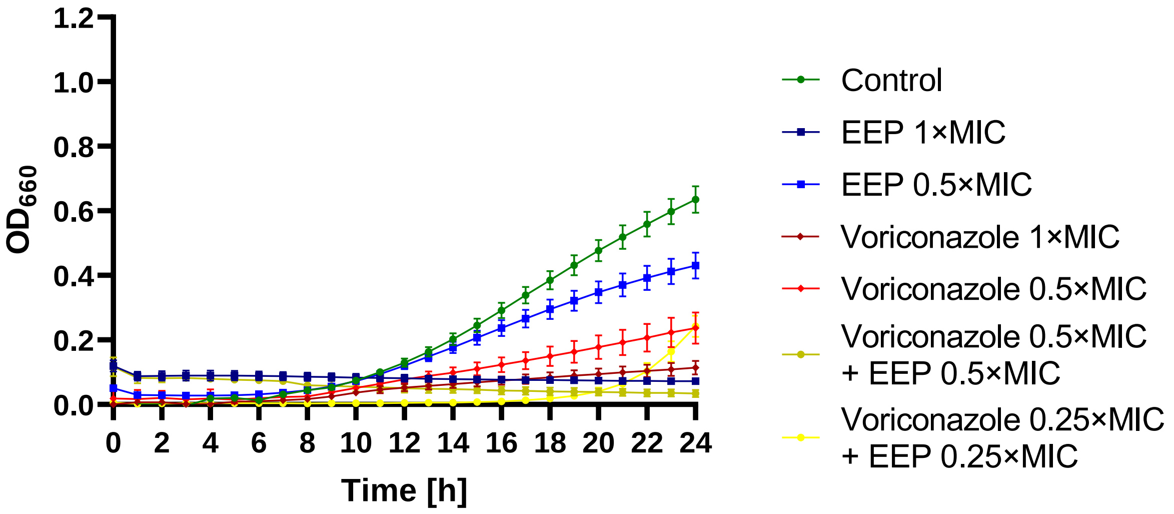
1.2
1.0
0.8
0.6
0.4
0.2
0.0
$OD_{660}$
0 2 4 6 8 10 12 14 16 18 20 22 24
Time [h]
Control
EEP 1×MIC
EEP 0.5×MIC
Voriconazole 1×MIC
Voriconazole 0.5×MIC
Voriconazole 0.5×MIC + EEP 0.5×MIC
Voriconazole 0.25×MIC + EEP 0.25×MIC

B

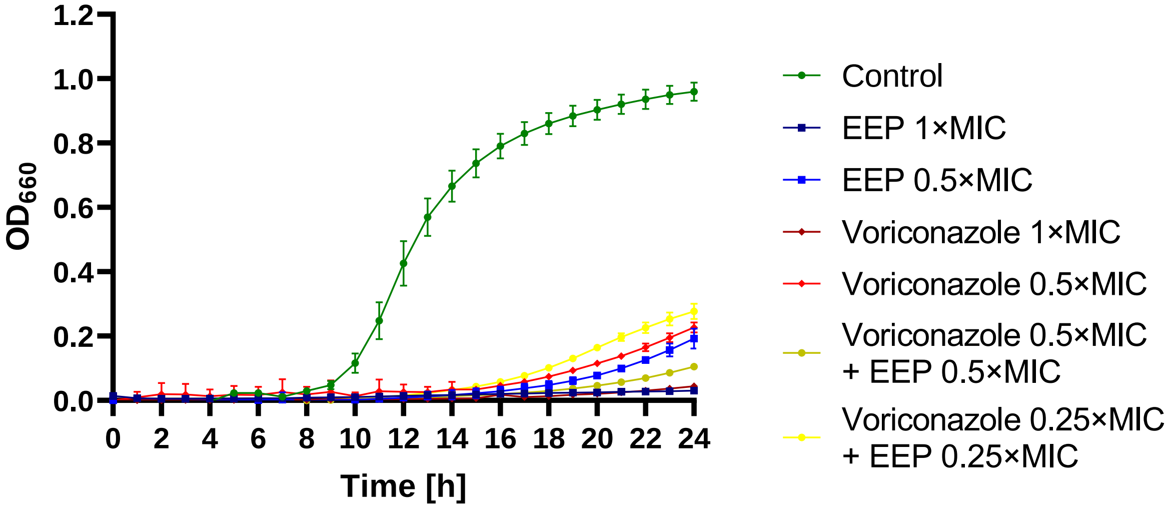
1.2
1.0
0.8
0.6
0.4
0.2
0.0
$OD_{660}$
0 2 4 6 8 10 12 14 16 18 20 22 24
Time [h]
Control
EEP 1×MIC
EEP 0.5×MIC
Voriconazole 1×MIC
Voriconazole 0.5×MIC
Voriconazole 0.5×MIC + EEP 0.5×MIC
Voriconazole 0.25×MIC + EEP 0.25×MIC

C

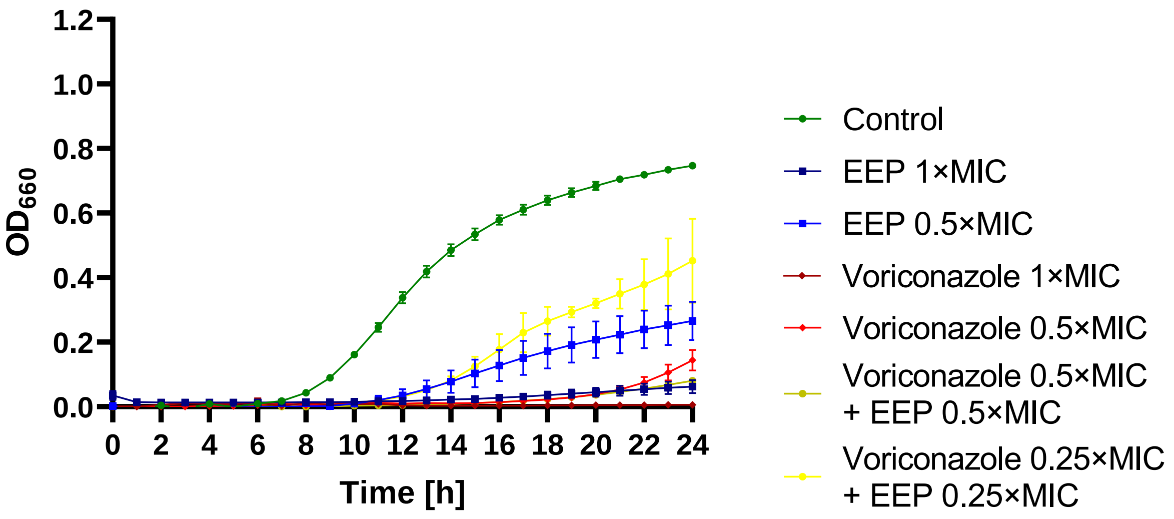


D

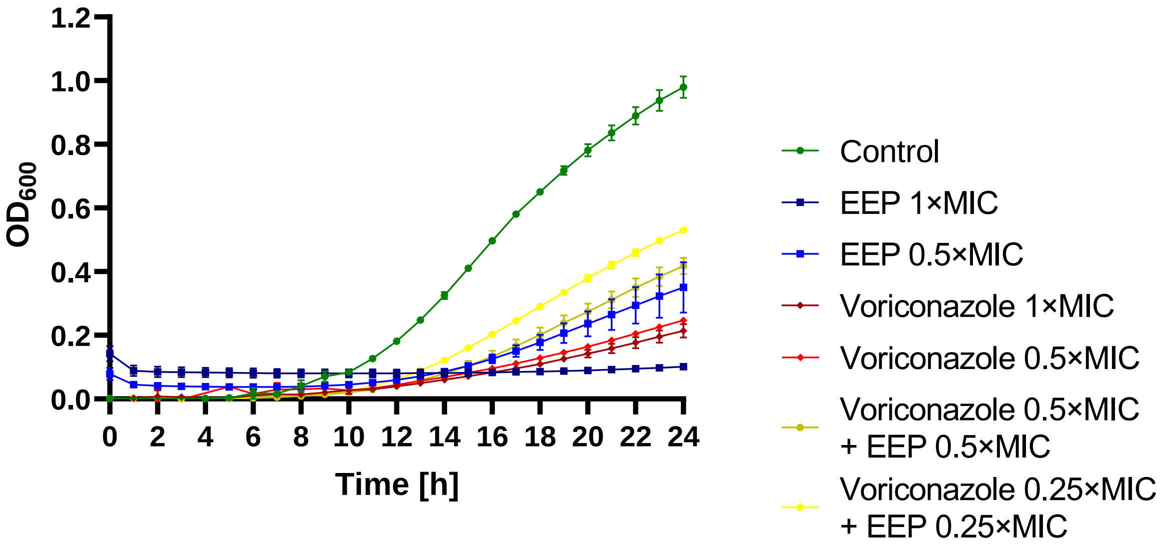


Figure S22 – Antifungal activity of EEP, voriconazole, and their mixtures – growth curves (monitoring of the growth kinetics of the yeast cells measured as turbidity of the cultures at wavelength 660 nm or 600 nm for *C. auris*) against A – *C. albicans*, B – *N. glabratus*, C – *P. kudriavzevii* and D – *C. auris*.

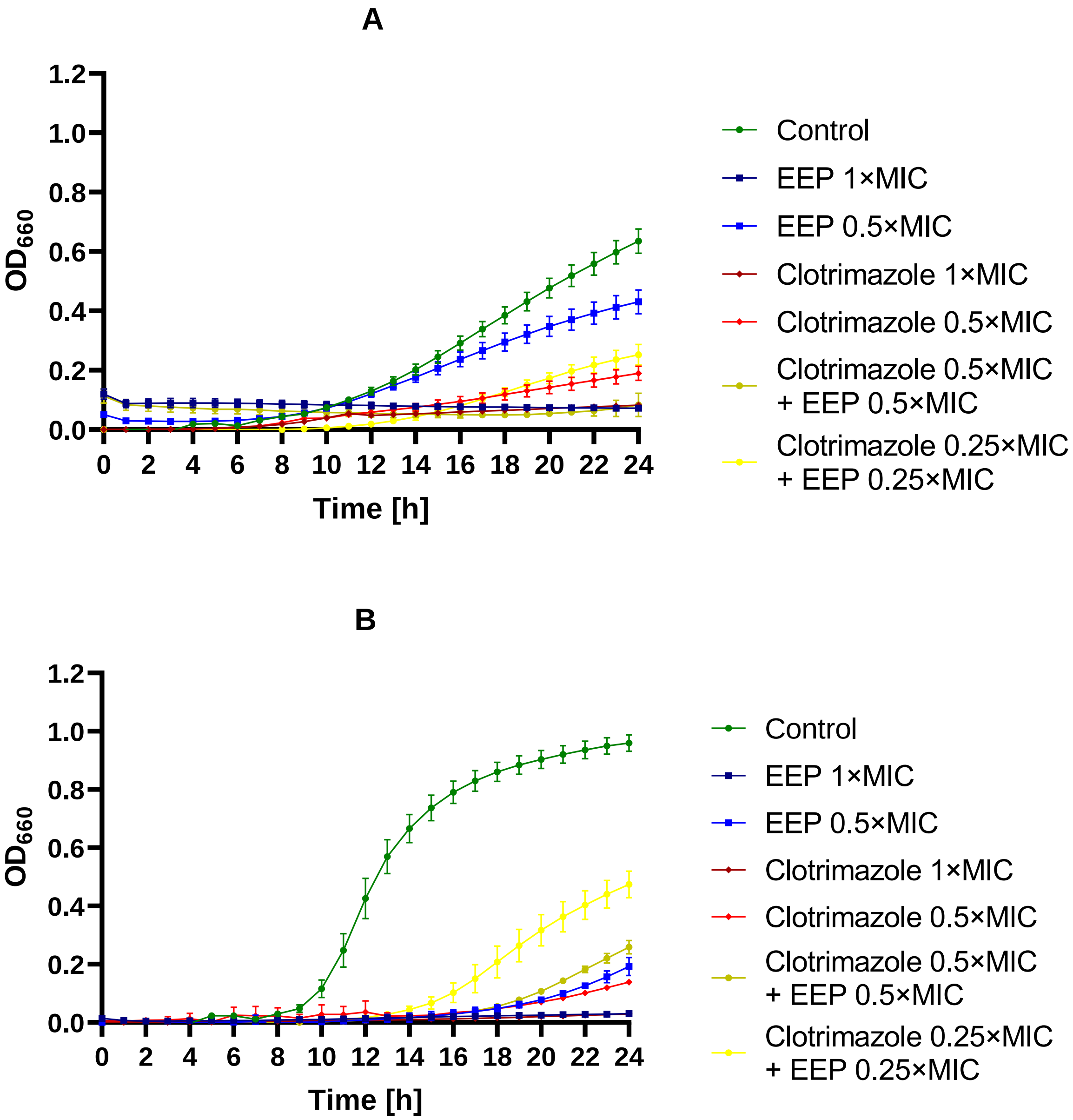
A
1.2
1.0
0.8
0.6
0.4
0.2
0.0
OD660
0 2 4 6 8 10 12 14 16 18 20 22 24
Time [h]
Control
EEP 1×MIC
EEP 0.5×MIC
Clotrimazole 1×MIC
Clotrimazole 0.5×MIC
Clotrimazole 0.5×MIC + EEP 0.5×MIC
Clotrimazole 0.25×MIC + EEP 0.25×MIC
B
1.2
1.0
0.8
0.6
0.4
0.2
0.0
OD660
0 2 4 6 8 10 12 14 16 18 20 22 24
Time [h]
Control
EEP 1×MIC
EEP 0.5×MIC
Clotrimazole 1×MIC
Clotrimazole 0.5×MIC
Clotrimazole 0.5×MIC + EEP 0.5×MIC
Clotrimazole 0.25×MIC + EEP 0.25×MIC

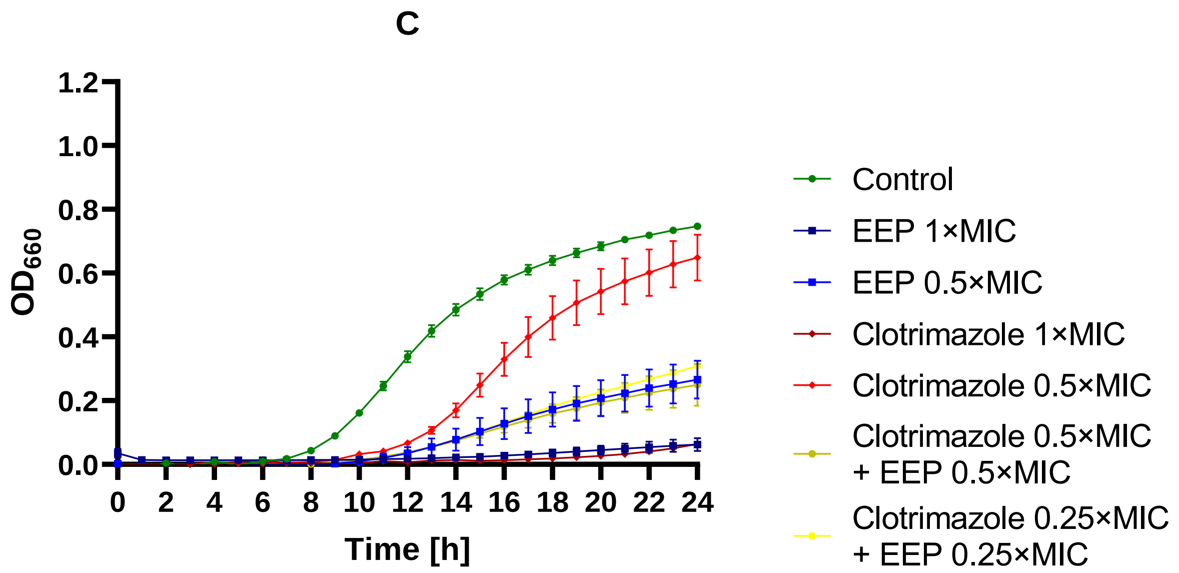


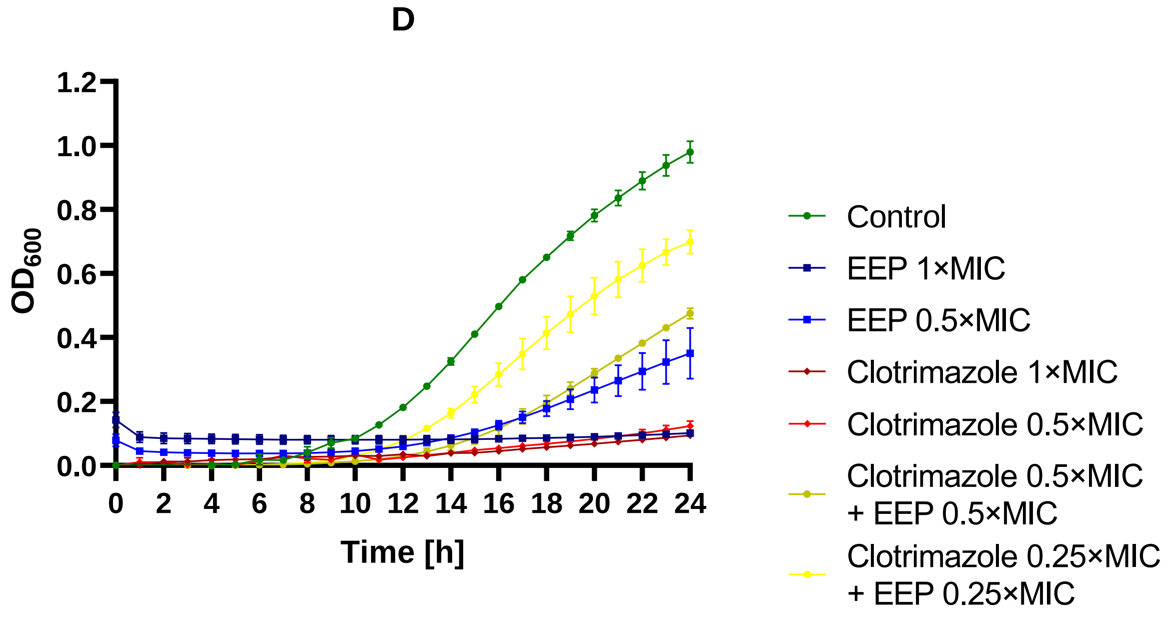


Figure S23 – Antifungal activity of EEP, clotrimazole, and their mixtures – growth curves (monitoring of the growth kinetics of the yeast cells measured as turbidity of the cultures at wavelength 660 nm or 600 nm for *C. auris*) against A – *C. albicans*, B – *N. glabratus*, C – *P. kudriavzevii*, and D – *C. auris*.

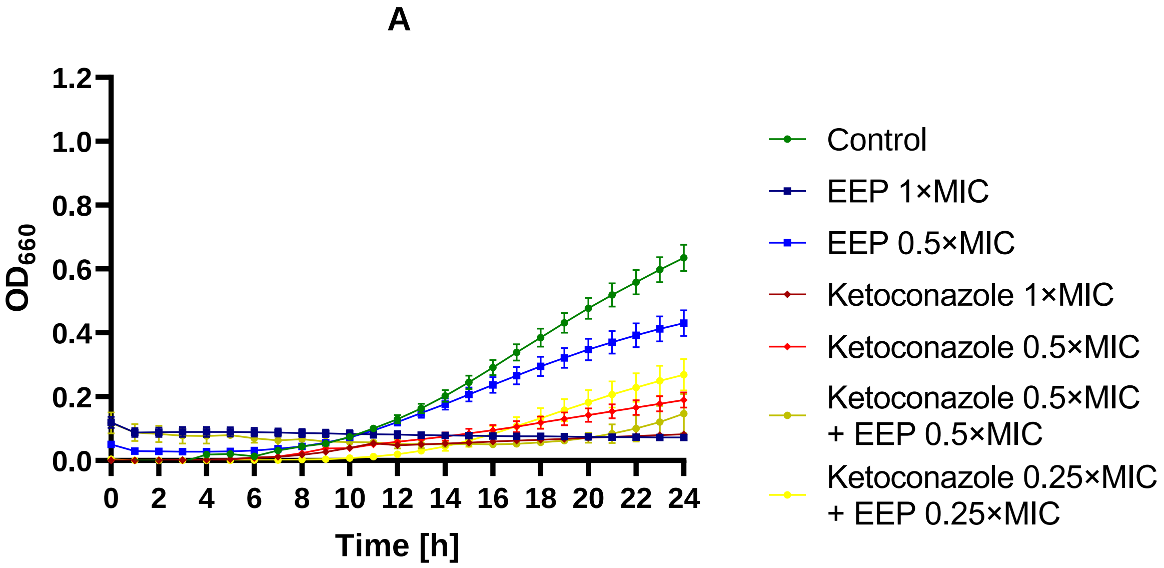
A
1.2
1.0
0.8
0.6
0.4
0.2
0.0
OD660
0 2 4 6 8 10 12 14 16 18 20 22 24
Time [h]
Control
EEP 1×MIC
EEP 0.5×MIC
Ketoconazole 1×MIC
Ketoconazole 0.5×MIC
Ketoconazole 0.5×MIC + EEP 0.5×MIC
Ketoconazole 0.25×MIC + EEP 0.25×MIC

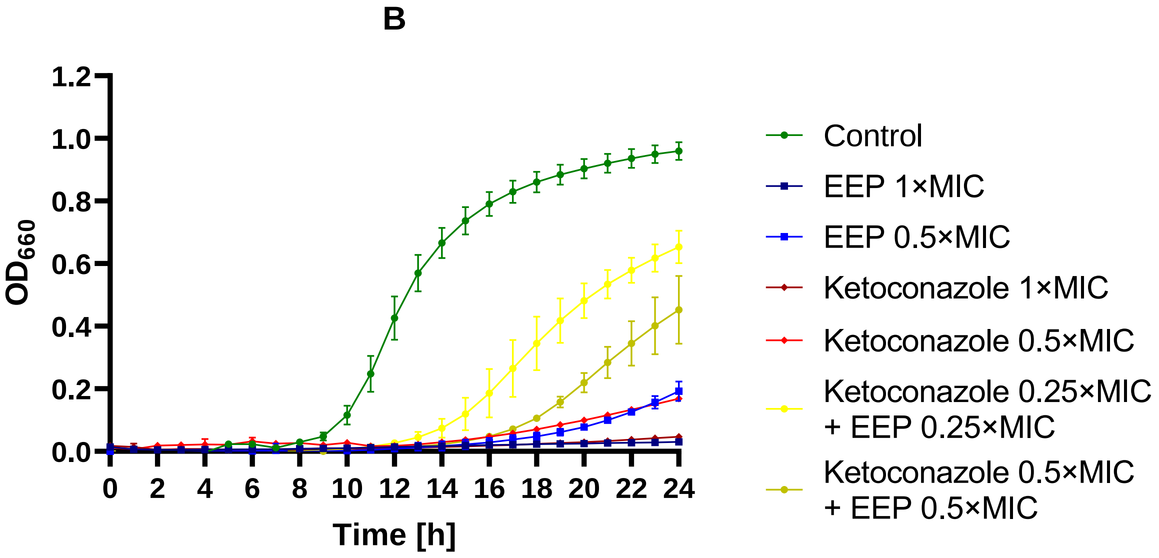
B
1.2
1.0
0.8
0.6
0.4
0.2
0.0
OD660
0 2 4 6 8 10 12 14 16 18 20 22 24
Time [h]
Control
EEP 1×MIC
EEP 0.5×MIC
Ketoconazole 1×MIC
Ketoconazole 0.5×MIC
Ketoconazole 0.25×MIC + EEP 0.25×MIC
Ketoconazole 0.5×MIC + EEP 0.5×MIC

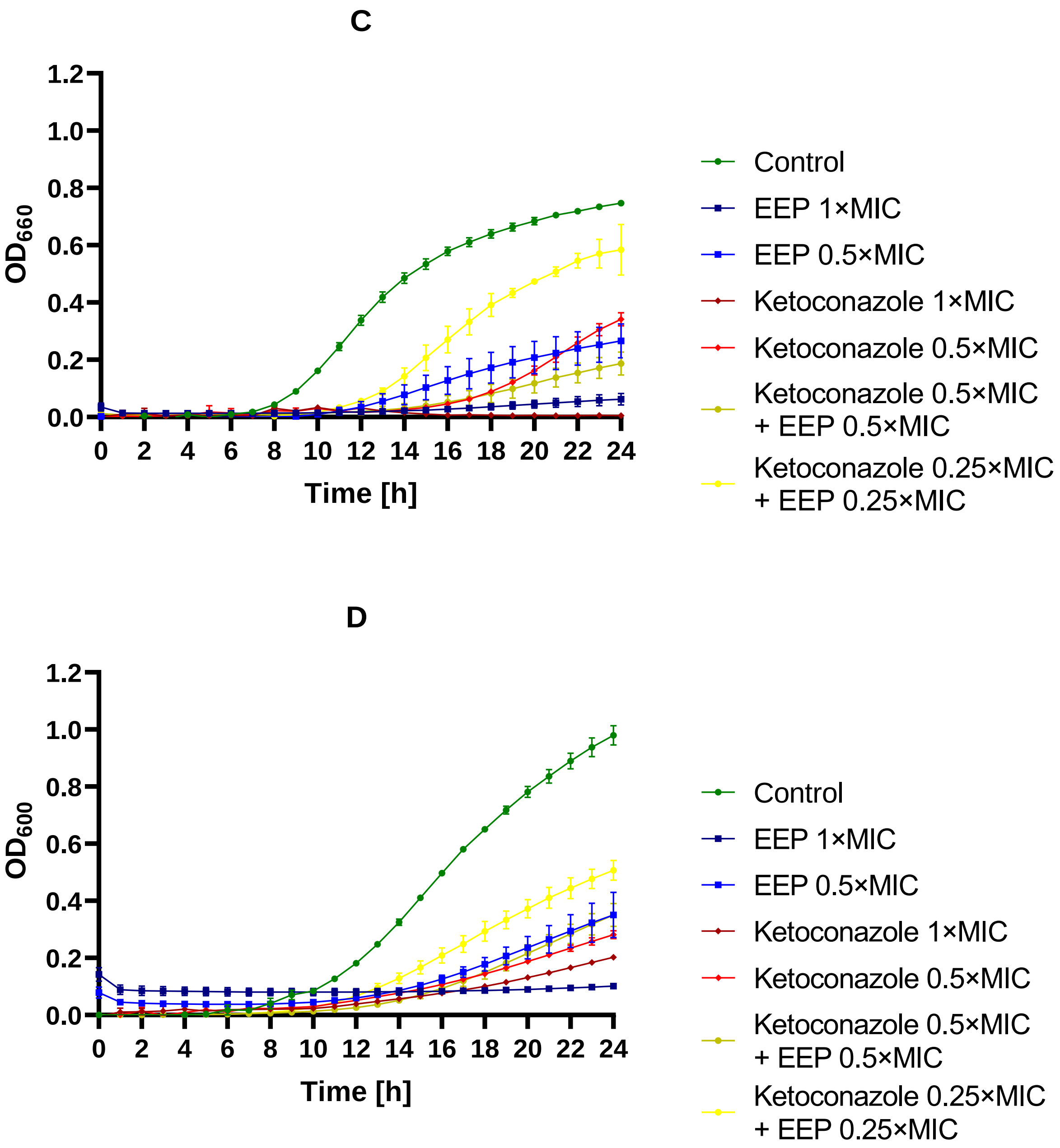


Figure S24 – Antifungal activity of EEP, ketoconazole, and their mixtures – growth curves (monitoring of the growth kinetics of the yeast cells measured as turbidity of the cultures at wavelength 660 nm or 600 nm for *C. auris*) against A – *C. albicans*, B – *N. glabratus*, C – *P. kudriavzevii*, and D – *C. auris*.

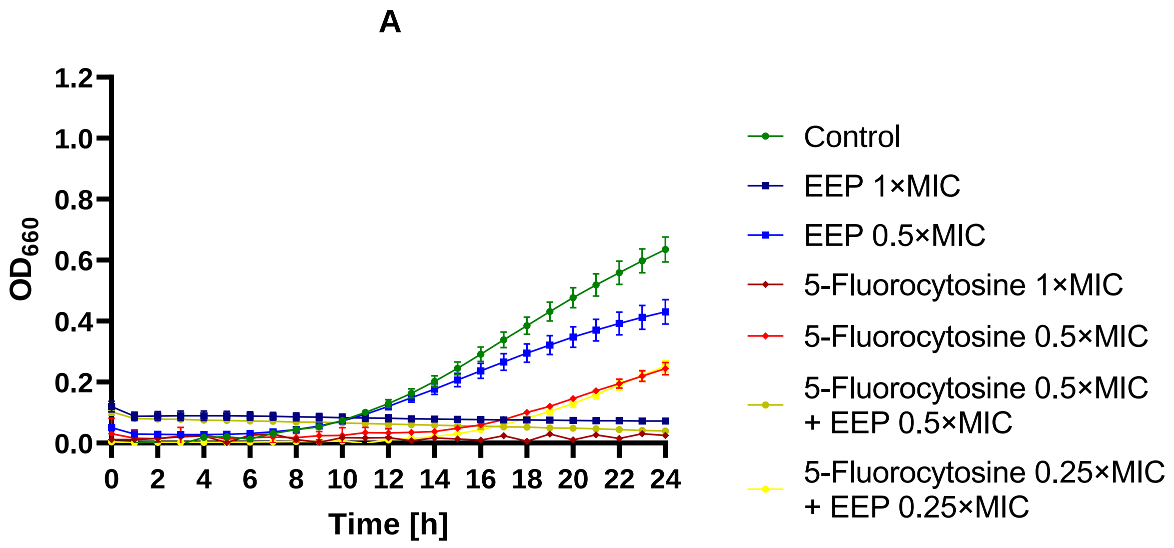
A
OD660
1.2
1.0
0.8
0.6
0.4
0.2
0.0
0 2 4 6 8 10 12 14 16 18 20 22 24
Time [h]
Control
EEP 1×MIC
EEP 0.5×MIC
5-Fluorocytosine 1×MIC
5-Fluorocytosine 0.5×MIC
5-Fluorocytosine 0.5×MIC + EEP 0.5×MIC
5-Fluorocytosine 0.25×MIC + EEP 0.25×MIC

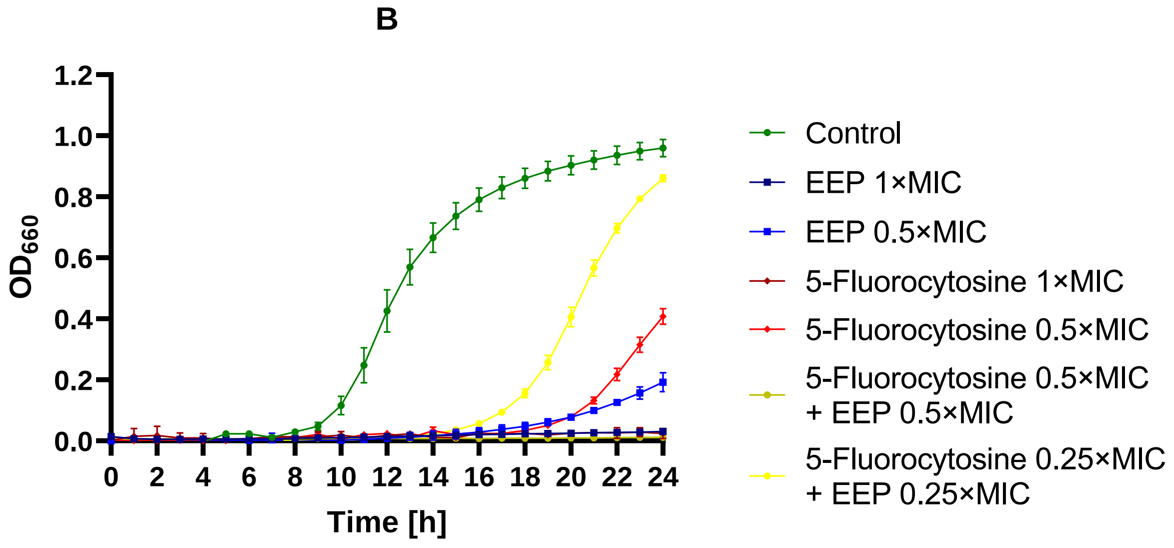
B
OD660
1.2
1.0
0.8
0.6
0.4
0.2
0.0
0 2 4 6 8 10 12 14 16 18 20 22 24
Time [h]
Control
EEP 1×MIC
EEP 0.5×MIC
5-Fluorocytosine 1×MIC
5-Fluorocytosine 0.5×MIC
5-Fluorocytosine 0.5×MIC + EEP 0.5×MIC
5-Fluorocytosine 0.25×MIC + EEP 0.25×MIC

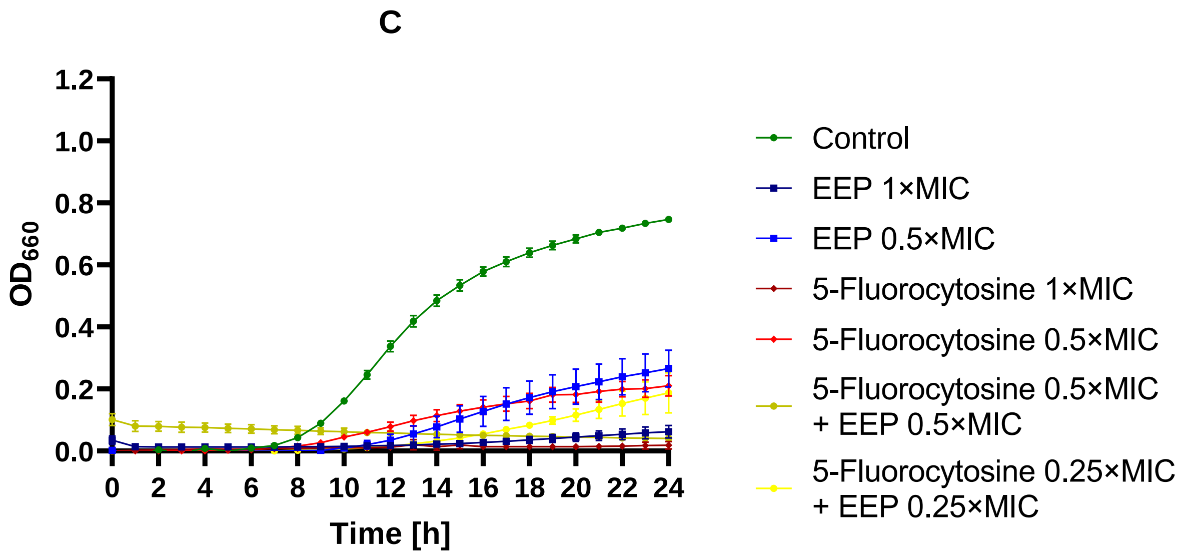


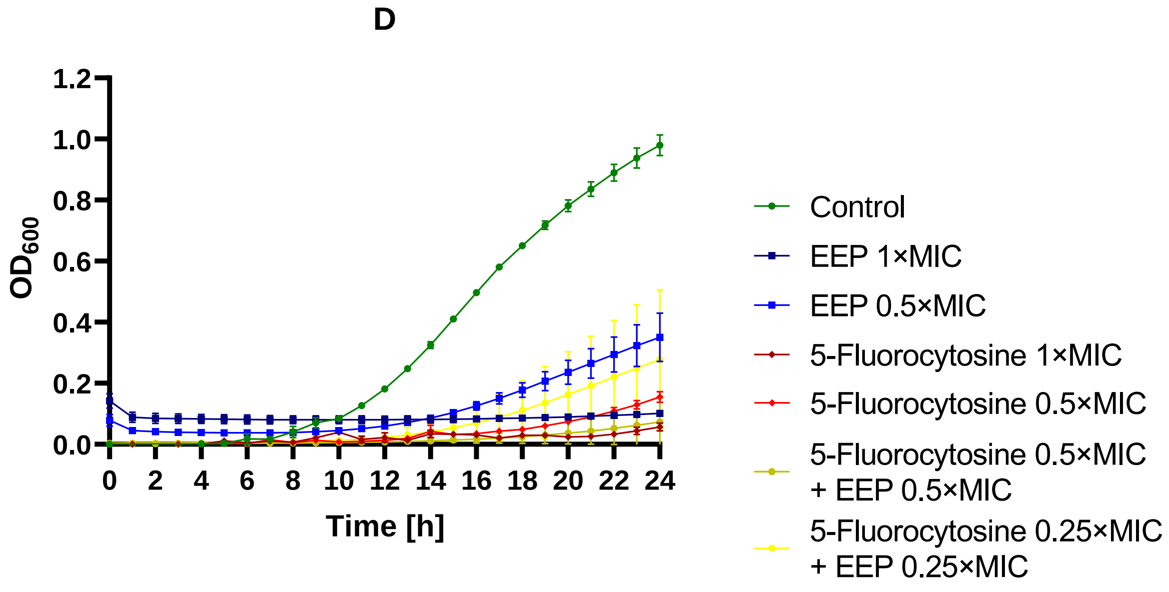


Figure S25 – Antifungal activity of EEP, 5-fluorocytosine, and their mixtures – growth curves (monitoring of the growth kinetics of the yeast cells measured as turbidity of the cultures at wavelength 660 nm or 600 nm for *C. auris*) against A – *C. albicans*, B – *N. glabratus*, C – *P. kudriavzevii*, and D – *C. auris*.

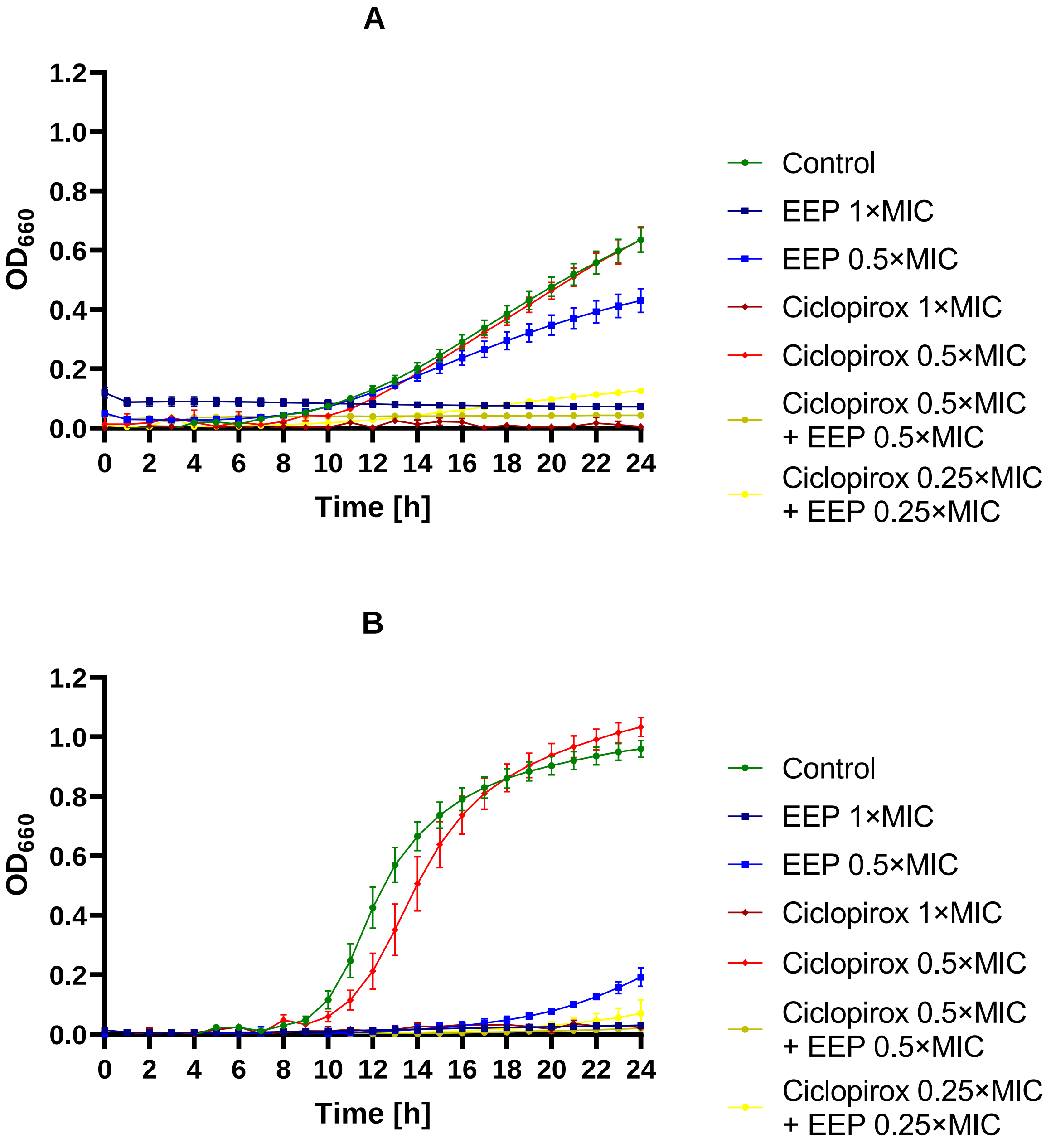
A
B
OD660
Time [h]
Control
EEP 1×MIC
EEP 0.5×MIC
Ciclopirox 1×MIC
Ciclopirox 0.5×MIC
Ciclopirox 0.5×MIC + EEP 0.5×MIC
Ciclopirox 0.25×MIC + EEP 0.25×MIC

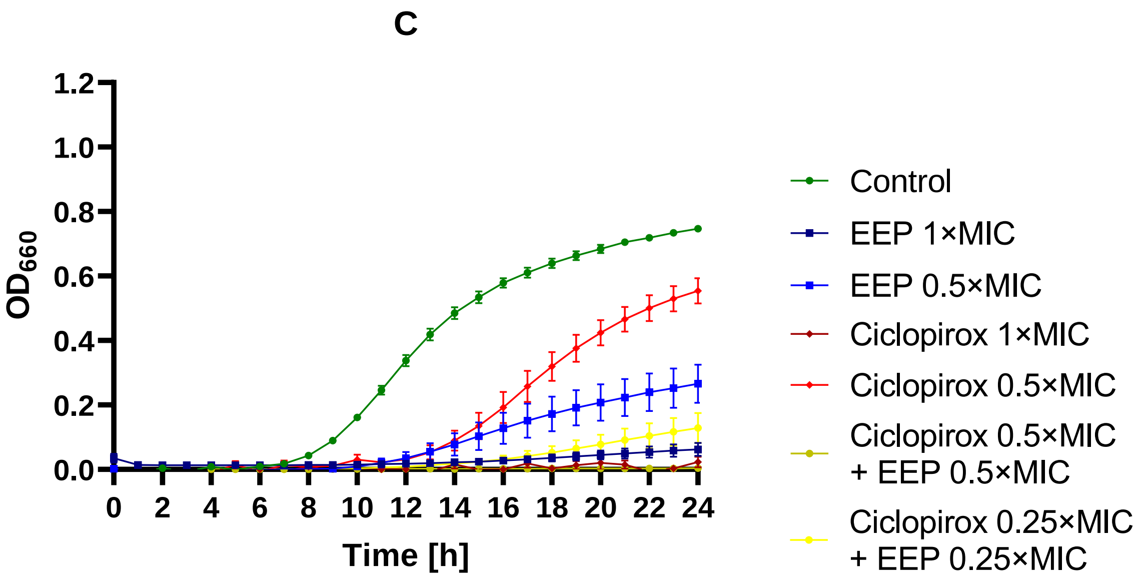


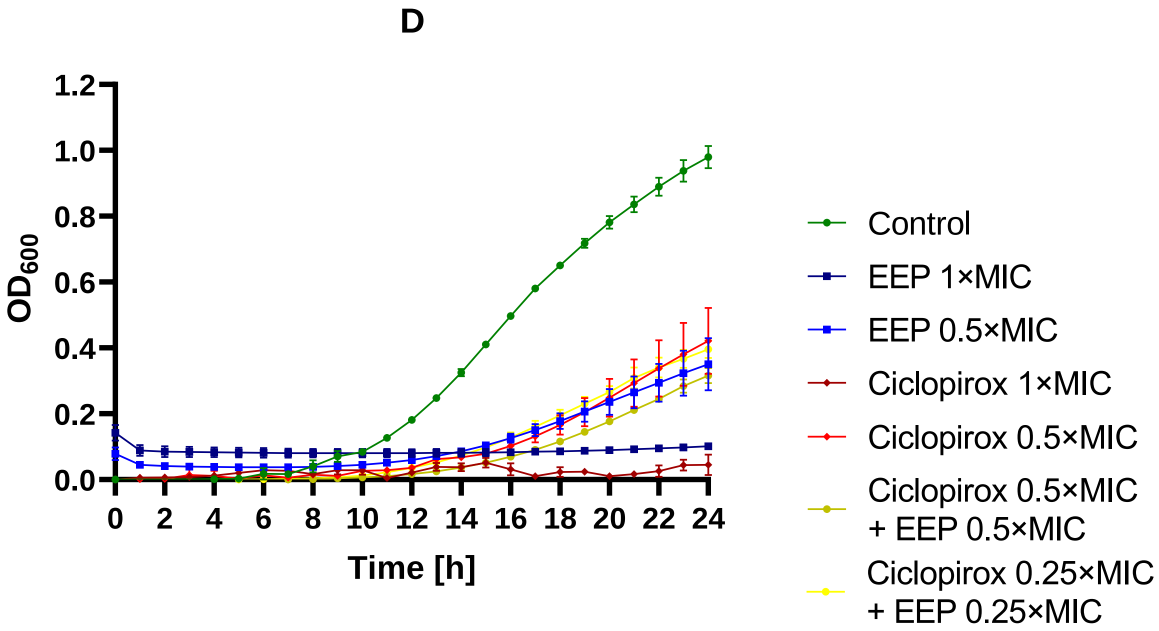


Figure S26 – Antifungal activity of EEP, ciclopirox, and their mixtures – growth curves (monitoring of the growth kinetics of the yeast cells measured as turbidity of the cultures at wavelength 660 nm or 600 nm for *C. auris*) against A – *C. albicans*, B – *N. glabratus*, C – *P. kudriavzevii*, and D – *C. auris*.

A

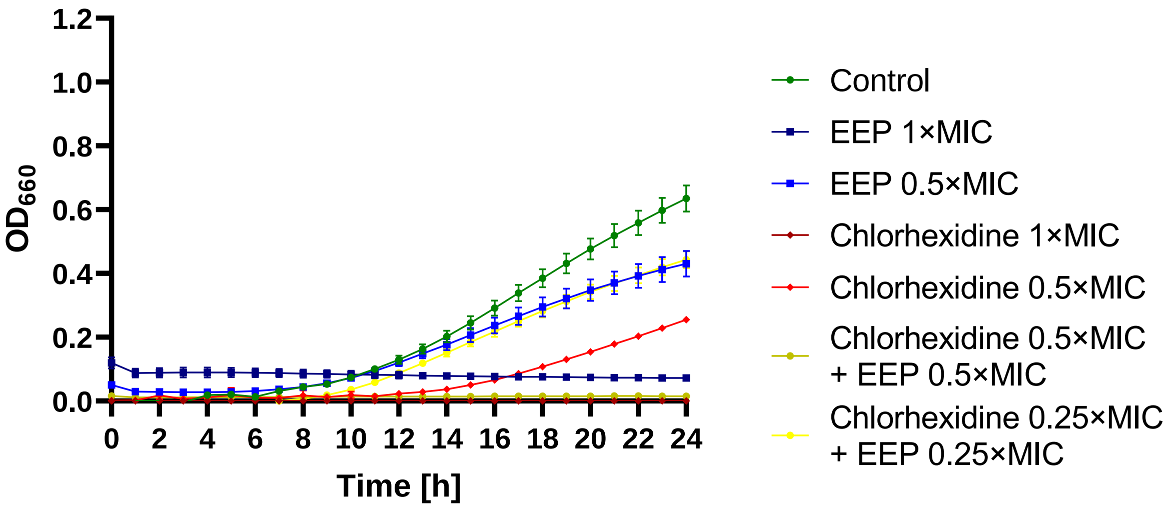
1.2
1.0
0.8
0.6
0.4
0.2
0.0
OD660
0 2 4 6 8 10 12 14 16 18 20 22 24
Time [h]
Control
EEP 1×MIC
EEP 0.5×MIC
Chlorhexidine 1×MIC
Chlorhexidine 0.5×MIC
Chlorhexidine 0.5×MIC + EEP 0.5×MIC
Chlorhexidine 0.25×MIC + EEP 0.25×MIC

B

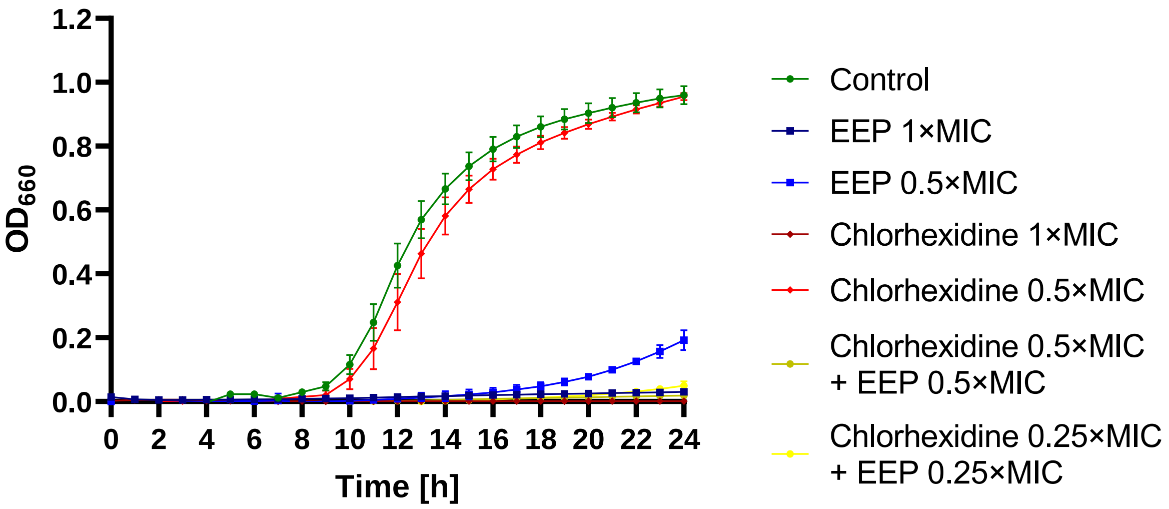
1.2
1.0
0.8
0.6
0.4
0.2
0.0
OD660
0 2 4 6 8 10 12 14 16 18 20 22 24
Time [h]
Control
EEP 1×MIC
EEP 0.5×MIC
Chlorhexidine 1×MIC
Chlorhexidine 0.5×MIC
Chlorhexidine 0.5×MIC + EEP 0.5×MIC
Chlorhexidine 0.25×MIC + EEP 0.25×MIC

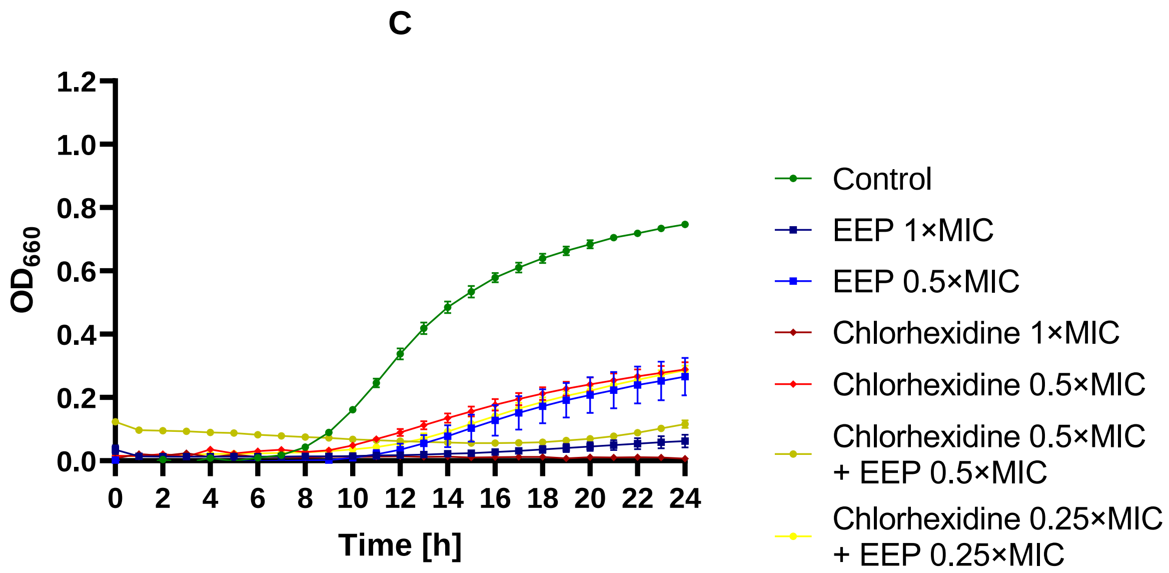


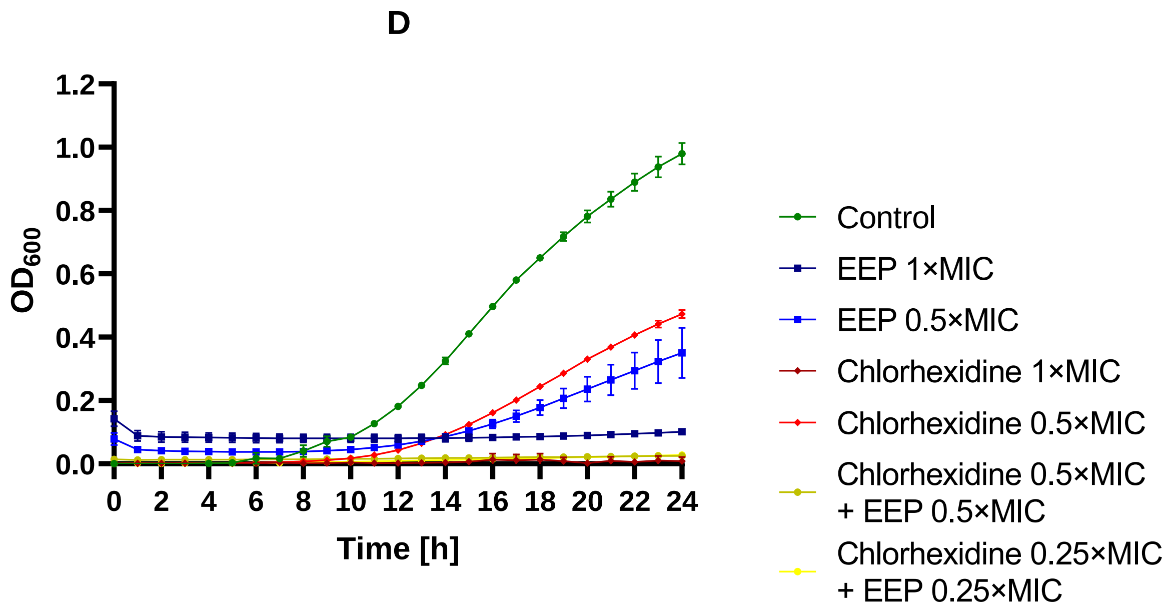


Figure S27 – Antifungal activity of EEP, chlorhexidine, and their mixtures – growth curves (monitoring of the growth kinetics of the yeast cells measured as turbidity of the cultures at wavelength 660 nm or 600 nm for *C. auris*) against A – *C. albicans*, B – *N. glabratus*, C – *P. kudriavzevii*, and D – *C. auris*.

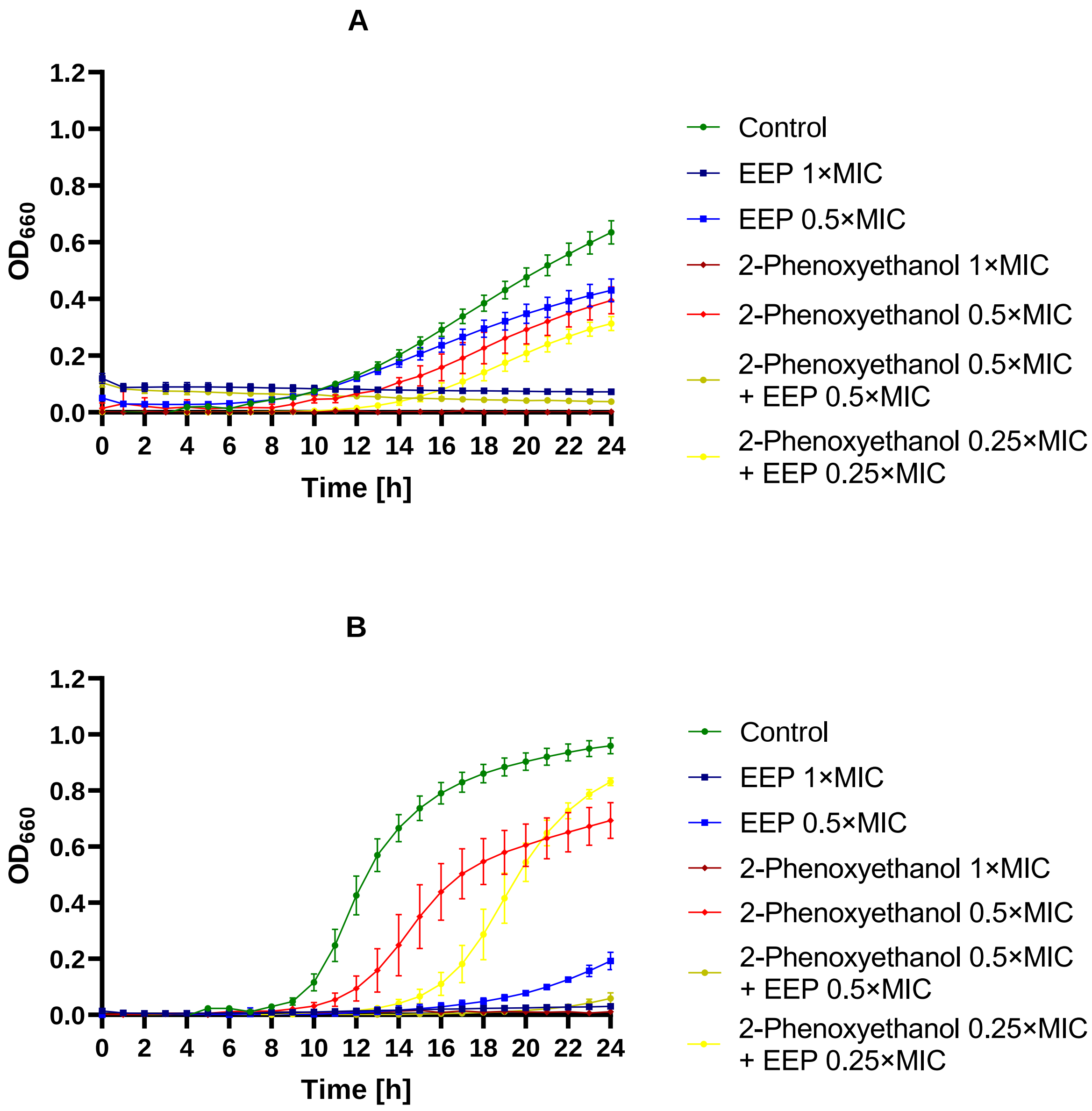
A
B
OD660
Time [h]
Control
EEP 1×MIC
EEP 0.5×MIC
2-Phenoxyethanol 1×MIC
2-Phenoxyethanol 0.5×MIC
2-Phenoxyethanol 0.5×MIC + EEP 0.5×MIC
2-Phenoxyethanol 0.25×MIC + EEP 0.25×MIC

C

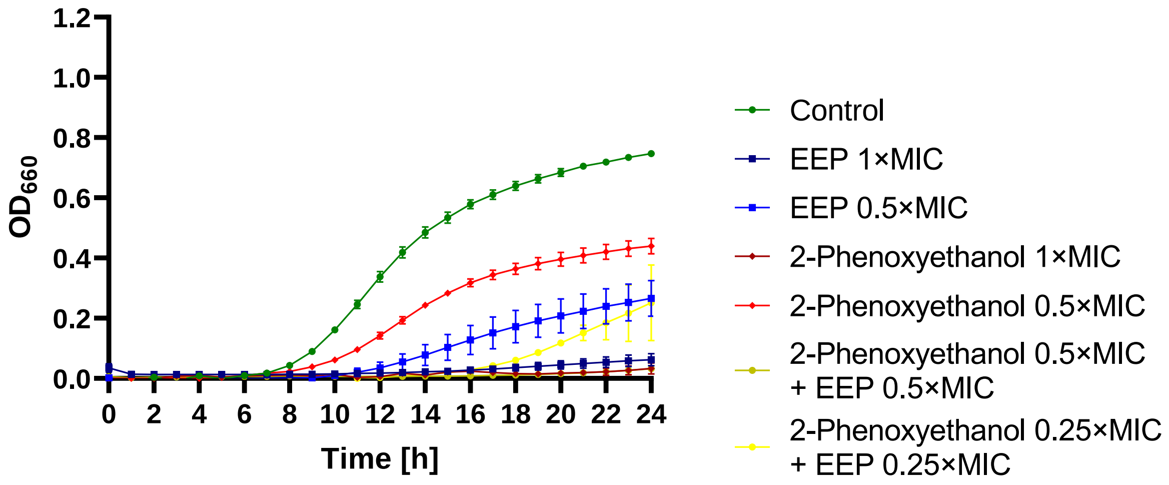


D

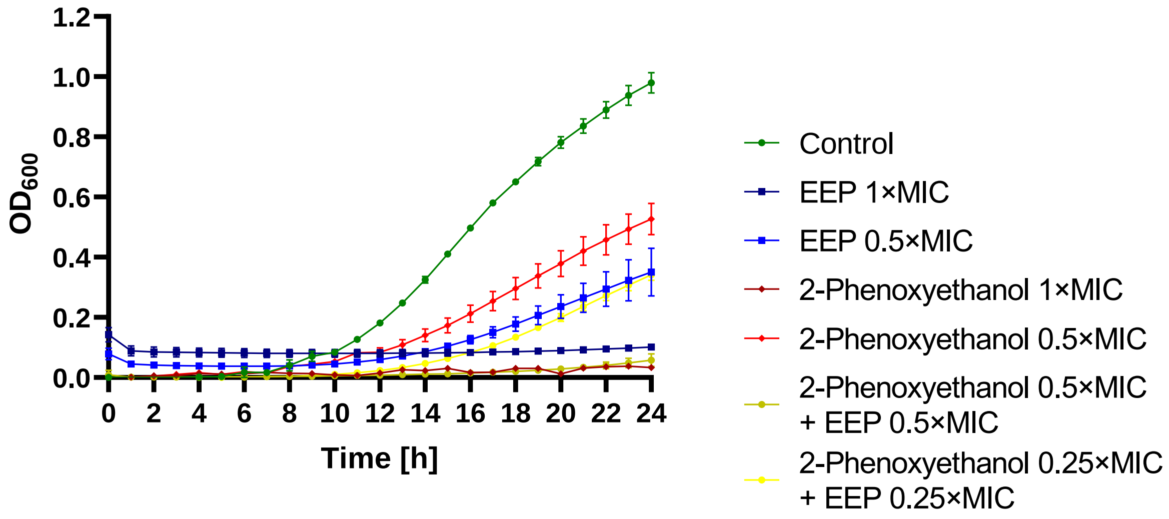


Figure S28 – Antifungal activity of EEP, 2-phenoxyethanol, and their mixtures – growth curves (monitoring of the growth kinetics of the yeast cells measured as turbidity of the cultures at wavelength 660 nm or 600 nm for *C. auris*) against A – *C. albicans*, B – *N. glabratus*, C – *P. kudriavzevii*, and D – *C. auris*.

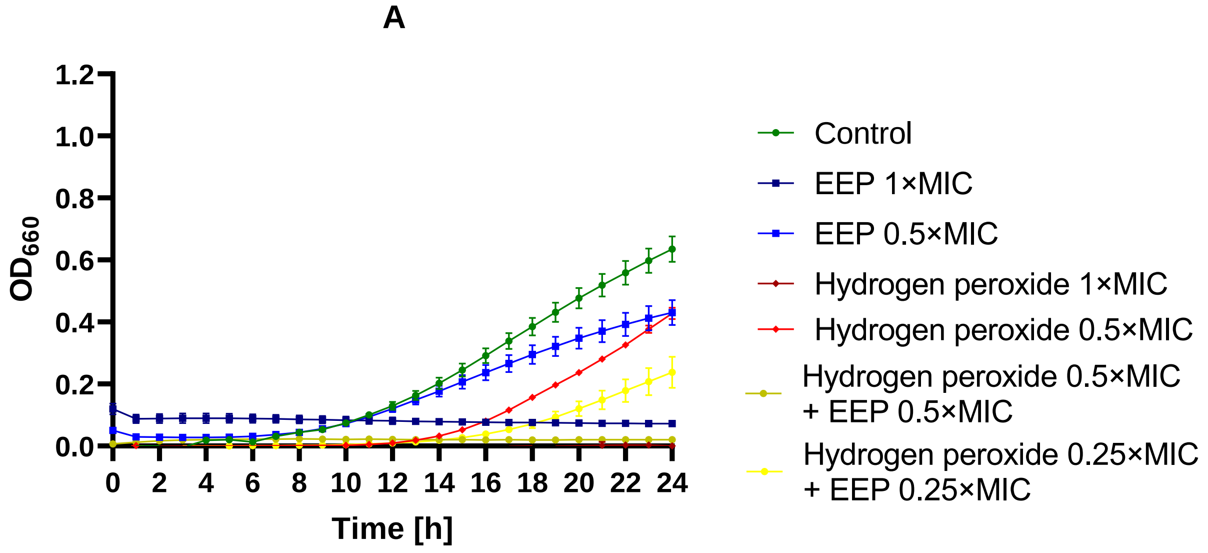
A
1.2
1.0
0.8
0.6
0.4
0.2
0.0
OD660
0
2
4
6
8
10
12
14
16
18
20
22
24
Time [h]
Control
EEP 1×MIC
EEP 0.5×MIC
Hydrogen peroxide 1×MIC
Hydrogen peroxide 0.5×MIC
Hydrogen peroxide 0.5×MIC + EEP 0.5×MIC
Hydrogen peroxide 0.25×MIC + EEP 0.25×MIC

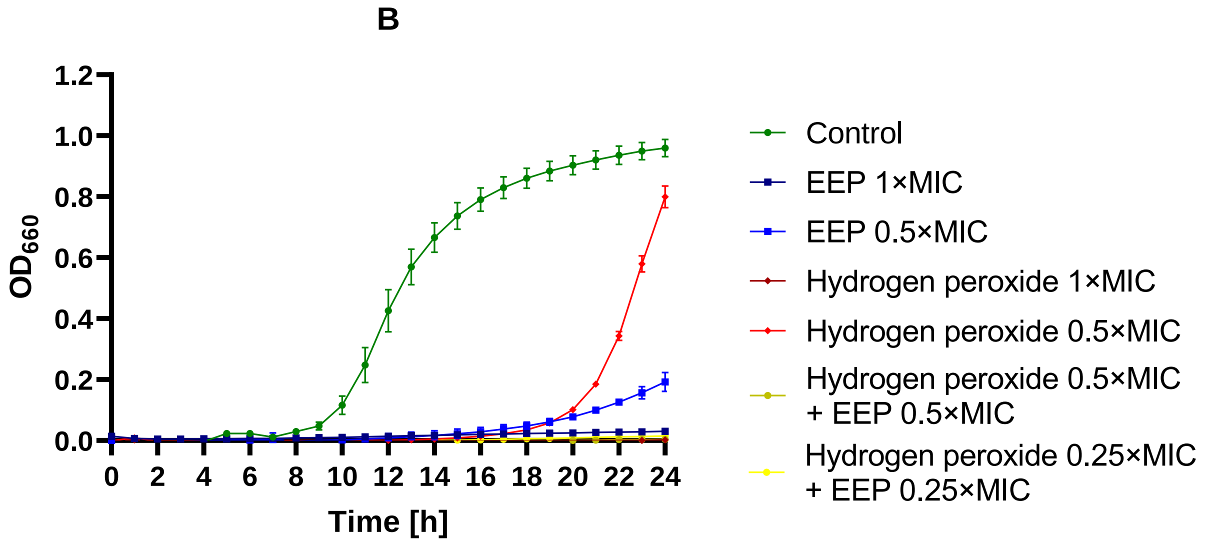
B
1.2
1.0
0.8
0.6
0.4
0.2
0.0
OD660
0
2
4
6
8
10
12
14
16
18
20
22
24
Time [h]
Control
EEP 1×MIC
EEP 0.5×MIC
Hydrogen peroxide 1×MIC
Hydrogen peroxide 0.5×MIC
Hydrogen peroxide 0.5×MIC + EEP 0.5×MIC
Hydrogen peroxide 0.25×MIC + EEP 0.25×MIC

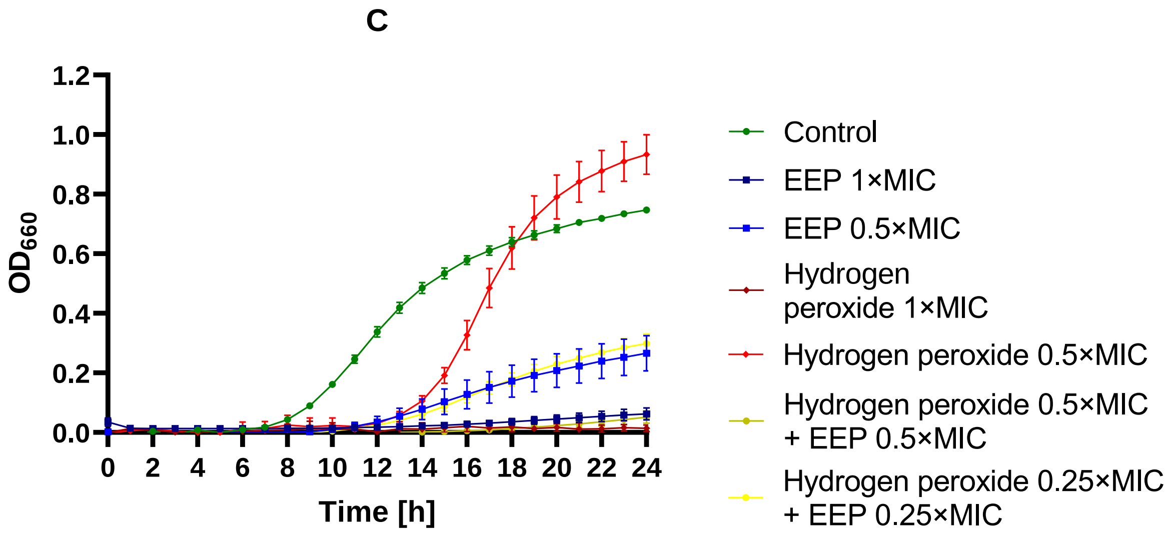


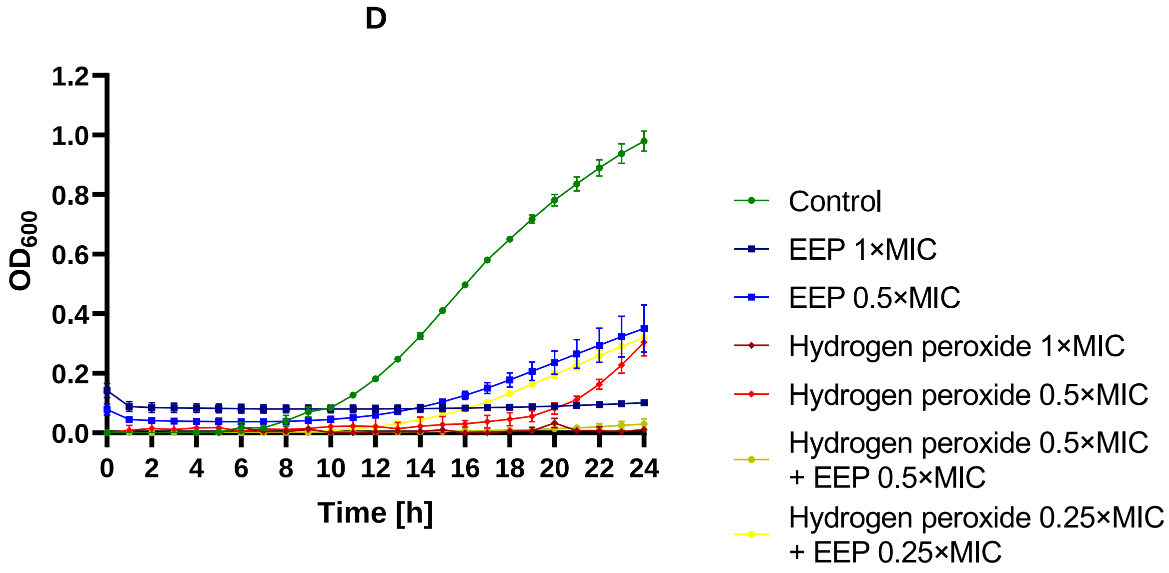


Figure S29 – Antifungal activity of EEP, hydrogen peroxide, and their mixtures – growth curves (monitoring of the growth kinetics of the yeast cells measured as turbidity of the cultures at wavelength 660 nm or 600 nm for *C. auris*) against A – *C. albicans*, B – *N. glabratus*, C – *P. kudriavzevii*, and D – *C. auris*.

A

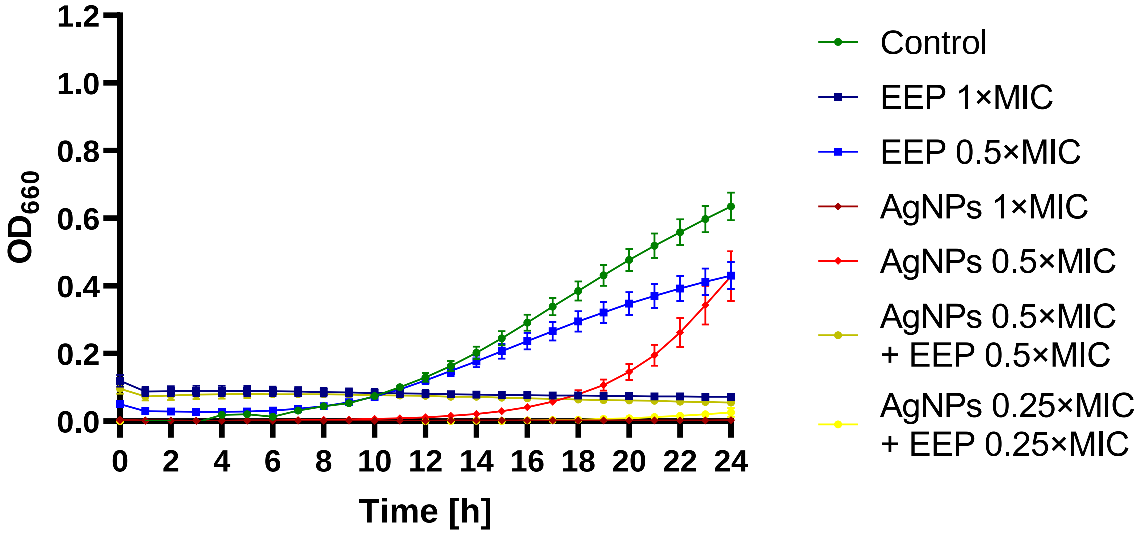
1.2
1.0
0.8
0.6
0.4
0.2
0.0
$OD_{660}$
0
2
4
6
8
10
12
14
16
18
20
22
24
Time [h]
Control
EEP 1×MIC
EEP 0.5×MIC
AgNPs 1×MIC
AgNPs 0.5×MIC
AgNPs 0.5×MIC + EEP 0.5×MIC
AgNPs 0.25×MIC + EEP 0.25×MIC

B

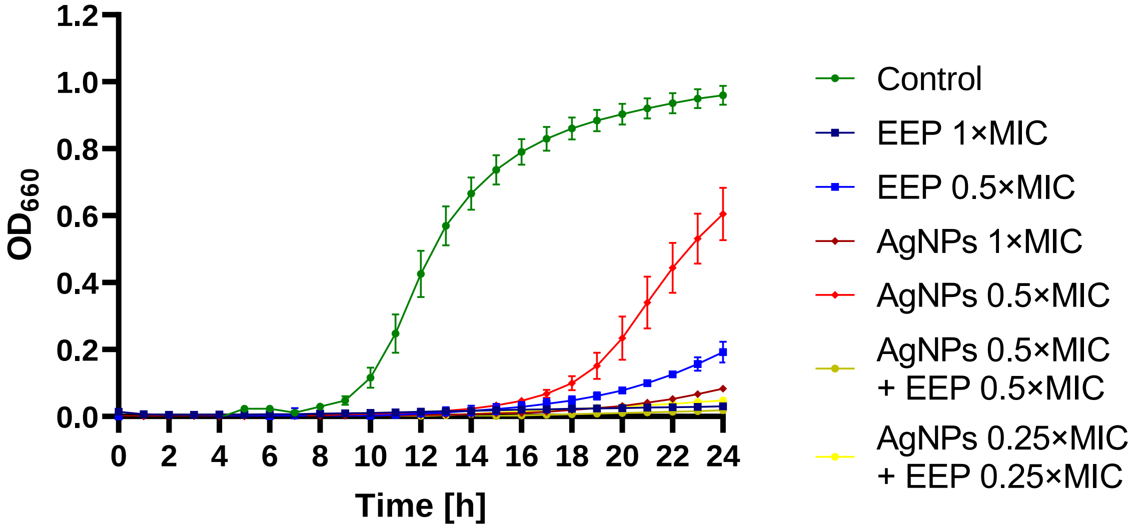
1.2
1.0
0.8
0.6
0.4
0.2
0.0
$OD_{660}$
0
2
4
6
8
10
12
14
16
18
20
22
24
Time [h]
Control
EEP 1×MIC
EEP 0.5×MIC
AgNPs 1×MIC
AgNPs 0.5×MIC
AgNPs 0.5×MIC + EEP 0.5×MIC
AgNPs 0.25×MIC + EEP 0.25×MIC

C

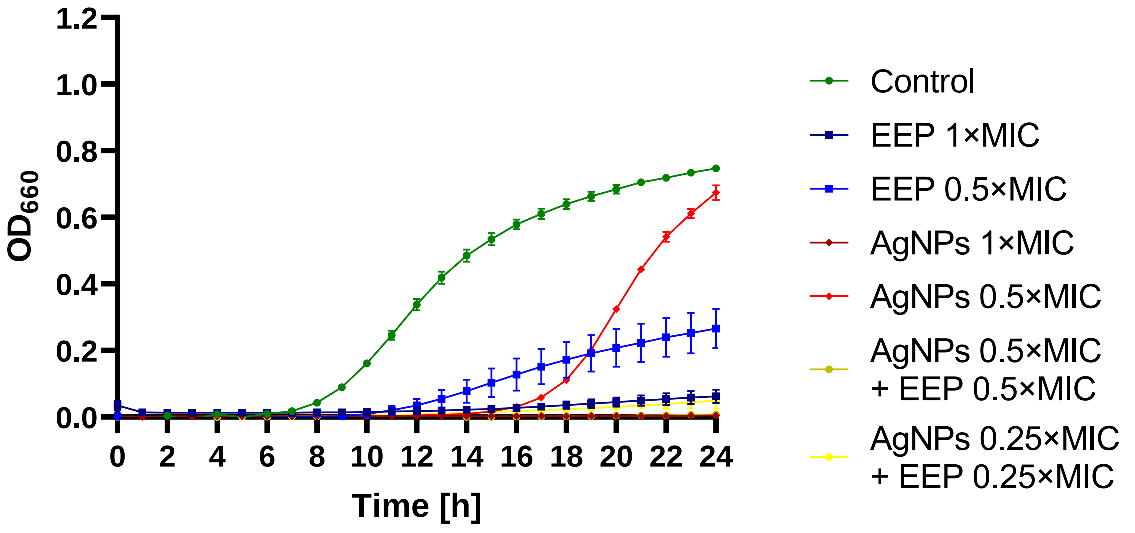


D

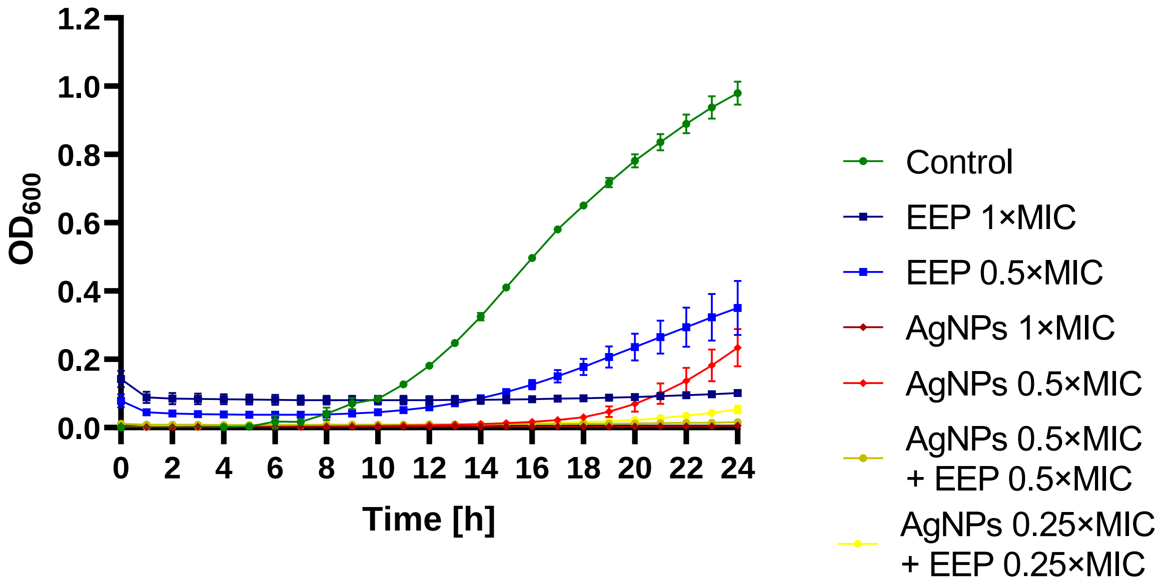


Figure S30 – Antifungal activity of EEP, silver nanoparticles (AgNPs), and their mixtures – growth curves (monitoring of the growth kinetics of the yeast cells measured as turbidity of the cultures at wavelength 660 nm or 600 nm for *C. auris*) against A – *C. albicans*, B – *N. glabratus*, C – *P. kudriavzevii*, and D – *C. auris*.

Table S20 – Concentrations of antifungals used in disc-diffusion assay.

| Antifungal | Form | Concentration used |
|---|---|---|
| Amfotericin B | Ready-to-use disc | 20 µg |
| Nystatin | Ready-to-use disc | 100 U |
| Caspofungin | Ready-to-use disc | 5 µg |
| Fluconazole | Ready-to-use disc | 100 µg |
| Voriconazole | Ready-to-use disc | 1 µg |
| Clotrimazole | Ready-to-use disc | 50 µg |
| Ketoconazole | Ready-to-use disc | 10 µg |
| 5-Fluorocytosine | Solution | 0,5 mg/ml |
| Chlorhexidine | Solution | 2 mg/ml |
| 2-Phenoxyethanol | Solution | 100% |
| Silver nanoparticles | Solution | 3 mg/ml |
| Hydrogen Peroxide | Solution | 1 % |
| Cyclopirox | Solution | 2 mg/ml |

Table S21 – Inhibition zones [mm] measured in disc-diffusion assay.

| | *C. albicans* | | | | | | *N. glabratus* | | | | | |
|---|---|---|---|---|---|---|---|---|---|---|---|---|
| | EtOH control | | | With EEP | | | EtOH control | | | With EEP | | |
| Amphotericin B | 20 | 16 | 18 | 24 | 24 | 25 | 18 | 17.5 | 18 | 22 | 21 | 21 |
| Nystatin | 24 | 24 | 24 | 28 | 28 | 28 | 24 | 24 | 24 | 29 | 29 | 29 |
| Caspofungin | 20 | 20 | 20 | 18 | 18 | 19 | 22 | 22 | 22 | 21 | 21 | 20 |
| Fluconazole | 9 | 10 | 10 | 20 | 20 | 19 | 28 | 28 | 28 | 32 | 28 | 33 |
| Voriconazole | 9 | 9 | 9 | 23 | 23 | 23 | 28 | 28 | 28 | 30 | 26 | 29 |
| Clotrimazole | 28 | 28 | 28 | 31 | 31 | 32 | 29 | 30 | 30 | 38 | 39 | 37 |
| Ketoconazole | 17 | 17 | 17 | 22 | 22 | 22 | 29 | 29 | 29 | 22 | 21 | 25 |
| 5-Fluorocytosine | 30 | 31 | 31 | 36 | 37 | 36 | 42 | 40 | 41 | 47 | 47 | 48 |
| Chlorhexidine | 13 | 13.5 | 18 | 15 | 14 | 16 | 13 | 12 | 13 | 14 | 12 | 12 |
| 2-Phenoxyethanol | 26 | 23 | 25 | 28 | 32 | 31 | 26 | 25 | 26 | 28 | 28 | 29 |
| AgNPs | 10 | 9 | 9 | 10 | 11 | 11 | 18 | 19 | 19 | 10 | 12 | 11 |
| Hydrogen Peroxide | 10 | 11 | 10 | 13 | 11 | 11 | 9 | 10 | 9.5 | 12 | 10 | 11 |
| Cyclopirox | 15.5 | 13 | 16 | 15 | 12.5 | 15 | 12 | 12 | 14.5 | 12.5 | 11.5 | 14 |

| | *P. kudriavzevii* | | | | | | *C. auris* | | | | | |
|---|---|---|---|---|---|---|---|---|---|---|---|---|
| | EtOH control | | | With EEP | | | EtOH control | | | With EEP | | |
| Amphotericin B | 17 | 17 | 17 | 19 | 19 | 18 | 17.5 | 17 | 17 | 19 | 18.5 | 19 |
| Nystatin | 22 | 22 | 22 | 25 | 25 | 24 | 19 | 19 | 18 | 20.5 | 19 | 18.5 |
| Caspofungin | 14 | 14 | 14 | 16 | 16 | 16 | 21 | 20.5 | 21 | 13 | 14 | 13 |
| Fluconazole | 19 | 21 | 20 | 18 | 18 | 17 | 10 | 10 | 10 | 12.5 | 12 | 12 |
| Voriconazole | 27 | 27 | 27 | 27 | 29 | 27 | 10 | 10 | 10 | 13 | 13 | 12 |
| Clotrimazole | 40 | 40 | 40 | 40 | 41 | 40 | 43 | 43 | 47 | 29 | 29 | 33 |
| Ketoconazole | 25 | 25 | 25 | 21 | 18 | 19 | 28 | 29 | 28 | 23 | 24 | 25 |
| 5-Fluorocytosine | 11 | 11 | 11 | 14 | 15 | 14 | 34 | 32 | 31 | 34 | 32 | 31 |
| Chlorhexidine | 15.5 | 16 | 14 | 13 | 13 | 13 | 10 | 9.5 | 10 | 8 | 11 | 8 |
| 2-Phenoxyethanol | 24 | 26 | 25 | 26 | 27 | 27 | 26 | 26 | 28 | 26 | 27 | 28 |
| AgNPs | 10 | 14 | 12 | 14 | 13 | 13 | 10 | 10 | 8 | 14.5 | 15.5 | 16 |
| Hydrogen Peroxide | 12 | 13 | 12 | 15 | 15 | 14 | 9 | 8 | 9 | 10.5 | 9.5 | 9.5 |
| Cyclopirox | 16 | 12 | 14 | 15.5 | 11 | 13.5 | 16 | 14.5 | 15.5 | 14.5 | 13.5 | 14.5 |

Table S22 – Mean growth reduction in $log_{10}$ CFU/ml values for each treatment at the 5th day in the simulated infection model with standard errors.

| | *C. albicans* | | *N. glabratus* | | *P. kudriavzevii* | | *C. auris* | |
|---|---|---|---|---|---|---|---|---|
| | $log_{10}$ CFU/ml | SE | $log_{10}$ CFU/ml | SE | $log_{10}$ CFU/ml | SE | $log_{10}$ CFU/ml | SE |
| EEP | 0.34 | 0.14 | 1.83 | 0.05 | 0.87 | 0.06 | 0.78 | 0.09 |
| Amphotericin B | 0.03 | 0.06 | 0.71 | 0.04 | 1.20 | 0.56 | 0.56 | 0.05 |
| Fluconazole | 1.00 | 0.26 | 0.19 | 0.03 | 1.74 | 0.24 | 0.61 | 0.07 |
| Ketoconazole | 2.26 | 0.02 | 1.46 | 0.35 | 4.73 | 0.11 | 0.94 | 0.05 |
| Clotrimazole | 0.86 | 0.26 | 4.10 | 0.12 | 3.17 | 1.08 | 0.31 | 0.28 |
| EEP + AMB | 4.88 | 0.04 | 3.09 | 0.12 | 3.61 | 0.10 | 1.64 | 0.10 |
| EEP + FLU | 4.88 | 0.04 | 2.61 | 0.14 | 0.84 | 0.22 | 0.92 | 0.11 |
| EEP + KET | 1.32 | 0.12 | 2.94 | 0.07 | 0.89 | 0.04 | 0.83 | 0.07 |
| EEP + CLO | 0.34 | 0.07 | 2.85 | 0.05 | 0.50 | 0.11 | 0.93 | 0.08 |

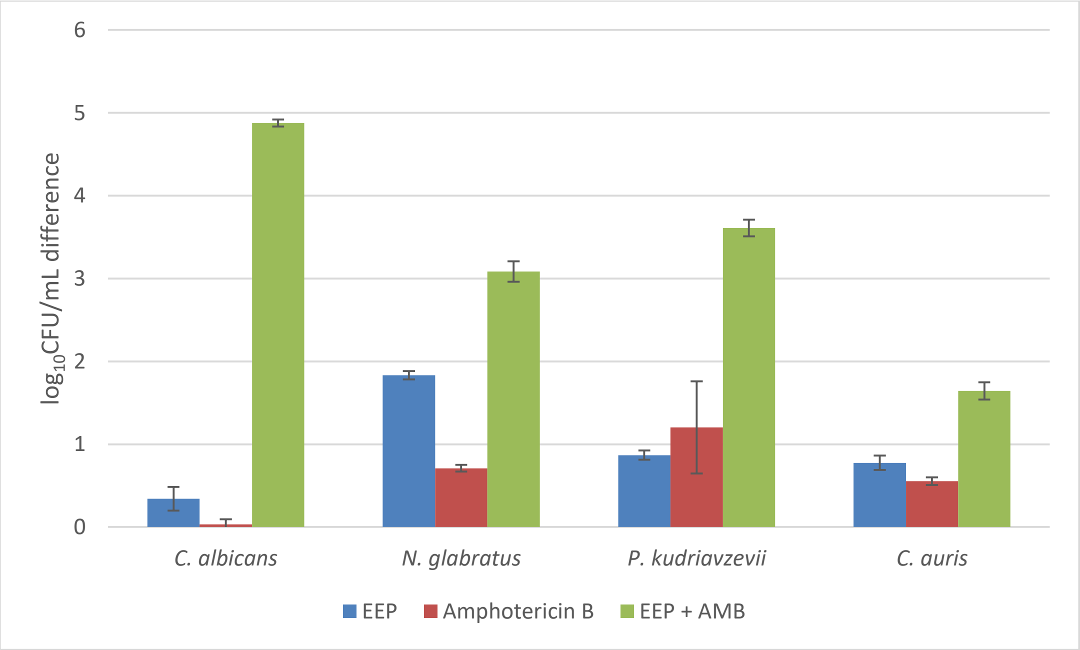


Figure S31 - Growth inhibition in comparison to control in simulated infection model for amphotericin B with EEP.

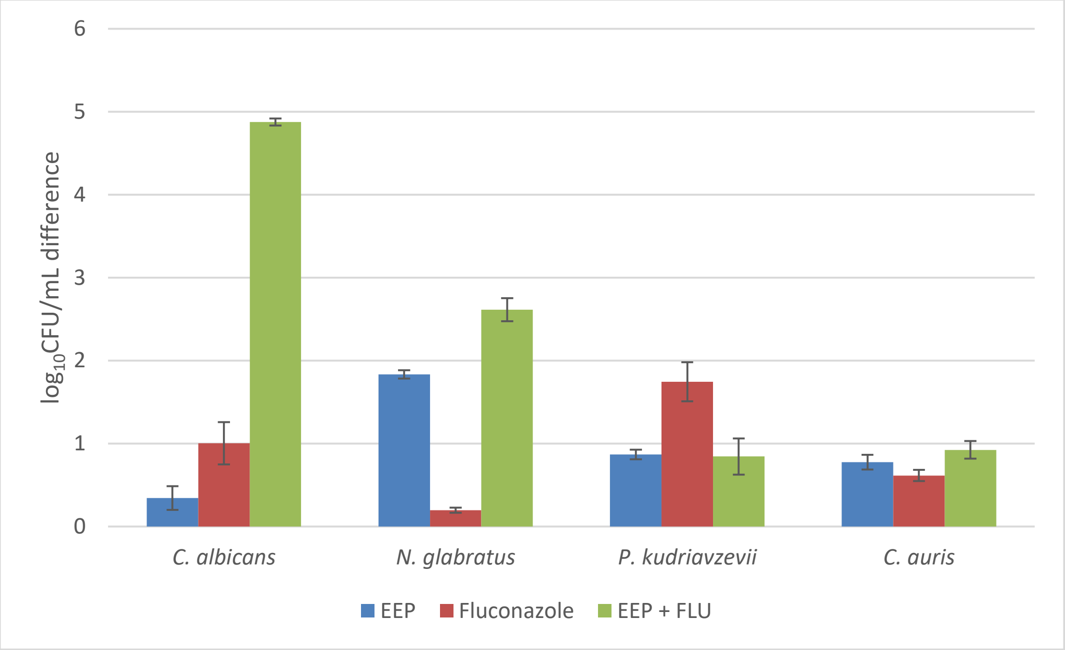


Figure S32 - Growth inhibition in comparison to control in simulated infection model for fluconazole with EEP.

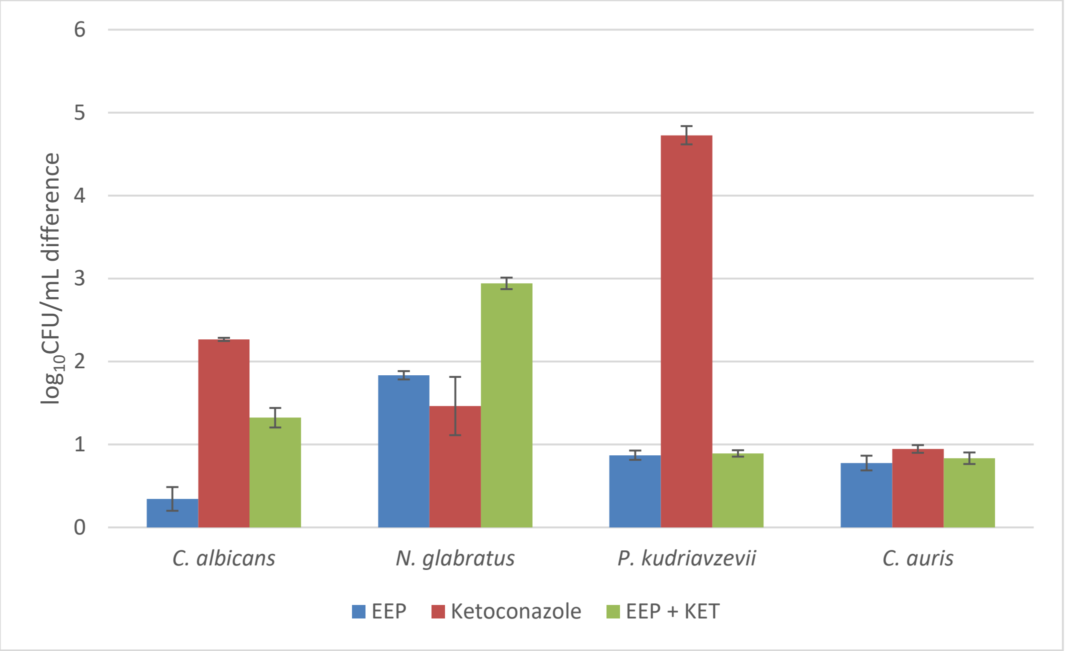


Figure S33 - Growth inhibition in comparison to control in simulated infection model for ketoconazole with EEP.

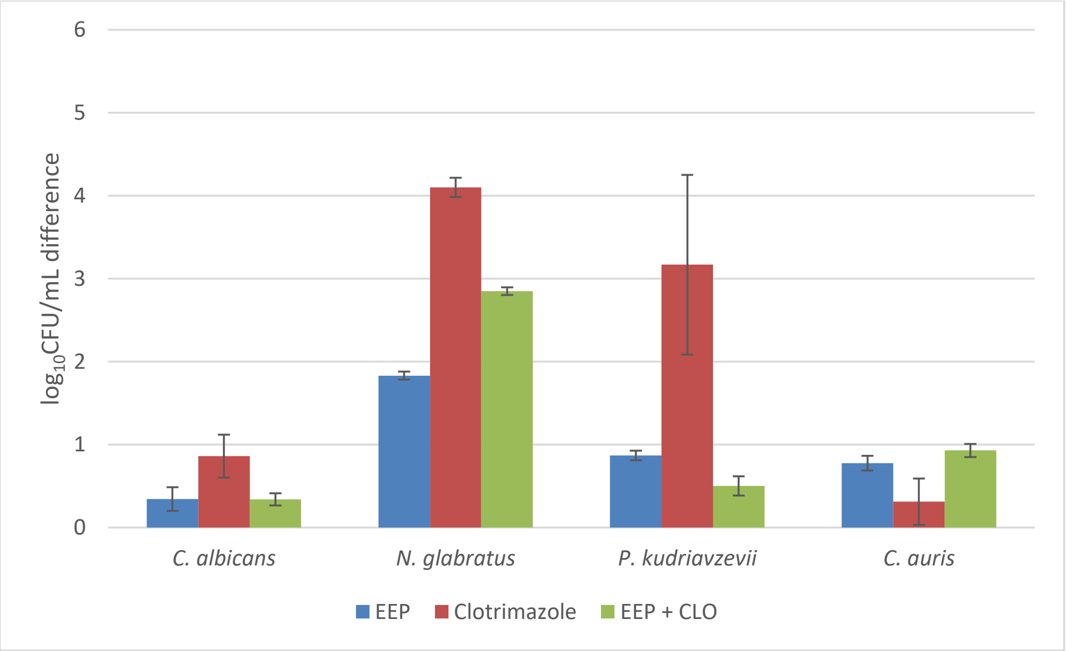


Figure S34 - Growth inhibition in comparison to control in simulated infection model for clotrimazole with EEP.
